# Genetic and pharmacological inactivation of peptidoglycan remodeling increases antibiotic susceptibility of vancomycin-resistant *Enterococcus faecium*

**DOI:** 10.1101/2025.09.11.675460

**Authors:** Kyong T. Fam, Pavan Kumar Chodisetti, Zifei Wang, Joshua A. Homer, Christopher J. Smedley, Seiya Kitamura, Benjamin Silva, Yijun Xiong, Althea Hansel-Harris, Matthew Holcomb, Simeon Babarinde, Adrianna M. Turner, Daria Van Tyne, Ian A. Wilson, Stefano Forli, Benjamin F. Cravatt, Donghyun Park, Dennis W. Wolan, John E. Moses, Howard C. Hang

## Abstract

Vancomycin-resistant *Enterococcus faecium* (VREfm) is a leading cause of healthcare-associated infections globally and demands new approaches for treatment. Here we show that genetic and pharmacological inactivation of a highly conserved NlpC/P60 peptidoglycan hydrolase, secreted antigen A (SagA), enhanced vancomycin susceptibility of VREfm *ex vivo* and *in vivo*. Notably, genetic deletion of *sagA* impaired VREfm peptidoglycan remodeling, growth and increased the activity of vancomycin. We then identified first-in-class covalent NlpC/P60 peptidoglycan hydrolase inhibitors and demonstrated that pharmacological inactivation of SagA activity also impaired peptidoglycan remodeling and increased the efficacy of vancomycin across genetically distinct VREfm clinical isolates. Our study reveals peptidoglycan hydrolases are druggable targets whose inactivation improves the efficacy of vancomycin against VREfm.

## Introduction

Increasing antimicrobial resistance (AMR) in bacterial pathogens and limited antibiotic discovery require new approaches to address this major threat to human health world-wide.^1^ Amongst the ESKAPE pathogens,^2^ vancomycin-resistant *E. faecium* (VREfm) infections have become more prevalent among healthcare-associated infections and can acquire resistance to last-resort antibiotics like linezolid, daptomycin, and tigecycline.^3,4^ The AMR crisis demands new approaches to prevent and treat VREfm infections, which are also correlated with poor patient outcomes leading to high mortality rates.^5,6^ While the synthesis of next-generation antibiotics provides new derivatives to address AMR^7,8^ and innovative approaches are being employed to discover new classes of antibiotics,^1,9,10^ a better understanding of *E. faecium* biology may provide new targets for antimicrobial development.

*Enterococcus* is a genus of ubiquitous Gram-positive bacteria, among which *E. faecium* and *E. faecalis* are the most prominent in humans and other mammals.^11^ While *E. faecium* can acquire antibiotic resistance and cause healthcare-associated infections,^2,11,12^ non-pathogenic strains of *E. faecium* have been reported to have beneficial effects on host physiology and been developed into probiotics.^13^ Our mechanistic dissection of commensal *E. faecium*-host interactions revealed that secreted antigen A (SagA), a highly conserved NlpC/P60 peptidoglycan hydrolase, can generate non-crosslinked muropeptides to promote host immunity.^14–19^ Of note, we also demonstrated *sagA* is essential for peptidoglycan remodeling, cell separation and growth in the commensal strain of *E. faecium* (Com15).^19^ Moreover, the commensal strains of *E. faecium* (Com15-Δ*sagA* and phage-resistant Com12 expressing catalytically inactive alleles of *sagA*) were more susceptible to cell-wall targeting antibiotics,^19–21^ suggesting SagA may be a potential antimicrobial target.

As *sagA* is highly conserved amongst *E. faecium* strains, including VREfm strains containing the vancomycin resistance *vanA* and *vanB* gene clusters,^17^ we investigated genetic and pharmacological inactivation of this key peptidoglycan hydrolase as a therapeutic target for treating VREfm infections. We discovered that VREfm-Δ*sagA* showed defective peptidoglycan remodeling, impaired cell separation and growth as well as increased susceptibility to last-line antibiotics. Notably, even though a VREfm-Δ*sagA* strain still encodes the *vanA* operon, the minimal inhibitory concentration (MIC) of vancomycin was decreased and attenuated by vancomycin treatment in a mouse model of VREfm-induced sepsis *in vivo*.

In parallel, we explored pharmacological inactivation of SagA activity. Bacterial essential enzymes and virulence factors can be attenuated by covalent inhibitors, including recent fluorosulfates and sulfonyl fluorides that can undergo Sulfur(VI) Fluoride Exchange (SuFEx) chemistry.^22,23^ Sulfonyl fluorides can covalently label a broad range of amino acid residues (Ser > Thr > Tyr > Cys > Lys > His), however their reactivity is context-dependent and largely governed by catalytic environment of the target enzyme.^22–24^ From a library of promiscuous SuFEx-based sulfonyl fluorides accessed through a Diversity Oriented Clicking (DOC) approach^24–26^, we identified β-chloro alkenyl sulfonyl fluorides as the first-in-class covalent inhibitors of the NlpC/P60 cysteine endopeptidases. The most potent β-chloro alkenyl sulfonyl fluoride SagA inhibitor impaired peptidoglycan remodeling in VREfm, reduced the MIC of vancomycin in genetically distinct VREfm strains, attenuated infection of macrophages *ex vivo,* and improved outcomes of VREfm-induced sepsis *in vivo*. Our studies demonstrate that peptidoglycan hydrolases are crucial for bacterial cell wall remodeling and are druggable targets to promote the efficacy of antibiotics against VREfm.

## Results

### SagA is critical for peptidoglycan remodeling and antibiotic susceptibility of VREfm

SagA is a member of the highly conserved NlpC/P60-family of cysteine endopeptidases that are important for bacterial physiology.^27^ We previously demonstrated SagA is crucial for commensal *E. faecium* (Com15 strain) peptidoglycan remodeling, cell separation and growth.^19^ Consistent with our previous analysis,^17^ phylogenetic analysis confirmed the presence of *sagA* using an international collection of publicly available vancomycin-susceptible *E. faecium* (n=164) and VREfm (n=395) isolates from 99 sequence types (Extended Data Fig.1). Our analysis also revealed the distribution of other NlpC/P60 hydrolases in *E. faecium*, with some being more prevalent in VREfm strains (Extended Data Fig.1), but their function was unknown. Only 4 out of 10 NlpC/P60 hydrolases were found in human clinical VREfm isolate ERV165 (sequence type 412; *vanA* genotype; Extended Data Fig. 2a). Using improved genetic methods for *Enterococcus*,^28,29^ we generated isogenic deletion strains of all 4 NlpC/P60 hydrolases in VREfm ERV165 strain (Extended Data Fig. 2b-e) and found that only the ERV165-Δ*sagA* (further referred as Δ*sagA*) mutant exhibited impaired growth and altered colony morphology (Fig. 1a-c and Extended Data Fig. 2f), which was rescued by *sagA* chromosomal complementation (Δ*sagA*::*sagA*). Transmission electron microscopy (TEM) analysis revealed that Δ*sagA* cells failed to properly separate, resulting in the formation of aberrant cellular clusters (Fig. 1d). Further cryo-electron tomography (cryo-ET) analysis of Δ*sagA* showed a decrease in cell wall thickness and a modest increase in septum thickness (Extended Data Fig. 3 and 4). The cell wall thickness was restored in the *sagA*-complemented strain, however no significant changes in septum thickness were observed (Extended Data Fig. 4c). Similar to Com15-Δ*sagA* mutant^19^, Δ*sagA* also exhibited lower amounts of non-crosslinked muropeptides and increased levels of crosslinked peptidoglycan fragments (Extended Data Fig. 5). These results demonstrate that in addition to its importance in commensal *E. faecium* strains, *sagA* is also essential for VREfm peptidoglycan remodeling, bacterial cell separation and growth, even though the majority of VREfm strains have acquired additional NlpC/P60 hydrolases (Extended Data Fig. 1).

**Fig. 1.**
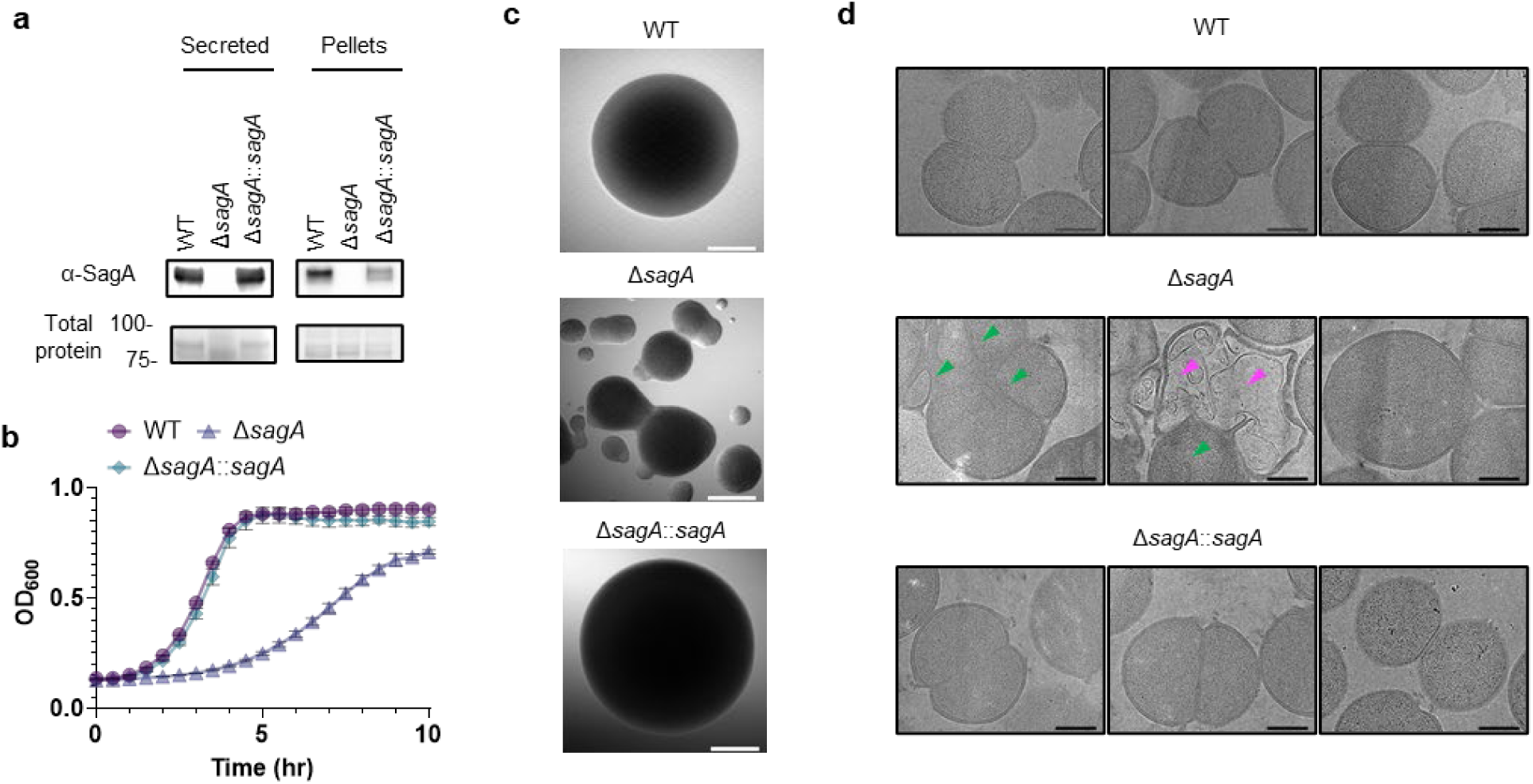
SagA is important for peptidoglycan remodeling, cell separation and growth in VREfm. **a**, α-SagA western blot analysis of bacterial proteins (secreted and in cell pellets) of VREfm ERV165 strains (WT, Δ*sagA* or Δ*sagA*::*sagA*). Stain-free imaging serves as total protein loading control. **b**, Growth curves of VREfm ERV165 strains in BHI. **c**, Representative DIC images of single colonies of VREfm strains. Scale bar, 200 µm. **d**, Representative low-magnification (3600×) electron microscopy images from lamellae of VREfm ERV165 strains. Dead cells are indicated with magenta arrows; defective cell divisions are marked with green arrows. Scale bars, 500 nm.

We next evaluated the contribution of SagA to VREfm antibiotic susceptibility. Based on previous antibiotic susceptibility studies in commensal *E. faecium* Δ*sagA* mutant strains,^19^ we analyzed the activity of ampicillin, daptomycin and ceftriaxone in VREfm Δ*sagA*. Surprisingly, VREfm Δ*sagA* only exhibited modest inhibition of bacterial growth with ampicillin, daptomycin and ceftriaxone compared to ERV165 wild-type (Extended Data Fig. 6a-c) and no significant differences in MIC (Extended Data Table 1). However, Δ*sagA*, but not other NlpC/P60 hydrolase deletion strains (Extended Data Fig. 6d), showed increased susceptibility to vancomycin with 2-fold difference of MIC (Extended Data Table 2), which was abrogated in the *sagA*-complemented strain (Fig. 2a,b and Extended Data Table 2). Adaptive laboratory evolution experiment in the presence of sub-MIC vancomycin concentration demonstrated that neither Δ*sagA*, nor WT or *sagA*-complemented strain developed additional vancomycin resistance after two weeks (Extended Data Fig 6e). To investigate the vancomycin susceptibility of ERV165-Δ*sagA*, we performed whole-genome sequencing (WGS), evaluated antibiotic binding and analyzed peptidoglycan remodeling further. WGS of the Δ*sagA* strain confirmed the *sagA*-gene deletion and revealed 7 missense, 1 frame-shift and 12 silent mutations (Extended Data Table 3 and 4), none of which were associated with peptidoglycan synthesis, cell wall remodeling or vancomycin resistance (Extended Data Fig. 7a). Of note, both Δ*sagA* and *sagA*-complemented strains have L10S and L68S missense mutations found in *mapZ* (Midcell Anchored Protein Z) homolog that encodes a protein essential for bacterial cell division (Extended Data Table 3 and 5). However, these mutations occurred outside annotated functional domains and in a region lacking conserved sequence features. Importantly, the increased vancomycin susceptibility of Δ*sagA* was abrogated in the *sagA*-complemented strain that retained the mutations as Δ*sagA* (Fig. 2b and Extended Data Table 2, 5 and 6). We then employed fluorescent vancomycin (Van-BODIPY)^30^ and fluorescent D-amino acid (HADA)^31^ to evaluate antibiotic binding and peptidoglycan stem peptide remodeling, respectively. Fluorescence microscopy revealed increased Van-BODIPY and HADA staining in Δ*sagA* that was abrogated by *sagA* complementation (Fig. 2c-e). These results suggest that loss of SagA expression and defective peptidoglycan remodeling led to the improved vancomycin binding and increased antibiotic susceptibility.

**Fig. 2.**
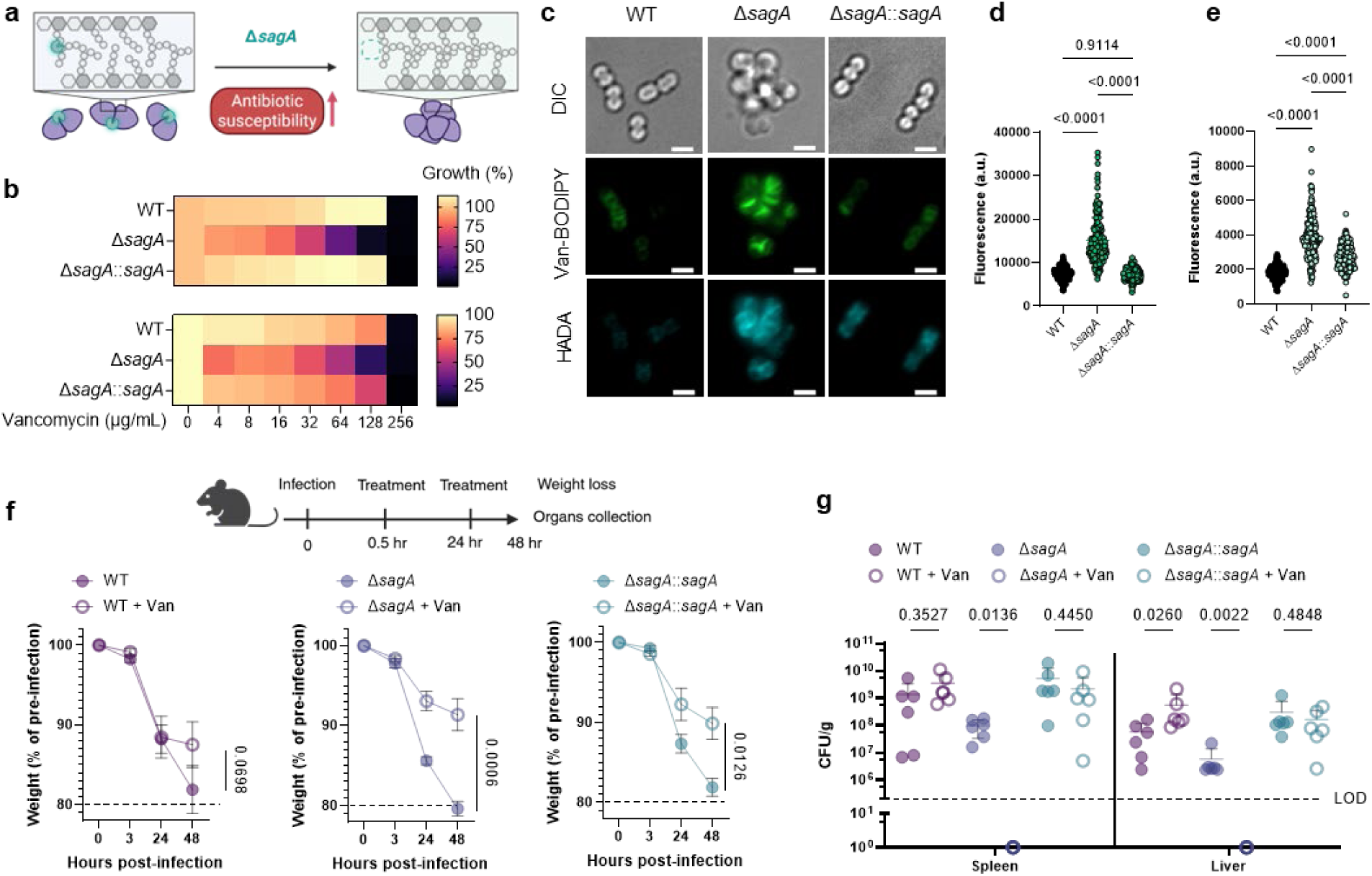
SagA is important for VREfm antibiotic susceptibility *ex vivo* and *in vivo*. **a**, Scheme illustrating inactivation of peptidoglycan remodeling upon genetic deletion of *sagA* in VREfm. **b**, Vancomycin susceptibility of ERV165 strains after 10 hours of growth. Top: susceptibility in BHI. Bottom: susceptibility in MHB. Data is a heat map of mean values, n=3 biological replicates. **c**, Fluorescence microscopy imaging of VREfm strains stained with Van-BODIPY (1 µg/mL) and fluorescent D-amino acids (HADA, 0.5 mM). Differential Interference Contrast (DIC) images show cell wall morphology, with cell clustering apparent in Δ*sagA*. Scale bar, 2 µm. **d**, Fluorescence of Van-BODIPY-stained VREfm strains. **e**, Fluorescence of HADA-stained VREfm strains. For (**d**), (**e**) each dot is an individual cell, n=200. **f**, Top: scheme of systemic infection *in vivo* with VREfm strains (ERV165 WT, Δ*sagA* or Δ*sagA*::*sagA*): mice were infected with 10^9^ VREfm CFU intraperitoneally. PBS or vancomycin (Van, 100 mg/kg) was administered subcutaneously. Weight was monitored at 3, 24, and 48 hours post-infection. Mice were sacrificed, organs were collected and VREfm burdens were analyzed by CFU counting. Bottom: weight (% of pre-infection) of mice infected with VREfm ERV165 strains (WT, Δ*sagA* or Δ*sagA*::*sagA*) ± vancomycin (Van, 100 mg/kg). Data is mean ± S.E.M. and analyzed by one-way ANOVA with uncorrected Fisher’s LSD post-test, n=6 mice. **g**, CFU analysis of organs from mice infected with VREfm strains (ERV165 WT, Δ*sagA* or Δ*sagA*::*sagA*) ± vancomycin (Van, 100 mg/kg). Horizontal line is mean ± S.E.M. Data is analyzed by unpaired Mann-Whitney test. Each dot is an individual mouse, n=6.

To evaluate the impact of SagA on VREfm infection *in vivo*, we employed a mouse peritonitis infection model^32^. All three strains (ERV165 wild-type, Δ*sagA* and *sagA*-complemented strain) caused similar levels of weight loss as a marker of infection and exhibited high bacterial burdens in the spleen and liver (Fig. 2f,g). However, treatment with clinical doses of vancomycin^33^ significantly improved weight loss and cleared bacterial burden in mice infected with Δ*sagA* strain, but not in those infected with the wild-type VREfm or *sagA*-complemented strains (Fig. 2f,g). These results demonstrate that SagA not only impacts peptidoglycan remodeling and activity of cell wall-targeting antibiotics in commensal *E. faecium* strains, but importantly also modulates vancomycin susceptibility in VREfm *ex vivo* and VREfm-induced sepsis *in vivo*.

### SuFEx-based sulfonyl fluorides covalently label and inactivate SagA NlpC/P60 hydrolase activity

To identify pharmacological inhibitors of SagA endopeptidase activity, we developed a high-throughput assay based on competitive labeling^34^ of the only cysteine residue (C433) in the NlpC/P60 hydrolase domain active site^35^ using a fluorescent tetramethyl rhodamine-iodoacetamide (TMR-IA) probe (Fig. 3a and Extended Data Fig. 8a). TMR-IA selectively labeled the recombinant SagA-NlpC/P60 hydrolase domain, but not the inactive C433A mutant or wild-type pretreated with cysteine-reactive controls N-methylmaleimide (NMM) or iodoacetamide (IA) (Extended Data Fig. 8b-d). Using this assay, we screened a library of SuFEx-based sulfonyl fluorides^24^ for potential covalent inhibitors of SagA. We identified 86 sulfonyl fluorides that reduced TMR-IA labeling greater than 80% (Extended Data Fig. 8e). Secondary screening of the top sulfonyl fluorides using gel-based competitive TMR-IA labeling (Extended Data Fig. 8f-i) and follow-up SagA peptidoglycan hydrolase activity assays^36^ (Extended Data Fig. 8a), revealed a subset of β-chloro alkenyl sulfonyl fluoride compounds (peptidoglycan hydrolase inhibitors, pghi-1 to 5) with low micromolar IC_50_ values *in vitro* (Fig. 3b,c and Extended Data Fig. 9b). Small modifications in the phenyl ring (pghi-1 to 5) did not substantially affect SagA inhibition *in vitro*, whereas substitution of the β-alkenyl position (pghi-6) compromised inhibition potency (Fig. 3b,c and Extended Data Fig. 9b).

**Fig. 3.**
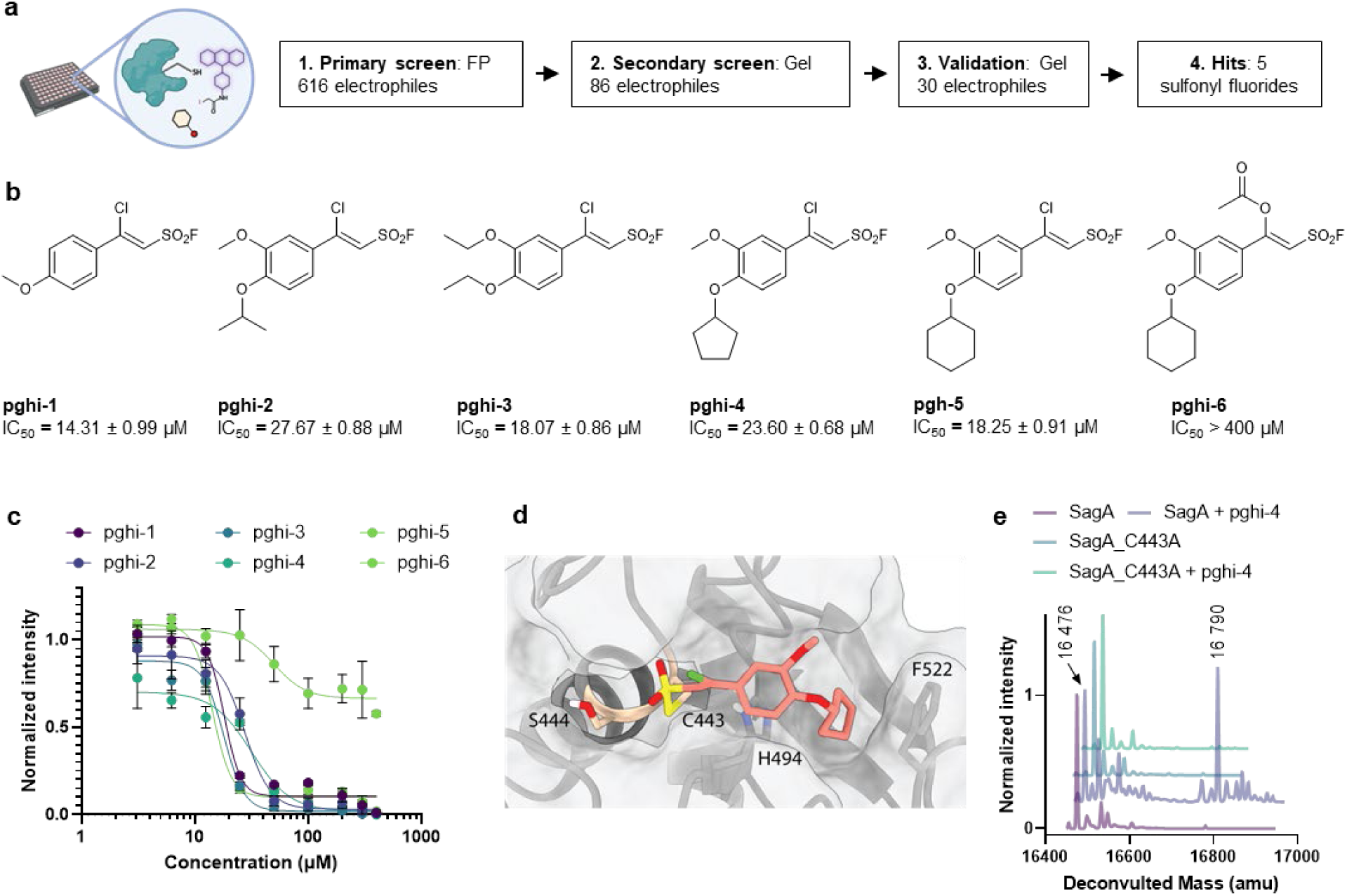
Activity-guided screening identifies sulfonyl fluorides as SagA inhibitors. **a**, Scheme and workflow of activity-guided screening. An agnostic screen of 616 electrophiles in competitive fluorescence polarization (FP) assay led to selection of 86 electrophiles that were screened in a competitive gel-based cysteine activity assay. Activities of 30 molecules were further validated in a gel-based peptidoglycan hydrolase activity assay (Extended Data Fig. 7). **b**, Structures of identified 5 most active (pghi-1 to 5) and inactive (pghi-6) sulfonyl fluorides as SagA inhibitors with biochemical IC_50_ for SagA activity (Extended Data Fig. 8a,b). **c**, IC_50_ curves of sulfonyl fluorides determined by gel-based peptidoglycan hydrolase activity assay. Data is mean ± S.E.M., n=3 biological replicates. **d**, Covalent docking model of pghi-4 bound to C433 of SagA active site as the product of reaction *via* sulfonyl fluoride. **e**, Intact protein mass analysis of SagA (10 µM) ± pghi-4 (200 µM) and inactive mutant SagA_C433A (10 µM) ± pghi-4 (200 µM). Numbers indicate detected masses of SagA and SagA-pghi-4 adduct.

Computational covalent docking suggested that these peptidoglycan hydrolase inhibitors can be accommodated in the active site, stabilized by interactions with nearby H494 and F522, after reacting with cysteine (C443) of SagA via the sulfonyl fluoride group (Fig. 3d) or β-alkenyl chloride group (Extended Data Fig. 10a). Intact protein mass spectrometry analysis of recombinant SagA treated with pghi-4 revealed a mass shift that corresponds to the mass of the SagA-pghi-4 adduct (Fig. 3e). The observed mass increase matched the calculated molecular weight loss of fluorine from pghi-4, suggesting covalent modification through the sulfonyl fluoride group and formation of a thiosulfonate adduct (Extended Data Fig. 10b-d). Indeed, treatment of the catalytically inactive C433A SagA mutant with pghi-4 did not result in a mass shift (Fig. 3e). Incubation of SagA with an inactive pghi-6 resulted in minor formation of thiosulfonate-linked SagA-pghi-6 adduct (Extended Data Fig. 10e) indicating the importance of the β-vinyl chloro group for binding to SagA. Our efforts to crystallize SagA-pghi-4 complex did not yield diffraction quality crystals (see Methods), likely due to instability of the SagA-pghi-4 adduct. Further intact protein mass spectrometry analysis of the SagA-pghi-4 adducts (peak 1, thiosulfonate, Extended Data Fig. 9b-d) identified peaks 3 and 5 with masses corresponding to thiosulfonic and sulfinic acid derivatives of SagA (Extended Data Fig. 10b-d). After 16 hours of treatment, the abundance of the SagA-pghi-4 adduct significantly decreased, while the levels of hydrolysis products increased (Extended Data Fig. 10b-d), indicating instability of the SagA-pghi-4 adduct. To explore the selectivity of identified pghi-1 to 5, we evaluated these compounds with other VREfm NlpC/P60 hydrolases *in vitro*. Pghi-1 to 5 also inhibited the endopeptidase activity of peptidoglycan hydrolase 2 (PGH2), which has NlpC/P60 domain with high amino acid sequence and structural homology to SagA, but not the more divergent CwlT-like peptidoglycan hydrolase 3 (PGH3) (Extended Data Fig. 11a-c). Taken together, these results suggest that the identified β-chloro alkenyl sulfonyl fluorides can bind and covalently react with the catalytic cysteine of SagA and peptidoglycan hydrolase orthologs with structurally similar NlpC/P60 hydrolase domains to inactivate their enzymatic activity.

### SagA inhibitors improve antibiotic susceptibility in VREfm strains

We next evaluated the activity of pghi-1 to 6 on VREfm growth and antibiotic susceptibility. Either pghi-4 or pghi-5 alone caused a mild growth delay of VREfm (ERV165), while the other compounds did not significantly affect VREfm growth under laboratory conditions (Extended Data Fig. 12a,b). Building upon our observations of SagA modulation of VREfm antibiotic susceptibility (Figs. 1 and 2), we evaluated these compounds on VREfm growth in combination with a low dose of vancomycin (Fig. 4a and Extended Data Fig. 12c). Notably, pghi-4 and pghi-5 significantly enhanced the activity of vancomycin compared to the other β-chloro alkenyl sulfonyl fluorides and the inactive analog pghi-6 (Fig. 4a,b and Extended Data Fig. 12c). The most active compound pghi-4 lowered the MIC values of vancomycin up to 8 fold in a concentration-dependent manner by checkerboard assay analysis (Fig. 4c and Extended Data Table 7). Bacterial time-kill analysis showed that pghi-4 in combination with vancomycin significantly limited VREfm growth compared to either agent alone (Fig. 4d and Extended Data Fig. 12d). Similar to Δ*sagA*, pghi-4 only showed very modest to no enhancement of ampicillin, daptomycin and ceftriaxone antibiotic activity in VREfm (Extended Data Fig. 12e,f).

**Fig. 4.**
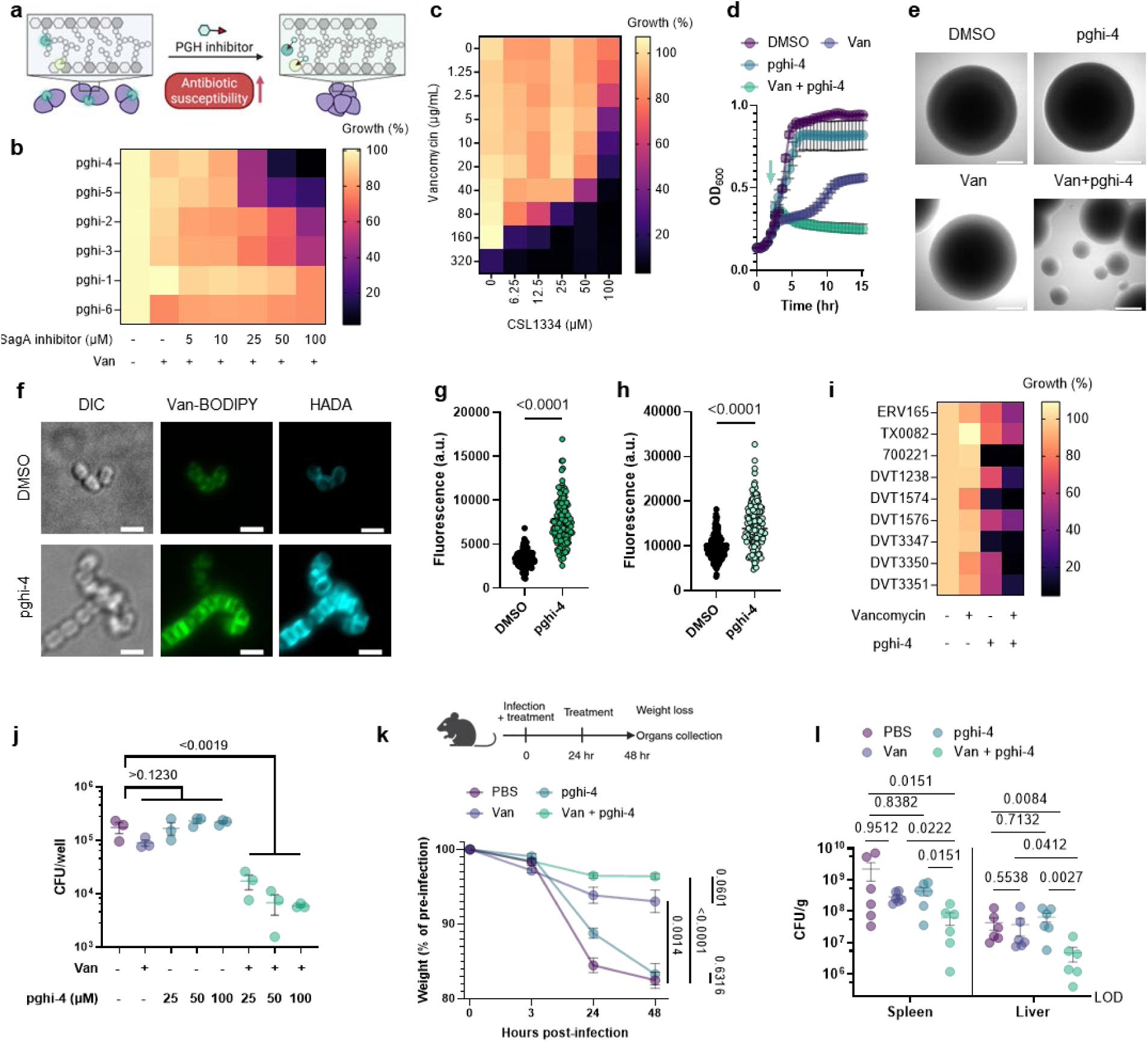
Pghi-4 is an antibiotic adjuvant against VREfm. **a,** Scheme illustrating inactivation of peptidoglycan remodeling upon pharmacological inhibition of SagA in VREfm. **b**, Effect of sulfonyl fluorides + vancomycin (Van, 5 μg/mL) on VREfm (ERV165) growth. **c**, Pghi-4 synergism with vancomycin against ERV165 evaluated by checkerboard assay. Data in **b** and **c** is heatmap of median values relative to DMSO control, n=3. **d**, Bacteria-kill growth curves. Exponentially grown ERV165 were treated with pghi-4 (100 μM), vancomycin (Van, 100 μg/mL) or in combination for 18 hours, followed by OD600 measurement and CFU analysis (Extended Data Fig. 11d). The green arrow indicates the time point when the treatment was added. **e**, DIC images of single colonies of VREfm ± vancomycin (Van, 100 μg/mL) ± pghi-4 (100 µM). Scale bar, 200 µm. **f**, Fluorescence microscopy imaging of VREfm ± pghi-4 (50 µM) stained with Van-BODIPY (50 µg/mL) and HADA (0.5 mM). Scale bar, 2 µm. **g**, Fluorescence of Van-BODIPY-stained VREfm ± pghi-4 (50 µM). **i**, Fluorescence of HADA-stained VREfm ± pghi-4 (50 µM). For **g**-**h** each dot is individual cell, n=143. **i**, Effect of pghi-4 (50 μM) ± vancomycin (Van, 5 μg/mL) on growth of VREfm clinical isolates. Data is heatmap of median values relative to DMSO control, n=3 biological replicates. **j**, CFU analysis of VREfm-infected RAW264.7 cells treated with vancomycin (100 µg/mL), pghi-4 alone or in combination. Data is mean and analyzed by one-way ANOVA with uncorrected Fisher’s LSD post-test ± S.E.M., n=3 biological replicates. **k**, Top: scheme of systemic infection *in vivo* with VREfm (ERV165): mice were infected with 10^9^ VREfm CFU with PBS (0.25% CMC), pghi-4 (25 mg/kg 0.25% CMC) ± vancomycin (Van, 100 mg/kg 0.25% CMC) intraperitoneally. 24 hours post-infection mice were given the second dose. Weight was monitored at 3, 24, 48 hours post-infection. Mice were sacrificed, organs were collected and VREfm burden was analyzed by CFU. Bottom: weight (% of pre-infection) of mice VREfm-infected ± 2 doses of pghi-4 (25 mg/kg 0.25% CMC) ± vancomycin (Van, 100 mg/kg 0.25% CMC). Data is mean ± S.E.M. and analyzed by one-way ANOVA with uncorrected Fisher’s LSD post-test, n=6 mice. **l**, CFU analysis of organs from VREfm-infected mice ± 2 doses of pghi-4 (25 mg/kg 0.25% CMC) ± vancomycin (Van, 100 mg/kg 0.25% CMC). Y axis starts from the limit of detection (LOD), horizontal line is mean ± S.E.M. Data is analyzed by Kruskal-Wallis test with Dunn’s uncorrected post-test. Each dot is individual mouse, n=6.

To characterize the mechanism of action of SagA inhibitor pghi-4, we performed imaging, peptidoglycan remodeling and chemoproteomic studies of VREfm. Differential interference contrast (DIC) microscopy revealed that pghi-4 in combination with vancomycin induced aberrant bacterial colony morphology and increased cell clustering compared to control or either agent alone (Fig. 4e,f). Cryo-ET showed dead cells with impaired peptidoglycan cleavage in VREfm treated with pghi-4 in combination with vancomycin (Extended Data Fig. 13). Both pghi-4 and vancomycin alone increased cell wall thickness, but their combination resulted in reduced thickness (Extended Data Fig. 14a,b). Interestingly, septum thickness was significantly increased under all treatment conditions (Extended Data Fig. 14a,c). These results suggest that the combination of pghi-4 and vancomycin impaired the ultrastructure of VREfm peptidoglycan. Similar to Δ*sagA*, pghi-4 also increased Van-BODIPY (5.9 fold, Fig. 4f,g) and HADA (Fig. 4f,h) labeling, suggesting enhanced vancomycin binding due to inhibition of peptidoglycan remodeling and the accumulation of vancomycin-target D-Ala-D-Ala-containing muropeptides^37^. Penicillin-BODIPY (Bocillin) labeling was only slightly increased in pghi-4 treated VREfm (1.3 fold) and in Δ*sagA* (2.3 folds, Extended Data Fig. 15a, b) by fluorescence microcopy as well as in-gel penicillin-binding protein 5 (PBP5) labeling (Extended Data Fig. 15c), which are consistent with our β-lactam (ampicillin) susceptibility analyses (Extended Data Fig. 6a-c, Extended Data Table 1 and Extended Data Fig. 12e,f). We observed mutations (M485A, A499T and E629V) in PBP5 of VREfm ERV165 compared to commensal *E. faecium* Com15 strain (Extended Data Fig. 15d), which are known to drive PBP5-dependent resistance to β-lactam antibiotics.^38^ Genetic deletion of *sagA* in VREfm ERV165 did not affect those mutations (Extended Data Fig. 15d), which may explain why β-lactam antibiotics are more active in commensal *E. faecium* Δ*sagA* strains^38^.

Quantitative LC-MS analysis of pghi-4-treated ERV165 did not yield similar profiles of soluble fragments from peptidoglycan compared to Δ*sagA* (Extended Data Fig. 16a,c). However, for these experiments ERV165 was only treated with sub-inhibitory and non-bactericidal dose of 50 µM pghi-4 to obtain sufficient material for LC-MS analysis. Moreover, the genetic inactivation of *sagA* may result in accumulation or depletion of soluble fragments from peptidoglycan that are not fully recapitulated with incomplete pharmacological inhibition of SagA. Nonetheless, LC-MS analysis of digested peptidoglycan fragments from pghi-4 and vancomycin co-treated ERV165 revealed higher levels of D-Ala-D-Ala-containing GlcNAc-MurNAc-pentapeptide in peptidoglycan (Extended Data Fig. 16d,e). As pghi-4 targets the active site C443 of SagA *in vitro* (Fig. 3d,e), we also evaluated covalent labeling of SagA and other reactive Cys-residues in ERV165 proteome by competitive chemoproteomics using an iodoacetamide-alkyne (IA-alk) probe (Extended Data Fig. 17a and Supporting Data 1). Indeed, treatment of ERV165 with pghi-4 reduced IA-alk labeling of the SagA by in-gel fluorescence labeling and western blot analysis (Extended Data Fig. 17b). The quantitative proteomic analysis of pghi-4-competitive Cys-reactive protein targets showed only a few other proteins were targeted in a dose-dependent manner (Extended Data Fig. 17c). However, none of these other Cys-reactive candidate pghi-4-target proteins have been implicated in VREfm peptidoglycan remodeling or antibiotic susceptibility. These results suggest that pharmacological inactivation of SagA by pghi-4 increases vancomycin susceptibility of VREfm by impairing peptidoglycan remodeling, which increases the levels of D-Ala-D-Ala-containing muropeptides and promotes vancomycin binding.

To characterize the scope of pghi-4 adjuvant activity with vancomycin, we evaluated additional VREfm strains including clinical isolates from patients that underwent chemotherapy or hematopoietic stem cell transplantation^39^. Notably, some VREfm strains (700221, DVT1574, DVT3347) showed increased susceptibility to pghi-4 and vancomycin co-treatment compared ERV165 and other VREfm strains (Fig. 4i and Extended Data Fig. 18a), even though they all harbor *vanA*-type resistance^39^, of different sequence types (Extended Data Table 8) and exhibit similar vancomycin MICs (Extended Data Table 9). Interestingly, pghi-4 adjuvant activity correlated with intracellular SagA protein expression levels in these VREfm strains (Extended Data Fig. 18b). These results suggest β-chloro-alkenyl sulfonyl fluoride SagA inhibitors can increase the vancomycin susceptibility of different VREfm clinical isolates, which correlates with their SagA protein expression levels.

### Pharmacological inactivation of SagA increases vancomycin susceptibility of VREfm *in vivo*

To investigate the therapeutic potential of SagA inhibitor pghi-4 as an antibiotic adjuvant, we evaluated its ability to potentiate vancomycin activity using cellular and mouse models of VREfm infection. The combination of pghi-4 and vancomycin, but not either agent alone, reduced VREfm infection of murine (RAW264.7) (Fig. 4j) and human (THP-1) monocytes (Extended Data Fig. 19a) in a dose-dependent manner, that was not attributed to pghi-4 cytotoxicity (Extended Data Fig. 19b, c). We next tested therapeutic efficacy of the combination therapy *in vivo*. A single dose of pghi-4 in combination with vancomycin did not significantly decrease VREfm counts in mouse organs 6- or 24-hours post-infection compared to PBS treatment (Extended Data Fig. 20). However, a two-dose therapeutic regimen significantly reduced weight loss in VREfm-infected mice co-treated with vancomycin and pghi-4 (Fig. 4k). Colony forming unit (CFU) analysis of spleen and liver further demonstrated a significant reduction of VREfm burden in the vancomycin and pghi-4 co-treated group compared to PBS, while monotherapies had no significant effect (Fig. 4l). These results demonstrate that pharmacological inactivation of peptidoglycan remodeling in combination with vancomycin can attenuate VREfm infection *ex vivo* and VREfm-induced sepsis *in vivo*.

## Discussion

Bacterial infections cause a significant healthcare and financial burden worldwide. While antibiotics help manage bacterial infections, many bacterial pathogens have acquired antibiotic resistance and now are difficult to treat, with 4.7 million deaths associated with AMR worldwide in 2019.^40^ Progress in development of new antibiotic agents and targets have resulted in novel approaches to combat antibiotic-resistant bacteria.^22,41–46^ Recently, combination therapeutic approaches using antibiotic adjuvants (non-antimicrobial agents enhancing antibiotic activity) have provided new entities to extend the lifespan of clinical antibiotics.^47–52^ For example, combination treatment with β-lactamase inhibitors (BLIs) overcame antibiotic resistance and restored β-lactam activity against bloodstream infections in hematological neutropenic patients.^53^ This therapeutic combination of an antibiotic adjuvant coupled with an existing antibiotic showed clinical potential and is promising for further development, however, antibiotic adjuvants for other classes of antibiotics are underdeveloped. The alarming increase of VREfm infections^54,55^ and evolution of antibiotic resistance in patients highlights the urgent need for new therapeutic approaches to overcome resistance and prevent adaptation in vulnerable hosts.^39^

While natural products and their derivatives have been reported to broadly inhibit peptidoglycan remodeling or target CHAP (cysteine, histidine-dependent amidohydrolases/peptidases) domain containing hydrolases in other Gram-positive bacterial pathogens,^56,57^ our studies demonstrate NlpC/p60 peptidoglycan hydrolases and their inhibitors may serve as important new antibiotic targets and agents. Here we demonstrated that SagA, a NlpC/p60 hydrolase important for peptidoglycan remodeling, modulates vancomycin susceptibility in VREfm and can be pharmacologically targeted for improved combination therapy (Extended Data Fig. 21). Unlike other NlpC/P60 hydrolases that are in VREfm strains (Extended Data Fig. 2), only deletion of *sagA* compromised cell growth, separation, peptidoglycan remodeling and increased VREfm susceptibility to vancomycin *ex vivo* and VREfm-induced sepsis *in vivo*, which could be rescued by *sagA* re-expression. Long-term serial passaging of the Δ*sagA* strain under sub-MIC vancomycin pressure did not substantially change vancomycin susceptibility (Extended Data Fig. 6e). While we have shown that SagA expression in *E. faecium* and probiotic bacterial species can promote intestinal immunity^14,15^ and cancer immunotherapy *in vivo*^17^, the VREfm Δ*sagA* mutant strain is not less pathogenic in this mouse peritonitis infection model, but is more susceptible to vancomycin *in vivo* (Fig. 2f,g). These results suggest SagA does not significantly contribute to VREfm-induced sepsis *in vivo*, but can be target for antibiotic adjuvants.

Based on these observations, we identified the first-in-class NlpC/p60 peptidoglycan hydrolase inhibitors (pghi-1 to 5) and demonstrated that these β-chloro alkenyl sulfonyl fluorides covalently label the active site of SagA and inhibit peptidoglycan hydrolase activity *in vitro*. We previously reported the synthesis of β-chloro alkenyl sulfonyl fluorides,^24^ however, their activity on NlpC/p60 peptidoglycan hydrolases was not evaluated. Notably, pghi-4 effectively inhibited peptidoglycan remodeling and increased vancomycin susceptibility in several VREfm strains *ex vivo* (Fig. 4i and Extended Data Fig. 18). Interestingly, pghi-4 showed 8-fold enhanced vancomycin susceptibility in VREfm ERV165 compared to the 2-fold enhanced vancomycin activity in the isogenic Δ*sagA* mutant strain (Figs. 2b, 4c and Extended Date Tables 2, 7). Moreover, pghi-4-treated VREfm ERV165 showed increased levels of vancomycin-target D-Ala-D-Ala-containing peptidoglycan fragments compared to the isogenic Δ*sagA* mutant strain (Extended Data Figs. 5 and 16). These observations suggests that β-chloro alkenyl sulfonyl fluorides, such as pghi-4, may target additional peptidoglycan remodeling enzymes in VREfm beyond SagA. In fact, we showed that β-chloro alkenyl sulfonyl fluorides, including pghi-4, can also inhibit the PGH2 NlpC/p60 endopeptidases *in vitro* (Extended Data Fig. 11c). Even though genetic deletion of other NlpC/P60 hydrolases (*pgh2*, *pgh3*, and *pgh4*) did not cause growth defect or changes in vancomycin susceptibility (Extended Data Fig. 2e and 6d), pharmacological inhibition of both SagA and PGH2 VREfm NlpC/P60 hydrolase may contribute to the overall observed activity of pghi-4 in VREfm strains. Although our quantitative competitive chemoproteomic analysis of cysteine-reactive proteins with pghi-4 did not reveal other NlpC/P60 hydrolases as potential targets in VREfm (Extended Data Fig. 17), these enzymes may not be effectively labeled by iodoacetamide reagents. It is also possible that pghi-4 may react with other nucleophilic amino acids on other proteins that were not identified in our analysis. The direct analysis of pghi-4 targets will require the generation of β-chloro alkenyl sulfonyl fluoride probes that are unfortunately not accessible by our current synthetic methods and will require the development of next-generation NlpC/P60 hydrolase inhibitor and probes. Nonetheless, our discovery and development of β-chloro alkenyl sulfonyl fluorides as covalent NlpC/P60 hydrolase inhibitors demonstrate pharmacological inhibition of peptidoglycan remodeling can improve vancomycin activity in VREfm and attenuate the infection of macrophages *ex vivo* (Fig. 4j) and VREfm-induced sepsis *in vivo* (Fig. 4k-l). The further development of more potent NlpC/p60 hydrolase inhibitors should afford new antimicrobial adjuvants to prevent and treat VREfm infections.

## Methods

### Chemistry

The synthesis and characterization of sulfonyl fluorides identified as SagA inhibitors in this work are reported elsewhere ^24^. Characterization of resynthesized pghi-4 matched the previous report (Extended Data Fig. 22-25). ^1^H NMR (400 MHz, CDCl_3_) δ 7.36 (dd, *J* = 8.5, 2.4 Hz, 1H), 7.17 (d, *J* = 2.3 Hz, 1H), 6.96 (d, *J* = 2.3 Hz, 1H), 6.93 (d, *J* = 8.6 Hz, 1H), 4.88 (tt, *J* = 6.3, 3.1 Hz, 1H), 3.93 (s, 3H), 2.09 – 1.98 (m, 2H), 1.97 – 1.89 (m, 3H), 1.90 – 1.80 (m, 2H), 1.67 (tdd, *J* = 10.6, 7.5, 4.8 Hz, 2H); ^13^C NMR (101 MHz, CDCl_3_) δ 152.4, 152.1, 150.0, 125.7, 121.8, 115.4, 115.2, 113.6, 110.7, 80.9, 56.4, 33.0, 24.3; ^19^F NMR (377 MHz, CDCl_3_) δ 65.0; LCMS (ESI^+^): calculated for C_14_H_16_ClFO_4_SNa [M+H]^+^: m/z = 335.05, m/z found 335.09.

### Bacteria

The bacterial species used in this study are listed in Extended Data Table 10. All *Enterococcus* were grown aerobically at 37 °C at 200 RPM shaking in Brain Heart Infusion (BHI) broth (Fisher Scientific, 237500) with appropriate antibiotics. Following Clinical & Laboratory Standards Institute (CLSI) guidelines, vancomycin susceptibility of *E. faecium* was evaluated in Mueller Hinton Broth (MHB, BD 275730). *E. coli* was grown in Luria-Bertani (LB, BD 244610) broth.

### Phylogenetic analysis

To understand the distribution of NlpC/P60 hydrolases in vancomycin-susceptible and -resistant *E. faecium* strains we downloaded all complete genomes of *E. faecium* on NCBI (n= 559 as of January 2026), consisting of genotypically vancomycin-susceptible (n=164) and -resistant E. faecium (n=395) isolates from 99 sequence types. *In silico* multi-locus sequence typing (MLST) was assigned using the program mlst (https://github.com/tseemann/mlst) (v2.19.0). The genome assemblies were screened for antimicrobial resistance determinants using abriTAMR^58^ (v1.0.18) with default settings.

All assemblies were annotated using the run_prokka function in Panaroo (v1.2.10) with clean-mode set to strict, which annotates each sample with the same gene model using Prokka (v1.14.6).^59,60^ The pangenome was defined using Panaroo (v1.2.10), which utilizes a pangenome graph-based approach for clustering. Core genes were defined as genes present in >99% of strains, with accessory in at least >1%. From the pangenome, functional annotation was assigned using eggNOG-mapper (v2.1.2) with default Diamond mode.^61^ All NlpC/P60 hydrolases were identified using the Clusters of Orthologous Genes identifier COG0791 and manually verified using CD-search (v3.2). A maximum-likelihood phylogenetic tree using the alignment of core genes (core_gene_alignment_filtered.aln) was inferred using IQ-TREE (v2.1.4)^62^ with a general time-reversible (GTR+G4) substitution model and 1,000 bootstrap replicates. All figures were generated in R (v.4.3.0, https://www.r-project.org/) using tidyverse (v.1.3.1), patchwork (v.1.1.1), ggtree (v.3.8.2), and ggnewscale (v.0.4.5).

### Plasmids construction

#### 1. pPK99 (pJC005.gent-Δ*sagA*)

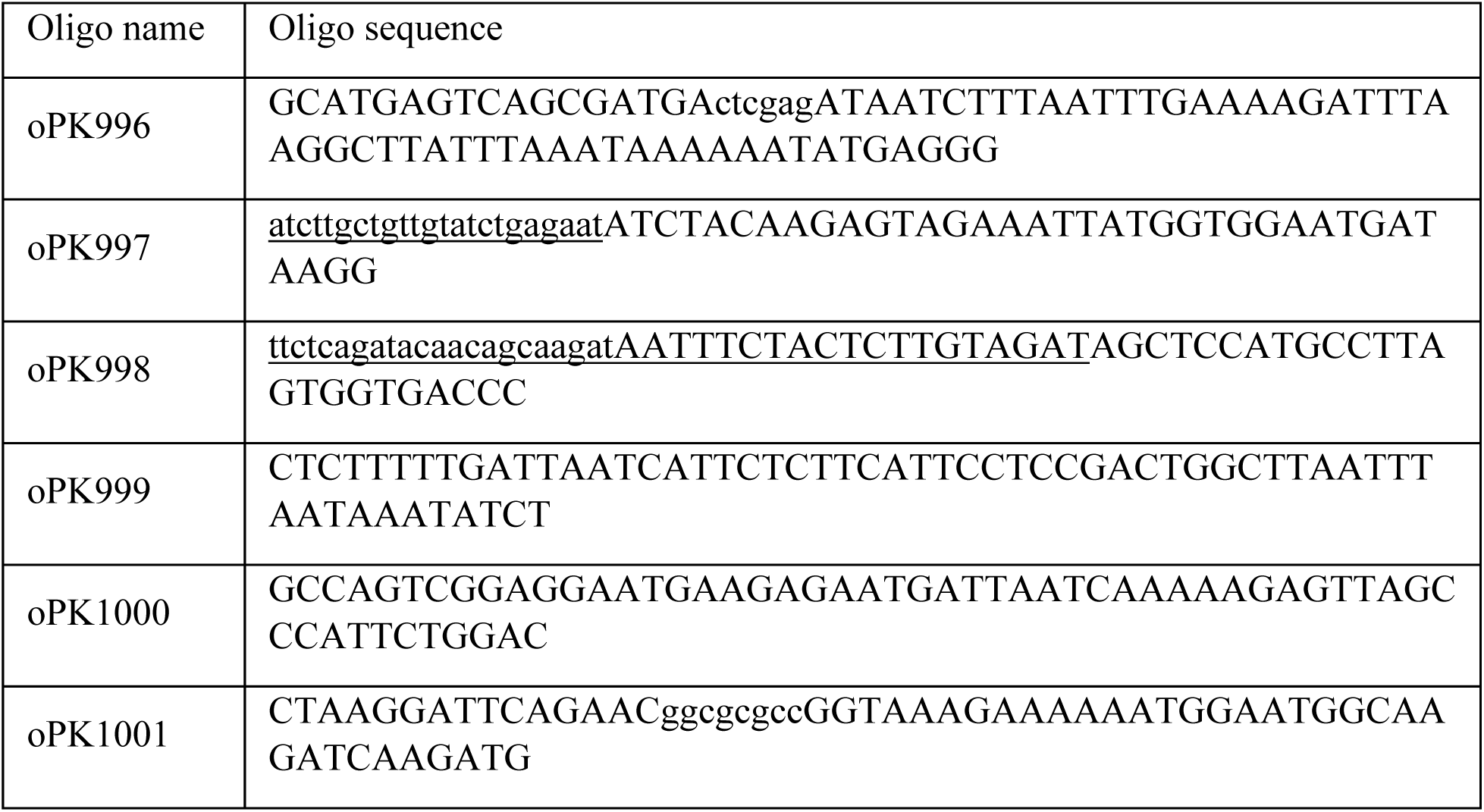

Using oligos oPK996 and oPK997, the sRNA promoter driving the *sagA* protospacer was PCR-amplified (using pUCsRNAP plasmid as template). oPK996 contains an XhoI restriction site (indicated in lowercase), and oPK997 includes the *sagA* protospacer from *E. faecium* ERV165 (indicated in lowercase and underlined) to serve as the CRISPR-Cas12a target for counter-selection during the recombineering process. Similarly, oPK998 (which includes the identical *sagA* protospacer sequence in underlined lowercase and a repeat region in underlined uppercase) and oPK999 were used to amplify the upstream flanking region of *sagA* using *E. fm* ERV165 genomic DNA (gDNA) as template, while oPK1000 and oPK1001 (the latter containing an AscI site, shown in lowercase) amplified the downstream *sagA* flanking region using the same gDNA as template. All oligos were designed such that the three resulting PCR products contained 35–40 bp overlapping regions and were assembled using splicing by overlap extension (SOE) PCR. The resulting fragment was cloned into the pJC005.gent vector via XhoI and AscI restriction digestion followed by ligation. Positive clones were screened in *E. coli* NEB-5α, yielding the construct pPK99. This plasmid was then transformed into *E. faecium* ERV165 for generation of the clean *sagA* deletion mutant (sPK377), as described below.

#### 2. pPK158 (pJC005.gent^R^-*sagA* chromosomal complementation plasmid)

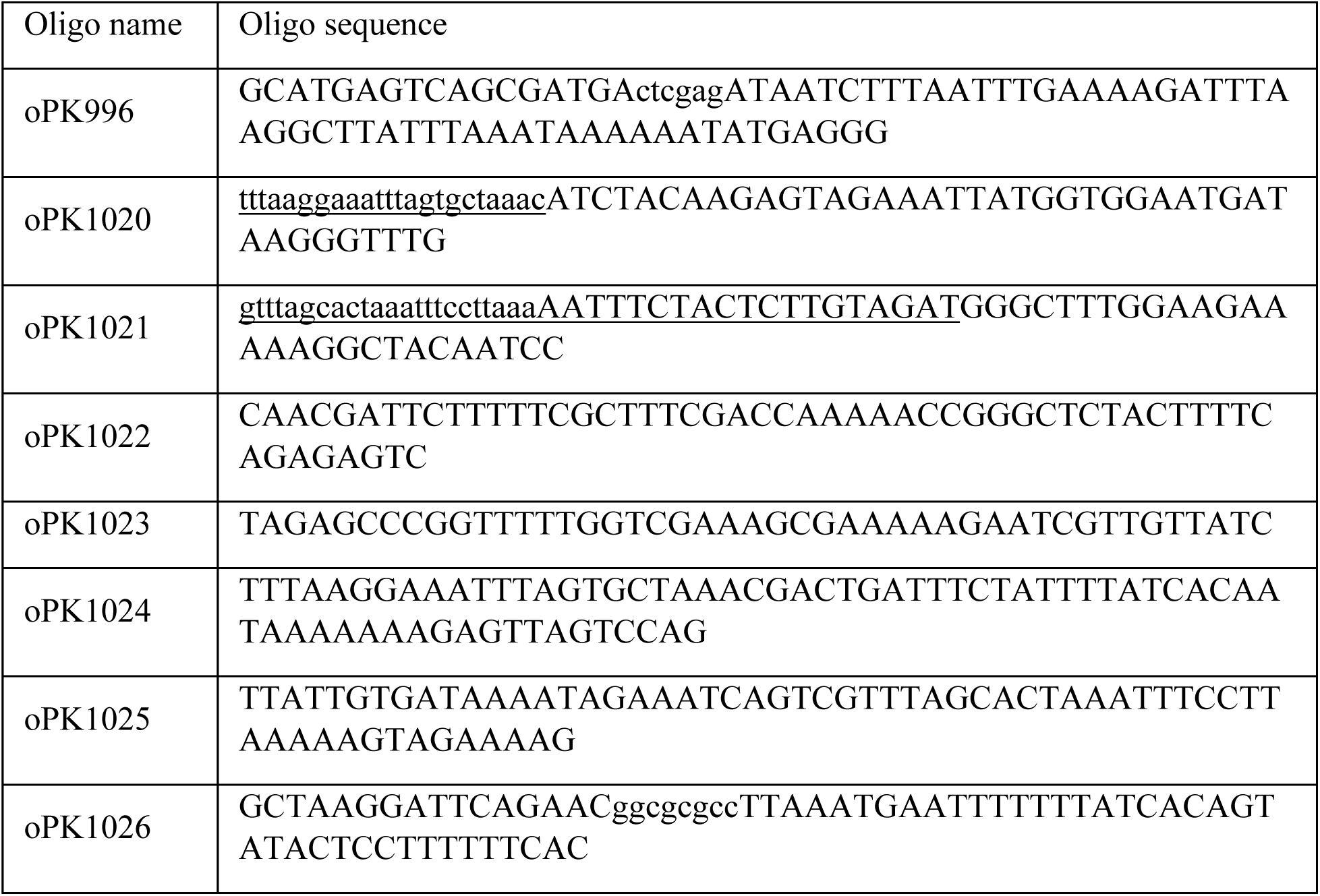

For *sagA* chromosomal complementation in the VREfm ERV165 Δ*sagA* clean deletion mutant (sPK377), a neutral chromosomal locus was selected downstream of the Holliday junction resolvase gene *ruvX*, where no signatures of nearby gene promoters or terminators were detected (Extended Data Fig. 1b). This site contained a protospacer adjacent motif (PAM), making it suitable for CRISPR-Cas12a-based counter-selection. The *sagA* complementation plasmid (pPK158) was constructed by integrating the *sagA* gene into the selected PAM site, thereby disrupting it for CRISPR-Cas12a-based counter-selection. The construct included all essential regulatory elements of ERV165 *sagA*: the native *sagA* promoter, ribosome binding site (RBS), open reading frame (ORF), and transcriptional terminator.

Briefly, the small RNA (sRNA) promoter, derived from the pUCsRNAP template and driving the neutral locus protospacer, was PCR-amplified using oligos oPK996 and oPK1020. oPK996 includes an XhoI restriction site (in lowercase), while oPK1020 contains the neutral locus protospacer from *E. faecium* ERV165 (in lowercase and underlined), which served as the CRISPR-Cas12a target during recombineering. The upstream region of the neutral locus was amplified using *sagA* clean deletion (sPK377) gDNA as template with oligos oPK1021 and oPK1022. oPK1021 has the protospacer (underlined lower case) and a repeat region (underlined uppercase). The ERV165 *sagA* promoter, RBS, ORF, and both translational and transcriptional terminators were amplified using oPK1023 and oPK1034 using *E. fm* ERV165 gDNA as template. Finally, the downstream region of the neutral locus was amplified using sPK377 gDNA as template with oPK1025 and oPK1026, which include AscI restriction sites (in lowercase). All oligos were designed such that the three resulting PCR products contained 35–40 bp overlapping regions and were assembled using splicing by overlap extension (SOE) PCR. The resulting fragment was cloned into the pJC005.gent vector *via* XhoI and AscI restriction digestion followed by ligation. Positive clones were screened in *E. coli* NEB-5α, yielding the construct pPK158. This plasmid was then transformed into *E. faecium* ERV165 for generation of *sagA* chromosomal complementation strain in Δ*sagA* clean deletion mutant (sPK394), as described below.

#### 3. pPK156 (pJC005.gent- Δ*pgh2*)

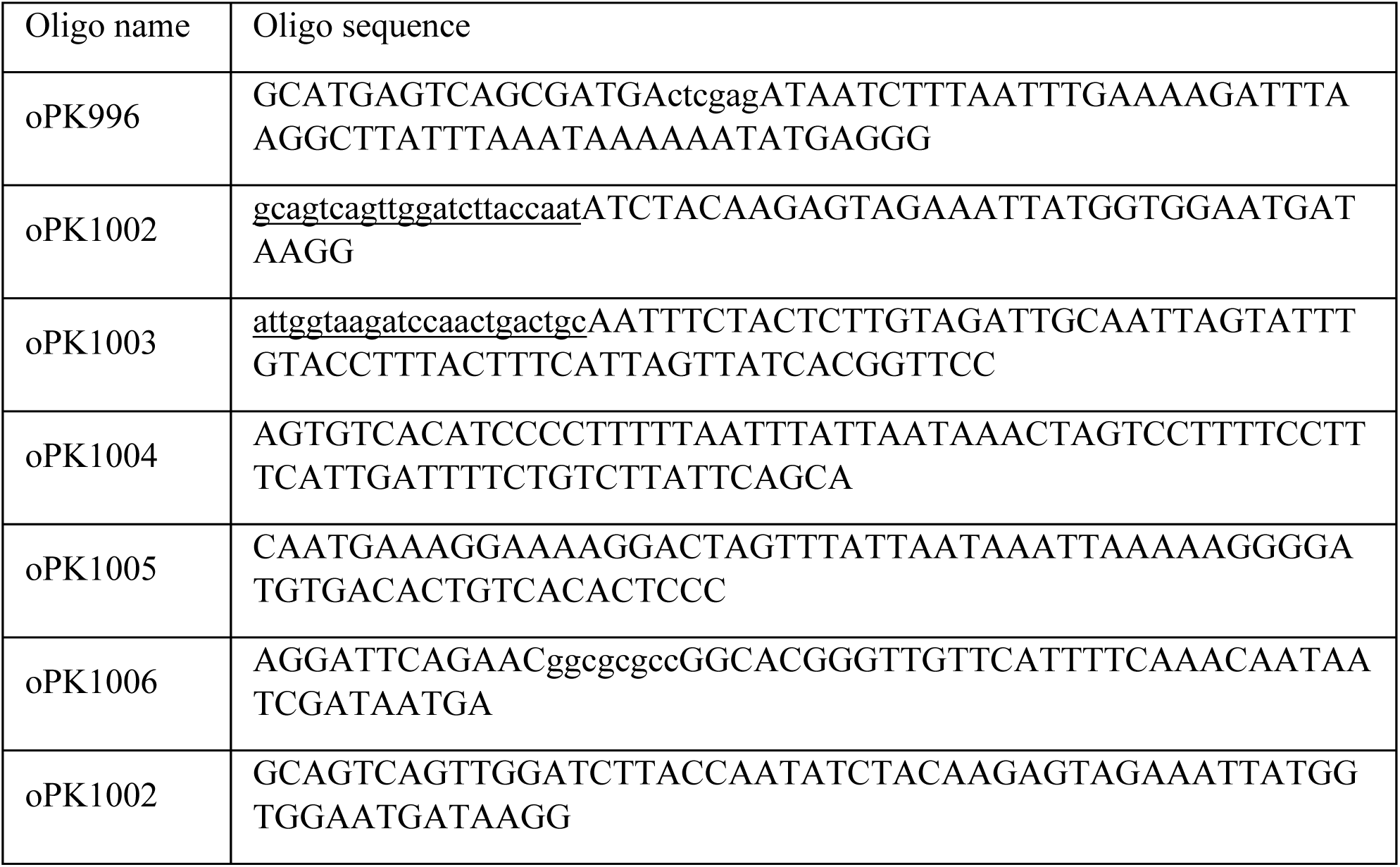

Using oligos oPK996 and oPK1002, the sRNA promoter driving the *pgh2* protospacer was PCR-amplified (using pUCsRNAP plasmid as template). oPK996 contains an XhoI restriction site (indicated in lowercase), and oPK1002 includes the *pgh2* protospacer from *E. faecium* ERV165 (indicated in lowercase and underlined) to serve as the CRISPR-Cas12a target for counter-selection during the recombineering process. Similarly, oPK1003 (which includes the identical *pgh2* protospacer sequence in underlined lowercase and a repeat region in underlined uppercase) and oPK1004 were used to amplify the upstream flanking region of *pgh2* using *E. fm* ERV165 genomic DNA (gDNA) as template, while oPK1005 and oPK1006 (the latter containing an AscI site, shown in lowercase) amplified the downstream *pgh2* flanking region using the same gDNA as template. All oligos were designed such that the three resulting PCR products contained 35–40 bp overlapping regions and were assembled using splicing by overlap extension (SOE) PCR. The resulting fragment was cloned into the pJC005.gent vector via XhoI and AscI restriction digestion followed by ligation. Positive clones were screened in *E. coli* NEB-5α, yielding the construct pPK156. This plasmid was then transformed into *E. faecium* ERV165 for generation of the clean *pgh2* deletion mutant, as described below.

#### 4. pPK107 (pJC005.gent- Δ*pgh3*)

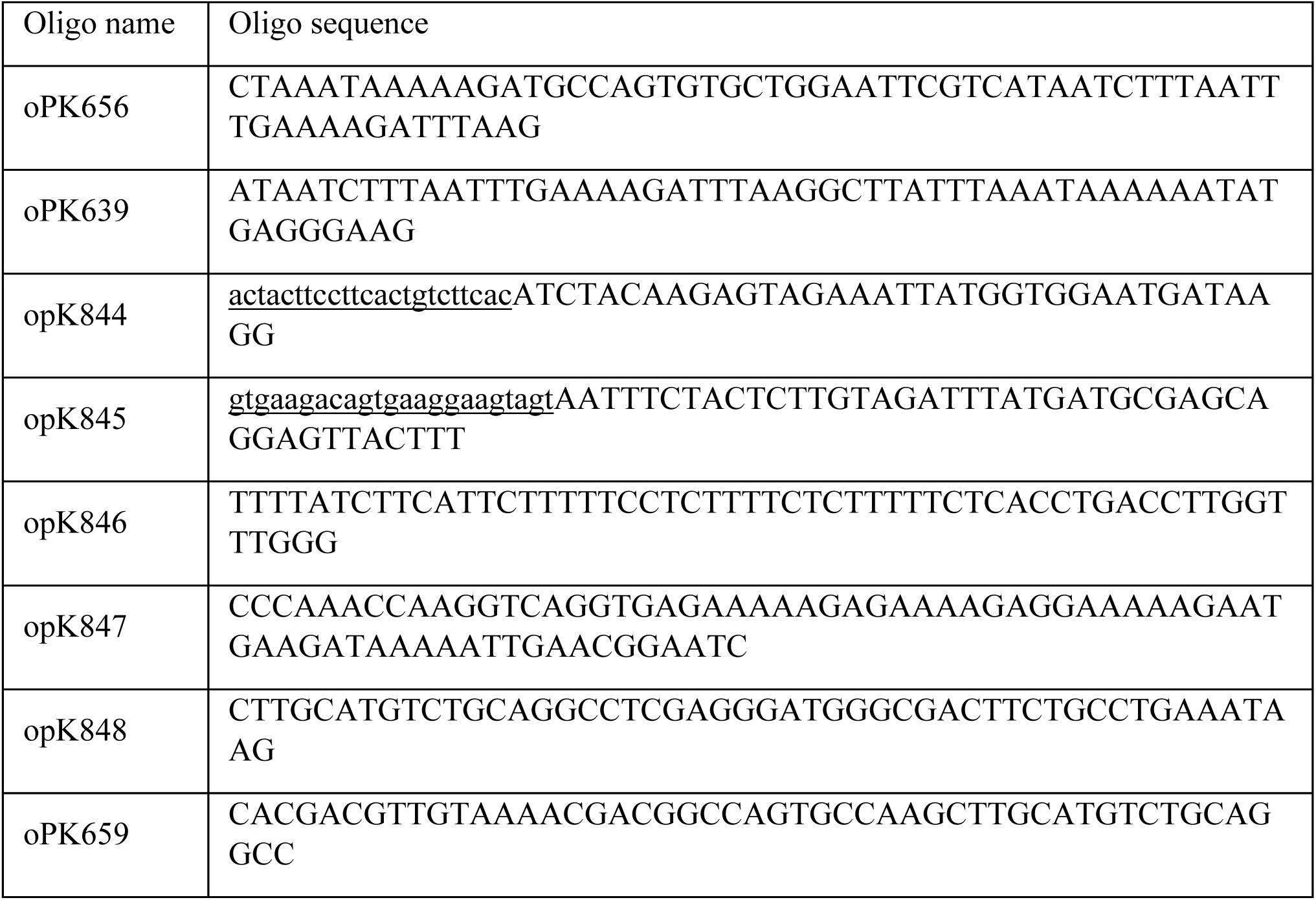

Using oligos oPK639, the sRNA promoter driving the *pgh3* protospacer was PCR-amplified (using pUCsRNAP plasmid as template). oPK844 contains *pgh3* protospacer from *E. faecium* ERV165 (indicated in lowercase and underlined) to serve as the CRISPR-Cas12a target for counter-selection during the recombineering process. Similarly, oPK845 (which includes the identical *pgh3* protospacer sequence in underlined lowercase and a repeat region in underlined uppercase) and oPK846 were used to amplify the upstream flanking region of *pgh3* using *E. fm* ERV165 genomic DNA (gDNA) as template, while oPK847 and oPK848 amplified the downstream *pgh3* flanking region using the same gDNA as template. All oligos were designed such that the three resulting PCR products contained 35–40 bp overlapping regions and were assembled using splicing by overlap extension (SOE) PCR using oPK656 and oPK659. The resulting fragment was cloned into the pJC005.gent vector via XhoI and AscI restriction digestion followed by ligation. Positive clones were screened in *E. coli* NEB-5α, yielding the construct pPK107. This plasmid was then transformed into *E. faecium* ERV165 for generation of the clean *pgh3* deletion mutant, as described below.

#### 5. pPK154 (pJC005.gent- Δ*pgh4*)

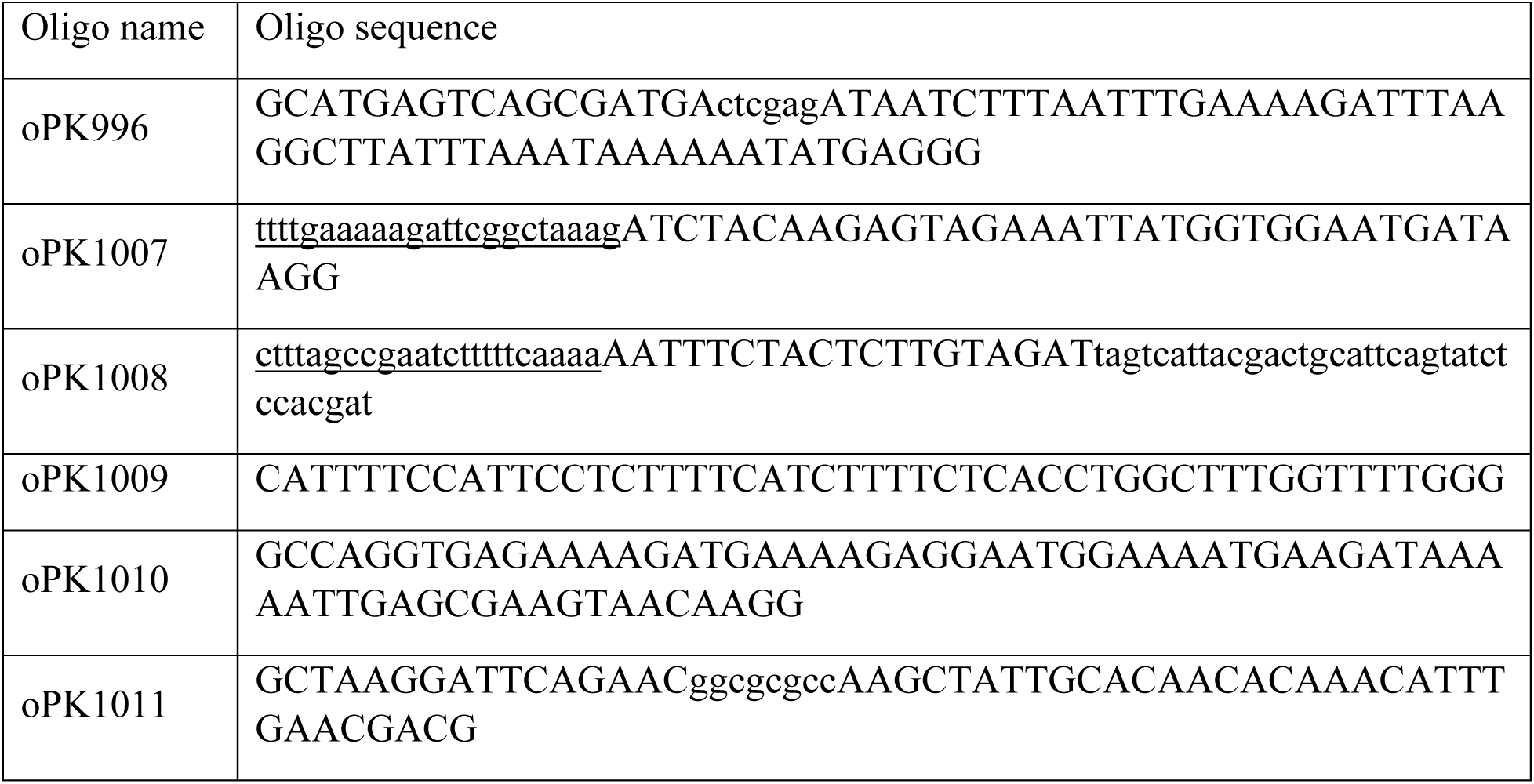

Using oligos oPK996 and oPK1007, the sRNA promoter driving the *pgh4* protospacer was PCR-amplified (using pUCsRNAP plasmid as template). oPK996 contains an XhoI restriction site (indicated in lowercase), and oPK1007 includes the *pgh4* protospacer from *E. faecium* ERV165 (indicated in lowercase and underlined) to serve as the CRISPR-Cas12a target for counter-selection during the recombineering process. Similarly, oPK1008 (which includes the identical *pgh4* protospacer sequence in underlined lowercase and a repeat region in underlined uppercase) and oPK1009 were used to amplify the upstream flanking region of *pgh4* using *E. fm* ERV165 genomic DNA (gDNA) as template, while oPK1010 and oPK1011 (the latter containing an AscI site, shown in lowercase) amplified the downstream *pgh4* flanking region using the same gDNA as template. All oligos were designed such that the three resulting PCR products contained 35–40 bp overlapping regions and were assembled using splicing by overlap extension (SOE) PCR. The resulting fragment was cloned into the pJC005.gent vector via XhoI and AscI restriction digestion followed by ligation. Positive clones were screened in *E. coli* NEB-5α, yielding the construct pPK154. This plasmid was then transformed into *E. faecium* ERV165 for generation of the clean *pgh4* deletion mutant, as described below.

### Preparation of electrocompetent *E. faecium* ERV165 cells and electroporation

The protocol described below is a standardized procedure used in our lab. A single colony of *E. faecium* ERV165 and their mutant derivatives were inoculated into 4 mL of plain BHI and grown overnight. The culture was then diluted 1:50 into 200 mL GS-BHI (BHI + 4% glycine + 0.5 M sucrose) and incubated overnight (14–16 hours). From this point onward, cells were handled gently to preserve viability. Cells were harvested by centrifugation at 1,000 × *g* for 15 min at room temperature (RT), gently resuspended in 100 mL GS-BHI (as described below), and incubated without shaking at 37°C for 1.5 hours. Cells were re-pelleted (1,000 × *g*, 15 min, 4°C), gently washed with 35–40 mL ice-cold electroporation solution, and kept on ice. The harvested cells were then resuspended in 1–2 mL electroporation solution and homogenized using a sterile serological pipette. Aliquots (100 µL) were prepared in pre-chilled Eppendorf tubes and stored at –80°C. For electroporation, thawed 100 µL aliquots were mixed with 2-4 µg DNA and transferred to 0.2 cm gap cuvettes. Electroporation was carried out at 25 µF, 400 Ω, and 2.5 kV. Immediately after, 0.9 mL of 1X SBHI was added, and cells were recovered for 3 h at 37°C without shaking before plating on selective BHI agar. For recombineering, the entire transformation mixture was pelleted (7,000 *× g*, 2 min) and plated to maximize recovery.

### Generation of the ERV165 Δ*sagA* clean deletion mutant (Δ*sagA*)

The gene-editing CRISPR-cas12a protocol from Chua et al.^29^ was modified to account for the robustness of the Δ*sagA* mutant, which made deletion challenging.

#### a. Generation of the ΔsagA mutant

Briefly, pPK99 plasmid was electroporated into VREfm ERV165 electrocompetent cells. A few transformants were inoculated into 5 mL of BHI broth supplemented with 250 µg/mL gentamycin and grown at 37°C with shaking for two days. Sub-culture (1:1,000 dilution) into fresh BHI broth supplemented with 250 µg/mL gentamycin and incubated under the same conditions for an additional two days. Subsequently, 1 µL of the culture was streaked onto BHI agar plates supplemented with 250 µg/mL gentamycin and 250 ng/mL anhydrotetracycline (ahTC) and incubated at 37°C for 3 to 5 days. Smaller single colonies were picked and grown in BHI broth supplemented with 250 µg/mL gentamycin and 250 ng/mL ahTC. Since D*sagA* mutant exhibits a sedimentation phenotype, the sedimented cells were selectively taken and re-inoculated into fresh BHI broth with the same supplements. This enrichment step was repeated twice to enhance the recovery of Δ*sagA* mutants.

#### b. Morphology-based screening and validation of ΔsagA mutants

Based on our previous studies ^19^, the *E. faecium* Com15 Δ*sagA* mutant is known to exhibit growth defects. Therefore, numerous sick colonies were screened but were ultimately found to be false positives. During this study, we identified an alternative screening method for Δ*sagA* mutants, as they exhibit a distinct colony morphology that can be visualized using light microscopy (Fig. 1d). To identify such mutants, BHI plates were continuously monitored for the appearance of smaller and sick colonies. Numerous BHI plates were examined under a light microscope using differential interference contrast (DIC) imaging with a 10× objective lens, and colonies were extensively analyzed for the characteristic Δ*sagA* mutant morphology. Colonies with a smaller size and a rough halo texture were selected for further screening by colony PCR. However, DNA sequencing analysis revealed a high frequency of false positives. In these cases, the *sagA* open reading frame (ORF) remained intact, but the protospacer was disrupted,likely due to CRISPR-Cas12a-mediated double-stranded break and repair events, leading to a phenotype that mimicked the *sagA* mutant. Despite these challenges, we successfully isolated a single, confirmed colony that carried a clean deletion of *sagA*. The whole genome sequence analysis was done to check for background mutations (Extended Data Tables 3 and 4).

### Generation of ERV165 Δ*sagA* chromosomal complementation strain (Δ*sagA*::*sagA*)

The complementation strain was constructed using the pPK158 plasmid, following a method similar to that described above for generating the Δ*sagA* mutant. Whole-genome sequencing analysis confirmed that the observed growth defect of Δ*sagA* mutant was not due to background mutations (Extended Data Tables 5 and 6).

### Generation of the ERV165 *pgh2, pgh3 and pgh4* clean deletion mutants (Δ*pgh2*, Δ*pgh3* Δ*pgh4*)

Strains were constructed using the pPK156, pPK107 or pPK154 plasmids respectively, following a method similar to that described above for generating the Δ*sagA* mutant.

### Adaptive laboratory evolution

VREfm cultures grown overnight from single colonies were subcultured in fresh BHI supplemented with vancomycin (50 µg/mL) and were grown aerobically at 37 °C at 200 RPM shaking overnight. The next day, the cultures were passaged in fresh BHI supplemented with vancomycin (50 µg/mL) to reach OD_600_∼0.1 and were grown aerobically at 37 °C at 200 RPM shaking overnight. The passaging was repeated for 14 days and the vancomycin susceptibility was determined following CLSI guidelines.

### Recombinant protein expression and purification

Truncated SagA_NlpC/P60 proteins were expressed and purified from *E. coli* BL21-RIL (DE3) as previously described.^16,36^ pET-21a(+) plasmids containing the truncated SagA_NlpC/P60 genes with a C-terminal His_6_ tag were transformed into BL21-CodonPlus (DE3)-RIL *E. coli* (Agilent 230245) according to the manufacturer’s protocol and maintained in LB broth supplemented with 100 µg mL^-1^ ampicillin and 25 µg mL^-1^ chloramphenicol. Overnight bacterial culture was subcultured in 1 L fresh BHI supplemented with appropriate antibiotics, grown until OD_600_∼0.5, induced with 1 mM isopropyl-*D*-thiogalactopyranoside, and additionally grown for 2 hours at 37 °C and 200 RPM shaking. Bacterial cells were collected by centrifugation at 4 °C, resuspended in 20 mL lysis buffer (20 mM Tris-HCl, pH 8.0, 150 mM NaCl, 0.1% SDS, 0.025 U/mL benzonase, and 1*×* protease inhibitor cocktail). After 15 min of sonication followed by centrifugation at 18,000 *× g* for 30 min at 4°C, the supernatant containing the soluble target protein was collected and loaded on 2 mL of Ni-NTA agarose (Invitrogen) equilibrated with the binding buffer (PBS). The protein-bound resin was washed with 20 mM and 40 mM imidazole sequentially, then SagA_NlpC/P60 protein was eluted with 300 mM imidazole. Semi-purified protein was dialyzed into PBS buffer at 4°C overnight using 10K MWCO Slide-A-Lyzer MINI dialysis devices (Thermo Fisher Scientific). Protein was further purified on a ENrich™ SEC 650 column (Bio-Rad) pre-equilibrated with PBS using NGC chromatography system (Bio-Rad). Fractions containing the target protein were combined and concentrated. Protein concentration was estimated by Pierce™ BCA Protein Assay (ThermoFisher) and protein was stored at −80°C.

### Fluorescence polarization (FP) cysteine activity assay

The assay is based on competitive reaction of a tested compound with broad-spectrum cysteine-reactive probe tetramethylrhodamine-5-iodoacetamide (Anaspec) and was adapted from ^63^. 16 μL of recombinant SagA_NlpC/P60 (0.5 μM) in PBS was added to black 384-well plate. Tested compound (50 μM) were then added to corresponding wells using Bravo instrument (Agilent). The plate was covered with aluminum seal and incubated for 1 hour at room temperature with agitation, followed by treatment with tetramethylrhodamine-5-iodoacetamide (20 nM) for 30 min at 37°C shielded from light with agitation. Fluorescence polarization was measured using EnVision plate reader 2105 (Perkin Elmer) with BODIPY TMR FP filter set and calculated as follows:

Polarization (mP) = 1000 * (S – G*P)/(S + G*P), where S and P are the measured results with the S and P emission filters respectively. G is a correction factor (effect of emission filter transmission variations, differences in the emission light paths and sample viscosity).

Inhibition (%) = 100-(mP_sample_-mP_negative_)/(mP_Positive_-mP_negative_), where mP_sample_ is FP value of SagA_NlpC/P60 treated with a corresponding testing compound, mP_positive_ is FP value of SagA_NlpC/P60 treated with DMSO, mP_negative_ is FP value of SagA_NlpC/P60 (C433A, inactive mutant) treated with DMSO. Calculated inhibition > 100% was assigned as 100%.

### Gel-based cysteine activity assay

The assay was adapted from ^63^. For gel-based SagA activity assay, the conditions were similar to FP assay, except for treatment with tetramethylrhodamine-5-iodoacetamide (50 nM). The reaction was quenched with 4 × Laemmli buffer, heat-inactivated for 5 min at 90 °C, separated by SDS-PAGE. The gel was visualized using ChemiDoc MP imaging system (Bio-Rad). Relative labeling was determined by extracting fluorescence intensity of the band and compared to DMSO control. For analysis of bacterial proteome, after treatment with pghi-4, 100 μg of precipitated secreted proteins from supernatants or bacterial proteome from cell pellets lysed in PBS were treated with *N*-5-hexyn-1-yl-2-iodoacetamide (IA-alk, 100 μM) for 1 hour at room temperature with agitation, followed by 1 hour treatment with click mixture: 1 mM tris(2-carboxyethyl)phosphine hydrochloride (TCEP), 1 mM CuSO_4_, 0.1 mM tris[(1-benzyl-1H-1,2,3-triazol-4-yl)methyl]amine (TBTA), and 100 μM rhodamine-azide. The reactions were quenched by addition 4 × Laemmli buffer and proteins were separated by SDS-PAGE. In-gel fluorescence was detected by ChemiDoc MP imaging system (Bio-Rad). Relative labeling was determined by extracting fluorescence intensity of the SagA band and compared to DMSO control using Image Lab software (Bio-Rad). Protein loading was analyzed by Coomassie blue staining for recombinant SagA and by α-SagA Western Blot for bacterial proteome.

Biotin pull-down protocol was adapted from ^64^. For biotin pull-down experiments from bacterial lysates, 250 μg of each total cell lysates in PBS were treated IA-alk (100 μM) for 1 hour at room temperature with agitation, followed by 1 hour treatment with click mixture: 1 mM tris(2-carboxyethyl)phosphine hydrochloride (TCEP), 1 mM CuSO_4_, 0.1 mM tris[(1-benzyl-1*H*-1,2,3-triazol-4-yl)methyl]amine (TBTA), and 100 μM biotin-azide. Proteins were precipitated with 4 × volume cold methanol. Precipitated proteins were pelleted by centrifugation (18,000 × *g*, 4°C, 10 min), sequentially washed with cold methanol and centrifuged 3 times, followed by drying in SpeedVac. Resulted pellets were resuspended in 100 μL 4% SDS in PBS with bath sonication. 2.5% of solution was used as input (protein loading control). Total volume of incubated with 20 μL PBS-T-washed High Capacity NeutrAvidin agarose (Pierce) (500 μL PBS-T-washed twice, 2,500 *× g* for 60 s) at room temperature for 1 hour with end-to-end rotation. The agarose was then washed with 500 μL PBS (1% SDS) 3 times, 500 μL 1 M Urea in PBS three times, and 500 μL PBS three times. Samples were boiled with 2 × Laemmli buffer 95°C for 5 min and analyzed by western blot.

For biotin pull-down experiments from bacterial supernatants, proteins were precipitated from supernatants with 4 × volume cold methanol. Precipitated proteins were pelleted by centrifugation (18,000 × *g*, 4°C, 10 min). Resulted pellets were resuspended in PBS and protein concentration was estimated by BCA assay with BCA Protein Assay Kit (Thermo). 250 μg of each total proteins were processed further as described above.

### Competitive cysteine-directed chemoproteomic analysis

The protocol was adapted from ^65^. VREfm ERV165 were grown from overnight culture in 15 mL of fresh BHI till OD_600_∼0.6. Bacteria were then centrifuged and resuspended in 1 mL of BHI (OD_600_∼9), followed by incubation with pghi-4 (10, 25 or 50 μM) for 1 hour at 37°C and 200 RPM shaking. Bacterial pellets were collected by centrifugation (4,800 × *g* for 10 min), washed twice with PBS and immediately processed or stored at −80 °C. Bacterial pellets were resuspended in 360 μL PBS and lysed with 0.1 mm glass beads (BioSpec) using FastPrep system (MP Biomedicals, settings: 6 m/s, 2 cycles, 45 s for each cycle). Proteins were quantified by Pierce™ BCA Protein Assay (ThermoFisher) and normalized to 2 mg/mL. 1 mg/0.5 mL of bacterial proteome was incubated with iodoacetamide-desthiobiotin (IA-DTB, 100 μM) for 1 hour at room temperature agitating. Bacterial proteins were precipitated by cold methanol (600 μL), chloroform (200 μL), and water (100 μL), followed by vortexing and centrifugation at 16,000 × *g* for 10 min at 4°C. Liquids were carefully aspirated, proteins were washed with cold methanol, pelleted by centrifugation (16,000 × *g* for 10 min at 4°C) and air-dried for 5 min. Protein pellets were resuspended in 90 μL of buffer (9 M urea, 10 mM DTT, 50 mM triethylammonium bicarbonate (TEAB) pH 8.5), heated at 65°C for 20 min, followed by treatment with iodoacetamide (50 mM) for 30 min at 37°C. The insoluble residues were pelleted by centrifugation and clear solutions were sonicated. Samples were diluted with 300 μL TEAB buffer and trypsinized (5 μL of 0.4 μg/μL trypsin in trypsin buffer supplemented with 25 mM CaCl_2_) overnight at 37°C. Samples were treated with 400 µL of wash buffer (50 mM TEAB, 150 mM NaCl, 0.2% NP-40) containing 50 µL of streptavidin agarose to the peptide samples, followed by rotation at room temp for 2 hours. Suspensions were briefly centrifuged, and beads-containing suspensions were loaded on BioSpin columns. The beads were sequentially washed with 3 × 1 mL wash buffer, 3 × 1 mL PBS, 3 × 1 mL MiliQ water. Beads-bound peptides were eluted by addition of 2 × 200 µL of 80% acetonitrile (0.1% formic acid) and the eluate was concentrated by SpeedVac. Peptides were resuspended in 70 μL EPPS buffer (200 mM, pH 8.0), supplemented with 30% acetonitrile, vortexed and sonicated for 5 min. Peptides were tandem mass tag (TMT)-labeled by adding 3 μL of 10 mg/mL TMT^10plex^ tag and incubating for 1 hour at room temperature. The reaction was quenched by sequential addition of hydroxylamine (3 μL of a 5% aqueous solution, 15 min at room temperature) and formic acid (5 μL), followed by concentration using SpeedVac. Samples were resuspended in 500 μL Velos buffer A (95% water, 5% acetonitrile, 0.1% formic acid), acidified with 20 μL of formic acid, desalted (Sep-Pak C18 Cartridge) and concentrated by SpeedVac. Desalted and concentrated samples were redissolved in 500 μL Velos buffer A and HPLC-fractionated. Fractionation and TMT LC-MS analysis was followed as previously described ^65^.

Data were processed as previously described^65^ with small modifications regarding the proteome dataset. Raw files were uploaded to the Integrated Proteomics Pipeline (IP2, version 6.0.2) available at http://ip2.scripps.edu/ip2/mainMenu.html, and MS2 and MS3 files were extracted from the raw files using RAW Converter (version 1.1.0.22, available at http://fields.scripps.edu/rawconv/) and searched using the ProLuCID algorithm using the *Enterococcus faecium* ERV165 UniProt database (UP000005678, released 2012-04). Cysteine residues were searched with a static modification for carboxyamidomethylation (+57.02146 Da). A dynamic modification for IA-DTB labeling (+398.25292 Da) was included with a maximum number of two differential modifications per peptide. N termini and lysine residues were also searched with a static modification corresponding to the TMT tag (+229.1629 Da). Peptides were required to be at least 6 amino acids long. ProLuCID data were filtered through DTASelect (version 2.0) to achieve a spectrum false-positive rate below 1%. We excluded nonunique peptides and required at least one tryptic cleavage site and two peptides per protein. The MS3-based peptide quantification was performed with reporter ion mass tolerance set to 20 ppm with the IP2. Pghi-4 cysteine-directed activity was calculated as competition (%) of a cysteine site relative to DMSO treatment (0%) when IA-DTB fully occupies accessible cysteines. The full list of identified proteins is in Supplementary Data 1.

### Peptidoglycan isolation

The peptidoglycan isolation was followed as previously described.^36^ Overnight VREfm cultures were subcultured in fresh BHI supplemented with or without vancomycin, pghi-4 or in combination and were grown till OD_600_∼0.8-1. Bacterial cell pellets were collected by centrifugation and lysed in 0.25% SDS solution in 0.1 M Tris-HCl, pH 6.8 and boiling the suspension for 20 min at 100°C. The insoluble bacterial cell wall was collected by centrifugation, washed with distilled water to remove SDS. The cell wall was next sonicated for 30 min in distilled water and treated with benzonase followed by trypsin digestion. Then, insoluble cell wall was recovered by centrifugation (16,000 *× g*, 10 min, 4°C), and washed with distilled water. Next, the cell wall was treated with 1 M HCl for 4 hours at 37°C and 200 RPM shaking. The insoluble material was collected by centrifugation (16,000 *× g*, 10 min) and washed with distilled water until the pH was 5-6. Purified insoluble peptidoglycan was digested with mutanolysin from *Streptomyces globisporus* (Sigma, 10 KU/mL of mutanolysin in MiliQ H_2_O) in 10 mM sodium phosphate buffer, pH 4.9 for 16 hours at 37°C shaking. The mutanolysin was heat-inactivated and resulting soluble peptidoglycan was used in gel-based peptidoglycan hydrolase activity assay or LC-MS analysis.

### Gel-based peptidoglycan hydrolase activity assay

The peptidoglycan hydrolase activity assay is based on in-gel analysis of peptidoglycan fragments labeled with 8-aminonaphthalene-1,3,6-trisulfonic acid (ANTS). The assay was followed as previously described^36^ with some modifications. 10 μM recombinant SagA_NlpC/P60 (or other NlpC/P60 peptidoglycan hydrolase) was incubated with 250 μg of the mutanolysin-digested peptidoglycan in 50 mM Bis-Tris, pH 5.5 overnight at 37 °C and 220 RPM shaking. The enzymatic activity was heat-inactivated at 100°C for 5 min. The samples were centrifuged at 16,000 x *g* for 5 min and supernatants were transferred and concentrated. The concentrated samples were resuspended and incubated with 5 μL of each reagent (0.2M ANTS in water supplemented with 15% acetic acid and 1M NaBH_3_CN in DMSO) overnight at 37 °C, 220 RPM shaking and protected from light. The reaction mixtures were diluted with 10 μL 50% glycerol (*v/v*) and ANTS-labeled peptidoglycan fragments were separated by Native PAGE on 4-20% Criterion TGX precast gels (Bio-Rad) ran at 100 V for 30 min, and visualized by ChemiDoc MP imaging system (Bio-Rad) and SYBR-Safe settings. Inhibition was determined by extracting fluorescence intensity of the ANTS-labelled enzymatic product (GlcNAc-MurNAc dipeptide) and compared to DMSO control.

### Computational covalent docking

The docking receptor file was prepared from PDB: 6B8C.^16^ All crystallographic waters and alternative residue positions were removed with PyMOL (https://pymol.org/). The receptor was then protonated using Reduce^66^ and prepared for docking using Meeko (https://github.com/forlilab/Meeko) to assign atomtypes and Gasteiger partial charges and convert to a PDBQT file^67^. Gridmaps were calculated with AutoGrid4 with a box size of 16 Å, 18 Å, 27 Å (0.375 Å grid spacing) and with the box center at 101 Å, 80 Å, 140.5 Å. His 506 was also designated to be in the HID (neutral, δ-nitrogen protonated) form. The Cys 443 C⍺ and sidechain were also removed.

Covalent docking as performed using the “flexible side chain” model described by Bianco et al.^68^. 2D line drawings of pghi-4 adducts as either the (a) thioenol sulfonyl fluoride (Michael addition product) and (b) vinylchloride thiosulfonate (sulfonyl fluoride exchange product) were prepared in ChemDraw. The 3D conformation of the vinylchloride thiosulfonate was then prepared with MolScrubber (https://github.com/forlilab/molscrub) and checked to ensure the optimal cyclopentane conformation was generated with the ether in equatorial position. For the thioenol sulfonyl fluoride, a torsional energy scan of the phenyl-thioenol torsion was performed using Gaussian16 with 10-degree incremental steps between energy calculations. Minima were found for torsions of approximately 45° and −143° between the thioenol sulfur and the phenyl carbon adjacent to that bearing the methyl ether. Both thioenol sulfonyl fluoride geometries (with the phenyl-thioenol torsion held as non-rotatable) and the single vinylchloride thiosulfonate were prepared for docking using Meeko to add Gasteiger partial charges and convert to PDBQTs^67^. All three models were then docked as flexible side chains with AutoDock-GPU ^69^, keeping the top scoring pose for further analysis. The two docked thioenol sulfonyl fluoride models were evaluated visually, with the −143° model selected as the most reasonable docked conformation. Models were additionally corroborated using reactive docking^70^ to confirm the state prior to reaction could be accommodated.

### Intact protein analysis by mass spectrometry

10 μM recombinant SagA_NlpC/P60 (or C433A inactive control) was incubated with 50 μM pghi-4 (or inactive pghi-6) in 50 mM Tris-Cl, pH 7.6, 150 mM NaCl buffer for 1 hour (or 16 hours) at room temperature and shaking at 600 RPM. The mixtures were analyzed by ESI-TOF mass spectrometry.

### Protein crystallization

Crystallization efforts to obtain a SagA-pghi-4 complex were extensive and systematic using both co-crystallization and soaking methodologies. High-throughput crystallization screening was performed using the automated Rigaku CrystalMation system at The Scripps Research Institute with the JCSG Core Suite (QIAGEN), comprising 384 distinct conditions across four 96-well plates. Robust diffraction-quality crystals were obtained reproducibly for SagA under multiple conditions for both apo and ligand-incubated samples. For co-crystallization, the inhibitor was added at 5× and 10× molar excess to purified SagA at protein concentrations of 10, 15, and 18 mg/mL, followed by incubation for approximately 30 minutes at either room temperature or 4 °C prior to crystallization setup. In parallel, soaking experiments were performed by incubating pre-grown apo-SagA crystals in pghi-4-containing solutions for varying time intervals before flash freezing. Crystals from both co-crystallization and soaking experiments diffracted strongly to sub-2.0 Å resolution under multiple conditions, including 0.1 M sodium citrate–citric acid (pH 5.6) with ammonium or lithium sulfate, and 0.2 M magnesium acetate with 20% (w/v) PEG 3350. Data were processed and refined using standard crystallographic pipelines^16^. However, in all cases, molecular replacement and subsequent refinement yielded apo SagA structures, with no interpretable ligand-associated electron density observed in unbiased difference or omit maps. These results suggest that, under the crystallization conditions tested, the inhibitor either does not bind SagA with sufficient occupancy to be captured crystallographically, or is destabilized, or displaced during crystal growth or soaking. Alternatively, other factors such as transient binding, competition with crystallization components, or conformational heterogeneity cannot be excluded. Despite extensive crystallization efforts, a SagA-pghi-4 complex structure was not obtained.

### Bacterial growth defect and antibiotic susceptibility assay

Starting cultures were grown from single bacterial colonies overnight at 37°C at 200 RPM shaking in Brain Heart Infusion (BHI) broth. Next, the starting culture were diluted to OD_600_ ∼0.1 in fresh BHI in a sterile 96-well plate and supplemented with vancomycin alone, SagA inhibitor alone or combination of vancomycin and SagA inhibitor at indicated concentrations. The OD_600_ values were measured at 37°C with continuous orbital shaking using BioTek Cytation 5 plate reader (Agilent). For inhibition heat-maps, inhibition was calculated as percentage of OD_600_ values of treatment groups over vehicle control at the single time point. Following Clinical & Laboratory Standards Institute (CLSI) guidelines, vancomycin susceptibilities of ERV165 WT and generated mitants were also tested in Mueller Hinton Broth (MHB). Minimal inhibitory concentrations (MICs) were determined as the lowest concentration that inhibited the visible growth.

### LC-MS-based peptidoglycan analysis

The analysis was followed as we previously described.^19,36^ The mutanolysin-digested peptidoglycan was treated with sodium borohydride in 0.25 M boric acid (pH 9) for 1 hour at room temperature, quenched with orthophosphoric acid, and pH adjusted to 2-3. The samples were centrifuged at 20,000 × *g* for 10 minutes. Then, the reduced peptidoglycan was analyzed by 1290 Infinity II LC/MSD system (Agilent technologies) using Poroshell 120 EC-C18 column (3 × 150 mm, 2.7 μm). Samples were run at flow rate 0.5 mL/min in mobile phase (A: water, 0.1% formic acid) and an eluent (B: acetonitrile, 0.1% formic acid) using following gradient: 0-5 min: 2% B, 5-65 min: 2-10% B. All solvents were HPLC grade. The absorbance of the eluting peaks was detected at 205 nm. Masses of peaks were detected with MSD API-ES Scan mode (m/z = 200-2,500) (Extended Data Table 12). For quantification of relative abundance of muropeptides, the area under the curve of assigned individual peak from chromatograms was integrated and percentage of individual peak was calculated relative to all assigned peaks.

### Western blot analysis

Western blot analysis was performed as described previously.^16,19^ Overnight VREfm cultures were sub-cultured in fresh BHI to OD_600_∼0.1 and were grown overnight at 37°C and shaking at 200 RPM. The cultures were centrifuged at 4,800 x *g* for 10 min, cell pellets and supernatants were separated. Cell pellets were washed with PBS and lysed in PBS with 0.1 mm glass beads (BioSpec) using FastPrep system (MP Biomedicals, settings: 6 m/s, 2 cycles, 45 s for each cycle). Proteins were quantified by Pierce™ BCA Protein Assay (ThermoFisher) and separated by SDS-PAGE on 4-20% Criterion TGX precast gels (Bio-Rad), then transferred to nitrocellulose membrane. The membrane was blocked in 0.1% TBST supplemented with 5% milk for 1 hour at room temperature with agitation, incubated with primary antibody (rabbit anti-SagA polyclonal sera, diluted 1:50000 in 0.1% TBST supplemented with 5% milk) overnight at 4°C with agitation. After 2 washes (2 min each) the membrane was incubated with secondary antibody (goat anti-rabbit antibody, HRP-conjugate, diluted 1:20,000 in 0.1% TBST supplemented with 5% milk) for 1 h at room temperature with agitation. Membranes were washed with 0.1% TBST three times (10 min each) at room temperature with agitation. Blots were developed using Clarity Western ECL substrate (Bio-Rad) and imaged using a ChemiDoc MP imaging system (Bio-Rad).

### Fluorescence microscopy

VREfm were grown to OD_600_ ∼ 0.4 with or without pghi-4 (50 μM) in BHI and sequentially labeled with 0.5 mM HADA (Tocris Bioscience) for 30 min and 1 μg/mL Vancomycin-BODIPY (Invitrogen) for 15 min at 220 RPM shaking, 37°C and protected from light. For Bocillin staining, VREfm were incubated with Bocillin (Invitrogen) at concentration 5 μM for 30 min at 220 RPM shaking, 37°C and protected from light. The cultures were centrifuged at 4,800 *× g* for 10 min. The cells were washed with PBS twice and fixed with 1% formaldehyde in PBS for 10 min, followed by washing and resuspending in PBS. To image cells, an aliquot of cell suspensions was transferred to the surface of a 2% (*w/v*) agarose pad prepared in PBS, covered with a glass coverslip, and imaged with fluorescence microscope (Nikon Ti2-E Inverted) using DAPI and GFP filters. For single-cell fluorescence analysis, fluorescence intensities from single cells normalized to similar region of interest were extracted from images. The images were processed using Icy open source imaging software^71^.

### Cryo-electron tomography

The protocol was adapted from ^19^ with some modifications. VREfm ERV165 strains or ERV165 strain treated with vancomycin (5 μg/mL), pghi-4 (50 μM) or in combination used in the cryo-ET experiments were grown overnight at 37 °C in BHI broth. Fresh cultures were prepared from a 1:50 dilution of the overnight culture and then grown at 37 °C to early log phase. The culture was centrifuged at 4,800 × *g* for 10 min. The pellet was resuspended with growth media supplemented with 5% glycerol to OD600 ∼ 3. Next, 5 µL of bacterial samples were deposited onto freshly glow-discharged (Pelco easiGlow; 25 s glow at 15mA) Quantifoil R2/1 copper 200 mesh grids for 1 min, back-side blotted with filter paper (Whatman Grade 1 filter paper), and frozen in liquid ethane using a gravity-driven homemade plunger apparatus (inside a 4°C cold room with a ≥95% relative humidity). The samples were frozen using a Vitrobot Mark IV (Thermo Fisher Scientific) in liquid ethane/propane mixture. The Vitrobot was set to 22°C at 90% humidity, and manually back-side blotted. The vitrified grids were later clipped with Cryo-FIB autogrids (Thermo Fisher Scientific) prior to milling.

Cryo-FIB milling was performed using an Aquilos2 dual-beam cryo-FIB/SEM instrument (Thermo Fisher Scientific). Vitrified samples were sputter-coated with metallic platinum for 15 s, followed by a 30 s coating with organometallic platinum, and then sputter-coated again with metallic platinum for 15 s to prevent drift during milling. Targets were selected and milled at an 8° angle using MAPS and AutoTEM software, respectively (Thermo Fisher Scientific). The milling template performed rough milling with a current of 0.30 nA, followed by medium milling at 0.1 nA. Thinning was conducted with a current of 50 pA. Automated milling produced lamellae with a thickness of ∼300 nm, which were then manually polished to <200 nm using a 30 pA current. A final 20 s metallic platinum coating was applied to facilitate bead-like fiducial inclusions for tilt series alignment.

Cryo-lamellae were imaged using a 300 keV Titan Krios microscope (Thermo Fisher Scientific) equipped with a field emission gun, an energy filter, and a direct electron detector (Gatan K3). An energy filter with a slit width of 20 eV was used during data acquisition. The SerialEM^72^ package with PACEtomo^73^ scripts was used to collect 35 image stacks at tilt angles ranging from +51° to −51° in 3° increments using a dose-symmetric scheme with a cumulative dose of ∼105 e⁻/Å². Data were collected at a magnification corresponding to 2.64 Å/pixel and a nominal defocus of ∼ −5 µm.

Image stacks containing 10 frames were motion-corrected using MotionCor2^74^, then assembled into drift-corrected stacks using IMOD^75^. These were aligned and reconstructed into tomograms using IMOD marker-based alignment. Tomograms were binned 4×, resulting in a final pixel size of 10.55 Å/pixel. Missing wedge artifacts were corrected using IsoNet^76^, a deep learning-based software. Segmentation of cell membranes was performed using MemBrain^77^. ColabSeg^78^ was used to isolate segmented membranes of interest. A MATLAB script^79^ was used to pick subtomograms with defined Euler angles perpendicular to the membrane, spaced ∼30 nm apart. Segmented membranes and subtomogram picks were validated in UCSF Chimera ^80^. Subtomogram averaging was performed using the Dynamo ^81^ software package. Subtomograms were extracted with a box size of 120 pixels. Five iterations of averaging with minimal translational and angular searches were conducted to generate averages. To generate density profiles, IMOD drawing tools were used to draw a line through the subtomogram average. Density values along the line were extracted, plotted and used for cell wall components assignment. Manual measurements of the cell envelope (CE) were performed on the apical and septal regions of cells using IMOD’s measurement tool. Workflow of subtomogram averaging and analysis is summarized in Extended Data Fig. 2.

### Macrophage infection

The protocol was adapted from ^52^. Macrophages (RAW264.7, ATCC TIB-71, or THP-1, ATCC TIB-202, differentiated with phorbol 12-myristate 13-acetate) were plated in 96-well plate in the growth media (DMEM or RPMI supplemented with 4.5 g/L glucose, 2 mM L-glutamine, 10% FBS) at the cell density 3×10^4^ cells/ well. Cells were left to adhere and grow overnight at 37°C in humidified atmosphere with 5% CO_2_. VREfm (ERV165 strain) were grown to OD∼0.6 in BHI by sub-culturing starting cultures grown overnight from a single colony. Bacterial cultures were centrifuged, and pellets were washed with sterile PBS twice, followed by resuspension in DMEM or RPMI with or without pghi-4 (25, 50 or 100 μM), vancomycin (100 μg/mL) or in combination. Macrophages were incubated with bacterial suspensions for 3 hours at 37°C, 5% CO_2_. Next, bacterial suspensions were removed, and macrophages were washed with PBS three times. Macrophages were incubated with 1% Triton X-100 (Sigma-Aldrich) in PBS for 5 mins at room temperature to lyse cells for colony-forming units (CFU) analysis. Lysates of macrophages were immediately plated on BHI agar, incubated overnight at 37°C and intracellular VREfm were enumerated.

### Mouse peritonitis infection

The protocol was adapted from^32^. Specific pathogen-free C57BL/6 (B6,000664) mice were obtained from Scripps Rodent Breeding. Mice were fed with gamma-irradiated chow (LabDiet, 5053) and sterile drinking water ad libitum. Animal care and experiments were conducted in accordance with NIH guidelines and approved by the Institutional Animal Care and Use Committee at Scripps Research. For evaluating vancomycin susceptibility of Δ*sagA in vivo,* bacterial cultures of VREfm strains (WT, Δ*sagA* or Δ*sagA*::*sagA*) grown in BHI (OD_600_ ∼ 0.6) were washed with PBS and resuspended in PBS at 5×10^8^ CFU/mL. Next, 6-8 weeks female mice were infected with 10^9^ VREfm CFU intraperitoneally. After 30 minutes, PBS or vancomycin (Van, 100 mg/kg) was administered subcutaneously. The second dose of treatment was administered 24-hour post-infection. VREfm infection was monitored by mice weight loss compared to initial weight (before infection) over 48 hours post-infection. Mice were euthanized once the weight reached 80% of the initial weight. Spleen and liver were collected, homogenized in sterile PBS for colony-forming units (CFU) analysis. Homogenates were plated on HiCrome™ selective *Enterococcus faecium* agar plates (HIMEDIA 1580) with *Enterococcus faecium* selective supplement (FD226, HIMEDIA), incubated at 37°C overnight and viable colonies were enumerated.

For evaluating combination therapy, bacterial cultures of VREfm (ERV165 strain) grown in BHI (OD_600_ ∼ 0.6) were washed with PBS and resuspended in PBS (0.25% carboxymethyl cellulose, CMC) at 5×10^8^ CFU/mL. Next, 6-8 weeks female mice were co-injected intraperitoneally with 0.2 mL of bacterial suspension (10^9^ CFU) and treatment: PBS (0.25% CMC) alone or supplemented with vancomycin (100 mg/kg), pghi-4 (25 mg/kg) or in combination. The second dose of treatment (PBS 0.25% CMC alone or supplemented with vancomycin (100 mg/kg), pghi-4 (25 mg/kg) or in combination was administered 24-hour post-infection. VREfm infection was monitored by mice weight loss compared to initial weight (before infection) over 48 hours post-infection. Mice were euthanized once the weight reached 80% of the initial weight. Spleen and liver were collected and processed as described above.

## Data availability

The mass spectrometry proteomics raw data have been deposited to the ProteomeXchange Consortium via the PRIDE partner repository with the dataset identifier PXD075040. All other data supporting the findings of this study are available within the article and its supplementary information files and from the corresponding author on reasonable request.

## Acknowledgements

This project was funded by the National Institutes of Health (NIH) R21AT012958 grant and Scripps Research start-up funds to H.C.H. J.E.M. thanks Cold Spring Harbor Laboratory for developmental funds from the NCI Cancer Center Support Grant (5P30CA045508), the Australian Research Council (ARC) for a Future Fellowship (FT170100156), and the F.M. Kirby Foundation. We thank Juliel Espinosa for providing constructs for recombinant expression of PGH2 and PGH3. We thank Francisco Martínez-Peña, Luke Lairson, Kayla Nutsch, Caroline Stanton and Michael Bollong for assisting with automated liquid handlers and EnVision plate readers. We thank Kathryn Spenser and Scott Henderson for assistance at the Scripps Microscopy Core, K. Barry Sharpless and the Hang laboratory members for their feedback.

## Author information

These authors contributed equally: Kyong T. Fam, Pavan Kumar Chodisetti

## Contributions

K.T.F. and H.C.H. conceived the project and planned initial experiments. P.K.C. generated ERV165 isogenic deletion and their derivative strains. K.T.F. and P.K.C. characterized all generated strains. C.J.S, S.K. generated screening library of sulfonyl fluorides. J.E.M and C.J.S. both designed, and C.J.S. generated the lead sulfonyl fluorides. Z.W., J.H. resynthesized pghi-4. K.T.F. developed assays, conducted screening and identified hits. K.T.F. performed all biochemical and microbiological characterizations of identified hits. K.T.F. and Y.X. conducted competitive cysteine-directed chemoproteomics. A.H.-H., M.H. performed computational docking. K.T.F, B.S. and D.P. performed cryoET experiments. K.T.F. and S.B. attempted X-ray crystallography studies. A.M.T. performed phylogenetic analysis. D.V.T. provided VREfm clinical isolates. H.C.H, J.E.M., D.W.W, D.P., I.A.W., S.F., B.F.C. supervised experiments. K.T.F. conducted all animal studies. K.T.F., H.C.H wrote the manuscript, which was edited by all the other authors. All authors approved the manuscript before submission.

## Ethics declarations

Authors declare no competing interests.

---

**Extended Data Fig. 1.**
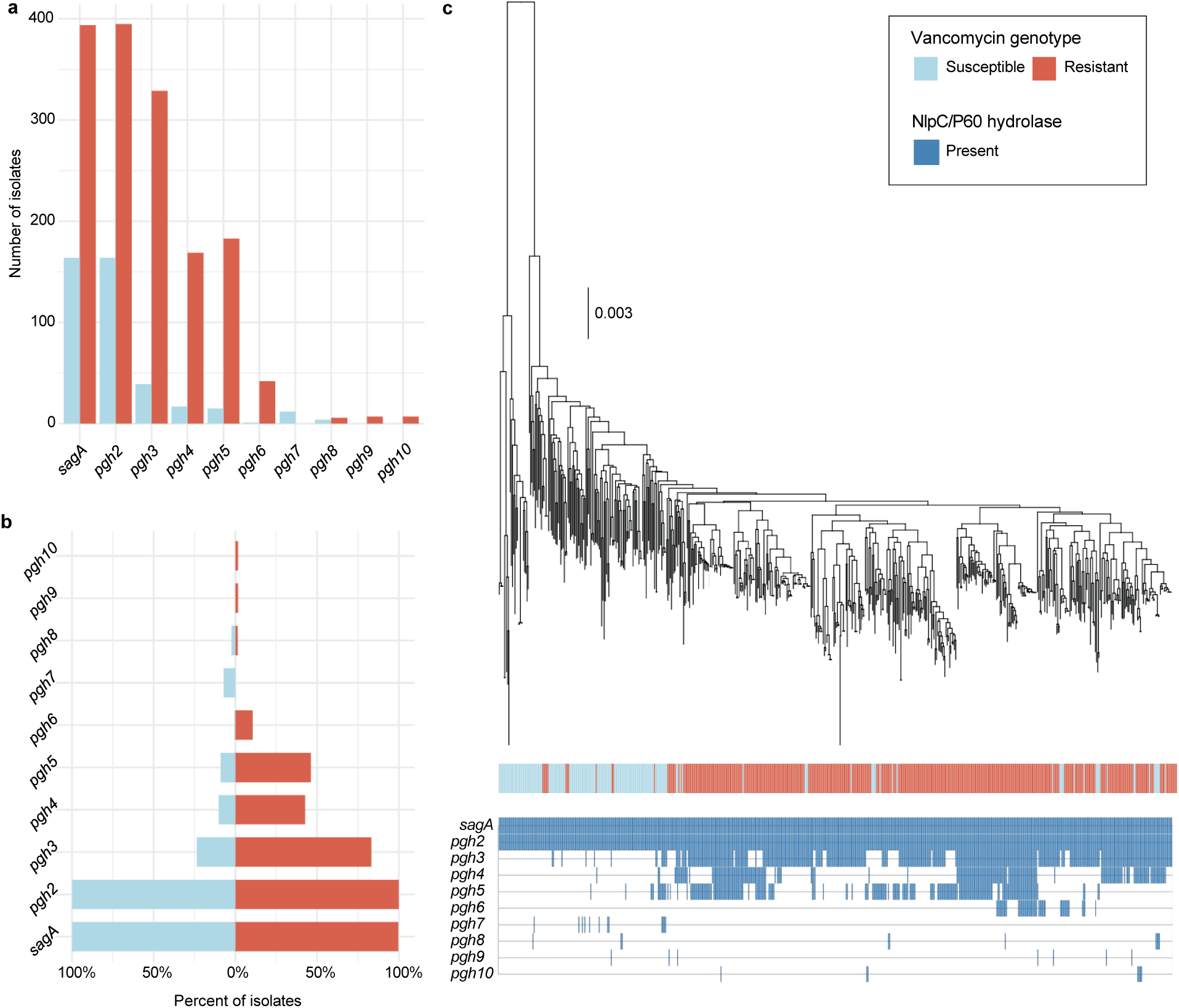
Phylogenetic analysis of NlpC/P60 hydrolases present in *E. faecium*. **a**, Identification of NlpC/P60 hydrolases present in *E. faecium,* labelled as *sagA* and <u>p</u>eptido<u>g</u>lycan <u>h</u>ydrolase (pgh) 2-10. **b**, Prevalence of the NlpC/P60 hydrolases in vancomycin-susceptible or - resistant *E. faecium*, shown as a percentage of isolates. Bars are colored by vancomycin genotype. **c**, Maximum-likelihood phylogenetic tree built of *E. faecium* (n=599) from the Panaroo core genome alignment. Overlaid is the vancomycin genotype (resistant defined as the presence of a *van* operon) and presence or absence of different NlpC/P60 hydrolases.

**Extended Data Fig. 2.**
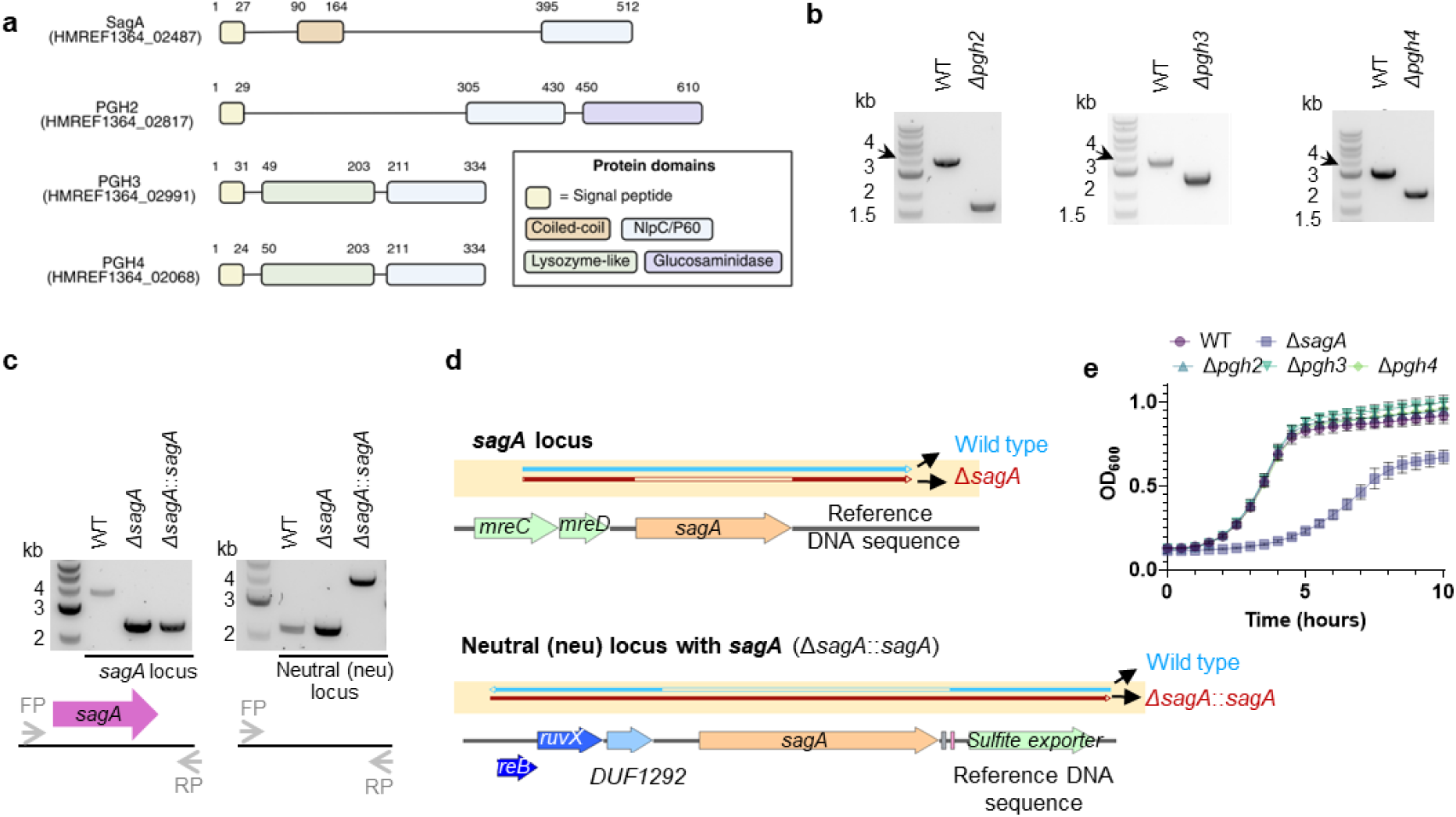
Characterization of *E. faecium* ERV165 NlpC/P60 hydrolases mutants. **a**, Comparison of the primary sequence homology and domain architecture of SagA orthologs in vancomycin-resistant *E. faecium* ERV165. Numbers above each bar are amino acid coordinates of the indicated domains. **b,** Colony PCR analysis of successful deletion of *phg2*, *pgh3*, *pgh4*. **c**, Colony PCR analysis using primers flanking the *sagA* and neutral loci to verify the *sagA* deletion mutant and chromosomal complementation strain, respectively. The left agarose gel shows successful deletion of *sagA* in both the mutant and the complementation strain. The right gel confirms *sagA* gene insertion at the neutral locus in the chromosomal complementation strain, but not in the wild-type (WT) or *sagA* mutant. **d**, Top: DNA sequencing reads from WT (blue) and *sagA* mutant (red) aligned to the ERV165 WT reference, confirming deletion. Bottom: DNA sequencing reads from WT (blue) and the chromosomal complementation strain (red) aligned to the complementation strain reference, confirming integration of *sagA* at a neutral locus. **e,** Growth curves of ERV165 strains in BHI. is mean ± S.D., n=3 biological replicates.

**Extended Data Fig. 3.**
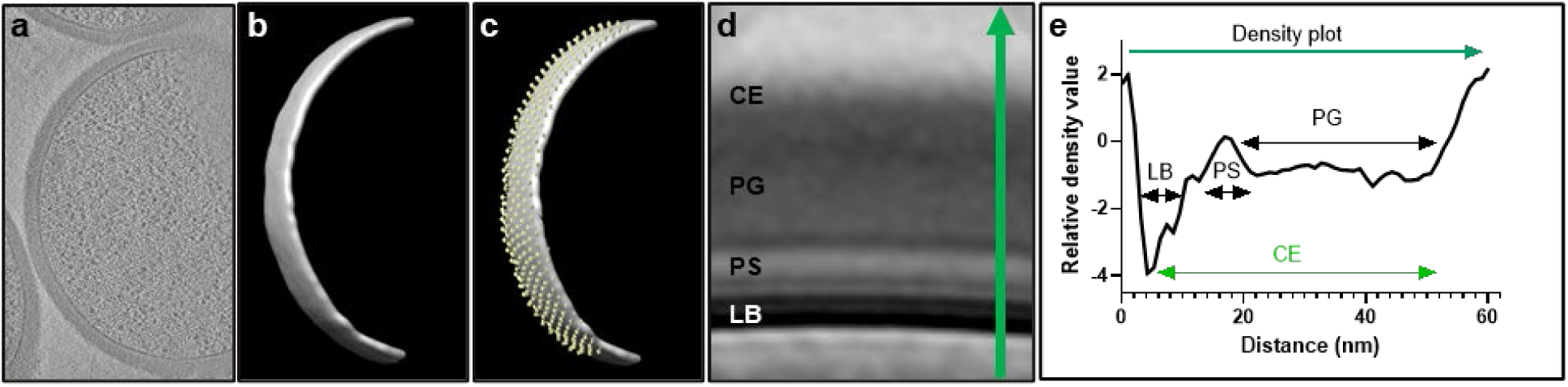
Subtomogram averaging and analysis workflow. **a**, Representative tomographic slice after reconstruction. **b**, 3D rendering of the segmented lipid bilayer. **c**, Subtomogram picks visualized on the segmentation (yellow); arrows indicate subtomogram positions and orientations. **d**, Averaged subtomogram map. Annotated layers include: CE, total cell envelope; PG, peptidoglycan (cell wall); PS, periplasmic space; LB, lipid bilayer. A green arrow is drawn across the cell envelope in the average map to collect relative density values. **e,** Plot of relative density values across the average map. Lower density values correspond to darker regions. Measurements were taken based on peak distances in the density profiles.

**Extended Data Fig. 4.**
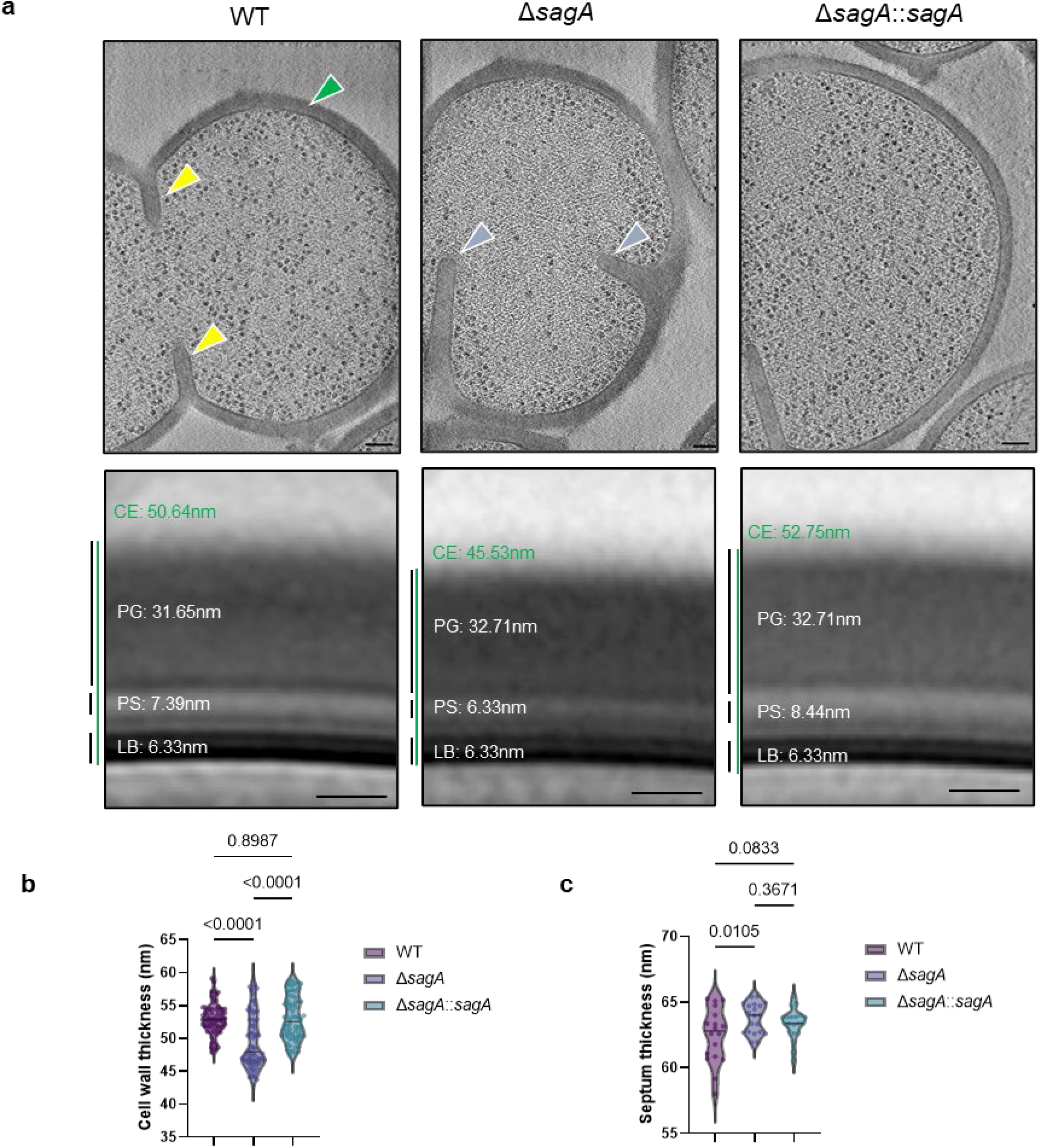
Ultrastructure of VREfm ERV165 strains. **a**, Top panel: representative cryo-electron tomographic (cryo-ET) slices of VREfm ERV165 strains. The cell wall is annotated with green arrows; normal septa are indicated with yellow arrows; and defective division septa are also marked with blue arrows. Images were acquired at a magnification corresponding to a pixel size of 2.638 Å (38,000×). Tomograms were 4× binned, resulting in a final pixel size of 10.55 Å. Scale bar, 100 nm. Bottom panel: subtomogram averages of the cell envelope of VREfm ERV165 strains were generated using Dynamo. Measured thicknesses of various envelope layers are indicated. Thickness measurements were derived from density plots of the subtomogram averages. Abbreviations: CE, total cell envelope; PG, cell wall; PS, periplasmic space; LB, lipid bilayer. Thickness measurements were obtained from density plots of the subtomogram averages (see Methods, Extended Data Fig. 2). Scale bar, 100 nm. **b**, Comparison of cell wall thickness. Data are shown as violin plots with dots representing individual data points and analyzed by one-way ANOVA with uncorrected Fisher’s LSD post-test, n=70. Grey horizontal lines represent median (WT: 52.93 nm, Δ*sagA*: 49.63 nm, Δ*sagA*::*sagA*: 53.01 nm). Grey dotted lines represent quartiles. **c**, Comparison of septum thickness. Data are shown as violin plots with dots representing individual data points and analyzed by one-way ANOVA with uncorrected Fisher’s LSD post-test, n=70. Grey horizontal lines represent median (WT: 62.48 nm, Δ*sagA*: 63.73 nm, Δ*sagA*::*sagA*: 63.40 nm).

**Extended Data Fig. 5.**
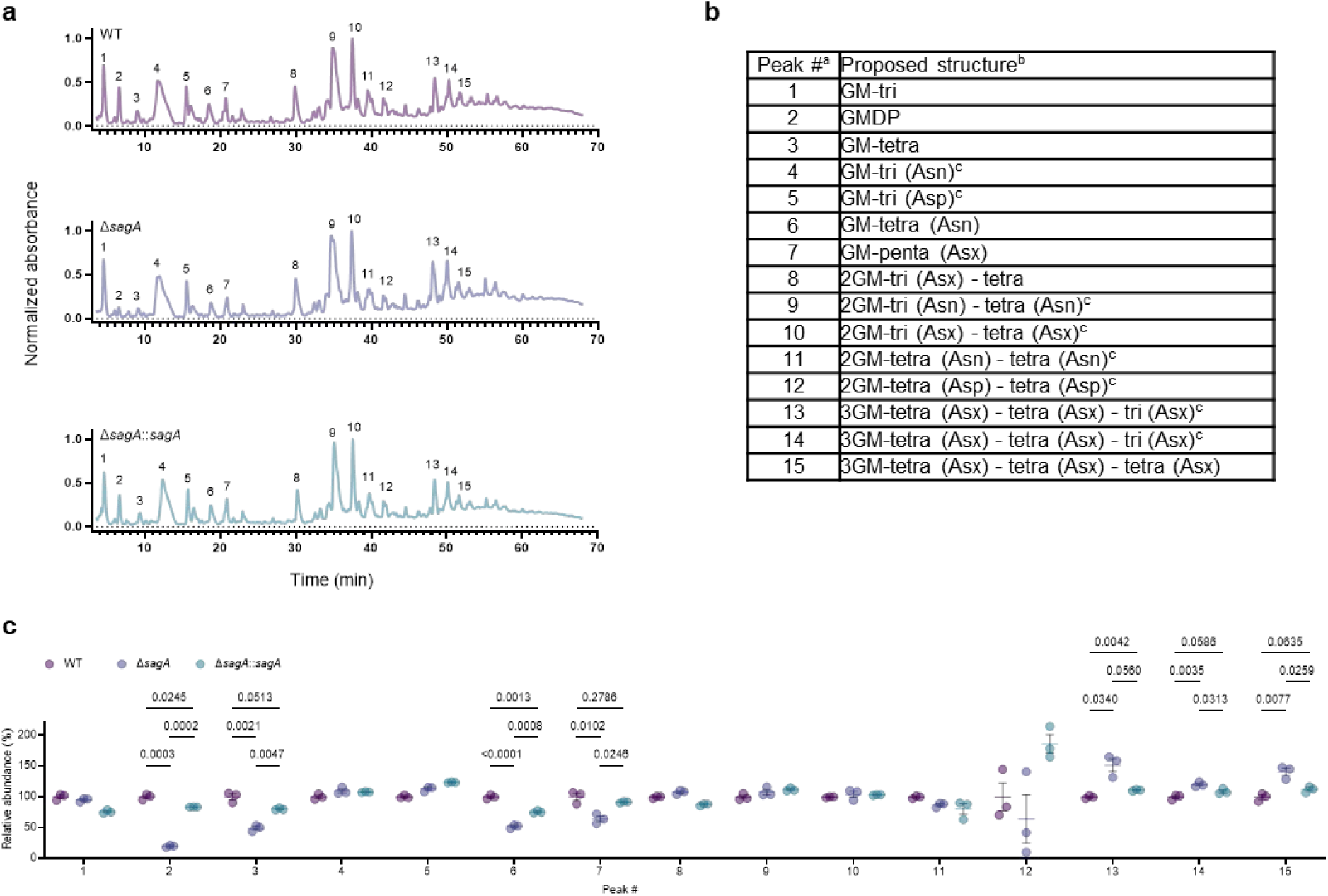
Inactivation of peptidoglycan remodeling in VREfm. **a**, Representative LC-MS chromatograms of mutanolysin-digested peptidoglycan isolated from sacculi of VREfm ERV165 strains. **b**, Composition of peptidoglycan isolated from VREfm sacculi. ^a^ Peak numbers refer to **a**. ^b^ GM, disaccharide (GlcNAc-MurNAc); 2 GM, disaccharide-disaccharide (GlcNAc-MurNAc-GlcNAc-MurNAc); 3 GM, disaccharide-disaccharide-disaccharide (GlcNAc-MurNAc-GlcNAc-MurNAc-GlcNAc-MurNAc); GM-Tri, disaccharide tripeptide (L-Ala-D-iGln-L-Lys); GM-Tetra, disaccharide tetrapeptide (L-Ala-D-iGln-L-Lys-D-Ala); GM-Penta, disaccharide pentapeptide (L-Ala-D-iGln-L-Lys-D-Ala-D-Ala). ^c^ The assignment of the amide and the hydroxyl functions to either peptide stem is arbitrary. Masses and retention time of peptidoglycan fragments are in Extended Data Table 12. **c**, Normalized abundance (relative to WT) of peptidoglycan fragments isolated from mutanolysin-digested sacculi of VREfm and analyzed by LC-MS. For **c** peak numbers indicate corresponding peptidoglycan fragment from LC-MS analysis listed in (**a**), data are mean ± S.D., n=3 biological replicates, analyzed with uncorrected Fisher’s LSD post-test, and p values of significant differences are indicated.

**Extended Data Fig. 6.**
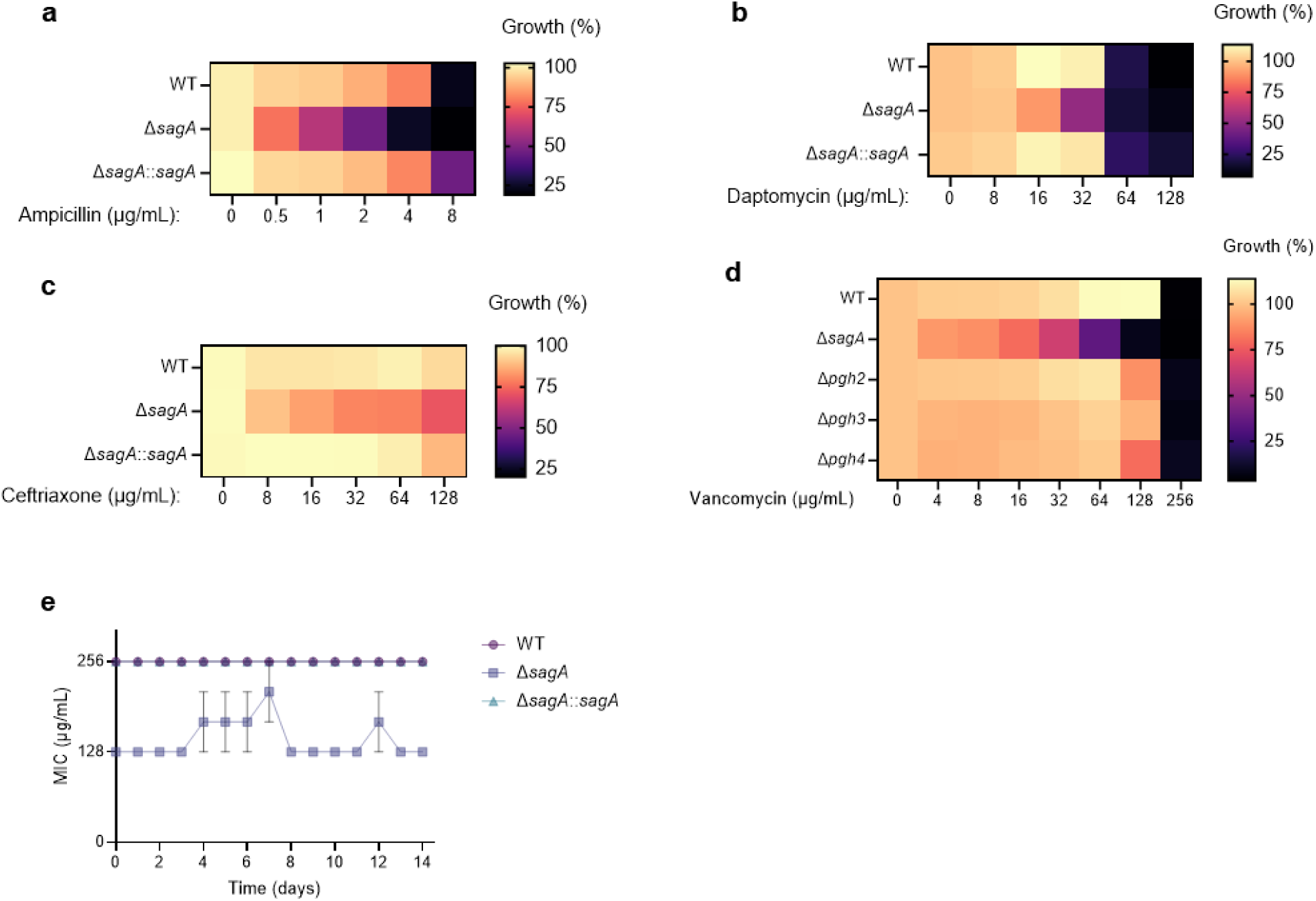
Antibiotic susceptibility of VREfm strains. **a**, Ampicillin susceptibility of ERV165 strains after 10 hours. **b**, Daptomycin susceptibility of ERV165 strains after 10 hours. **c**, Ceftriaxone susceptibility of ERV165 strains after 10 hours. **d**, Vancomycin susceptibility of ERV165 strains after 10 hours. For **a-d,** data are shown as a heat map of mean values, n=3 biological replicates. **e**, Vancomycin susceptibility of ERV165 strains after 14-day passaging under sub-MIC vancomycin concentration (50 µg/mL). Data are mean ± S.D., n=3 biological replicates.

**Extended Data Fig. 7.**
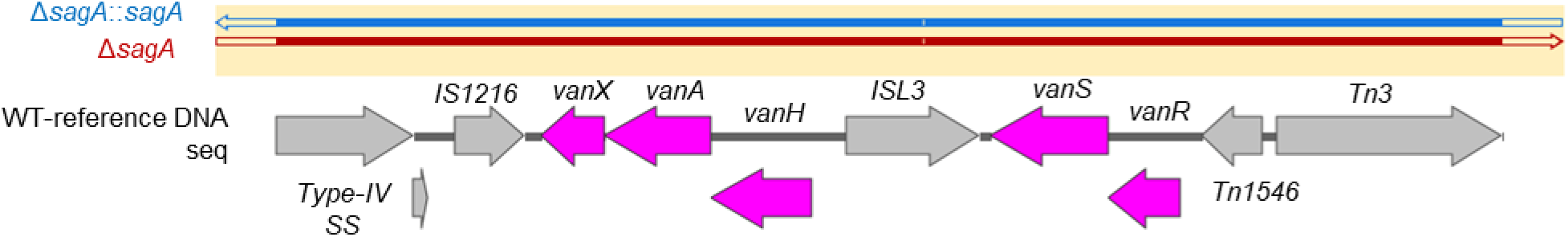
DNA sequencing reads from Δ*sagA*::*sagA* (blue) and *sagA* mutant (red) aligned to the ERV165 WT reference confirming intact *van* genes.

**Extended Data Fig. 8.**
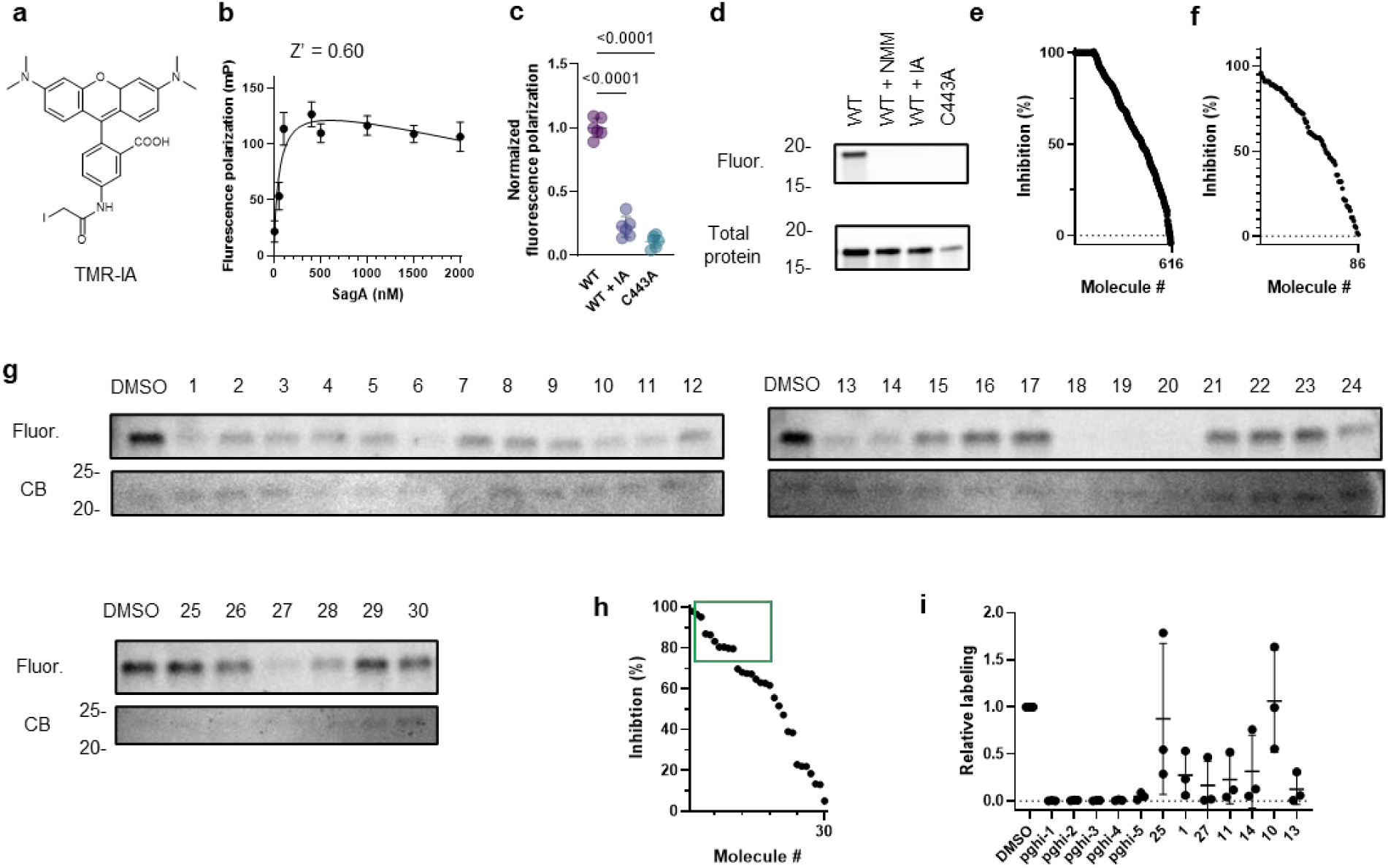
Validation of sulfonyl fluorides as SagA inhibitors. **a**, Structure of TMR-IA probe. **b**, Fluorescence polarization of TMR-IA (20 nM) with increasing concentration of recombinant SagA after 30 min incubation at room temperature in 384-well format. Z’ factor > 0.5 indicates the assay is suited for high-throughput screening. **c**, Fluorescence polarization (FP) assay for evaluating SagA (500 nM) cysteine reactivity with TMR-IA probe (20 nM) in 384-well format. FP signal was depleted when SagA was co-treated with non-specific cysteine-active iodoacetamide (IA, 1 mM) and TMR-IA (20 nM); or inactive SagA_C433A was treated with TMR-IA (20 nM). Data is mean ± S.D. and analyzed by one-way ANOVA, n=6 biological replicates. **d**, Gel of recombinant SagA (500 nM) treated with TMR-IA (50 nM) separated by SDS-PAGE and visualized by fluorescence. Fluorescent bands were absent when SagA was co-treated with non-specific cysteine-active *N*-methylmaleimide (NMM, 1mM), iodoacetamide (IA, 1 mM). TMR-IA treatment failed to produce fluorescent band of recombinant inactive mutant SagA_C443A (C433A). Stain-free imaging serves as total protein loading control. **e,** Plot of primary screen using competitive FP assay and TMR-IA probe. TMR-IA-treated inactive mutant SagA_C433A treated was used as lower limit of FP signal. Note: some molecules produce FP signal lower than SagA_C433A resulting in inhibition >100% that were assigned as 100% inhibition **f**, Plot of secondary screen of 86 molecules using competitive gel-based assay and TMR-IA probe. Inhibition (%) was calculated as TMR-IA competition relative to DMSO control, n=1. **g**, Gels of validation of 30 sulfonyl fluorides. Recombinant SagA (0.5 µM) treated with sulfonyl fluorides (50 µM), followed by TMR-IA (50 nM), separated by SDS-PAGE and visualized by fluorescence. Coomassie blue (CB) staining served as total protein control. **h**, Plot of validation of 30 molecules using competitive gel-based assay and TMR-IA probe. For **f**, **h** inhibition (%) was calculated as competition with TMR-IA relative to DMSO control, n=1. **i**, Plot of revalidation of 12 most active sulfonyl fluorides (green box in **e**) using competitive gel-based assay. Data are mean labeling values relative to DMSO ± S.D, n=3 biological replicates.

**Extended Data Fig. 9.**
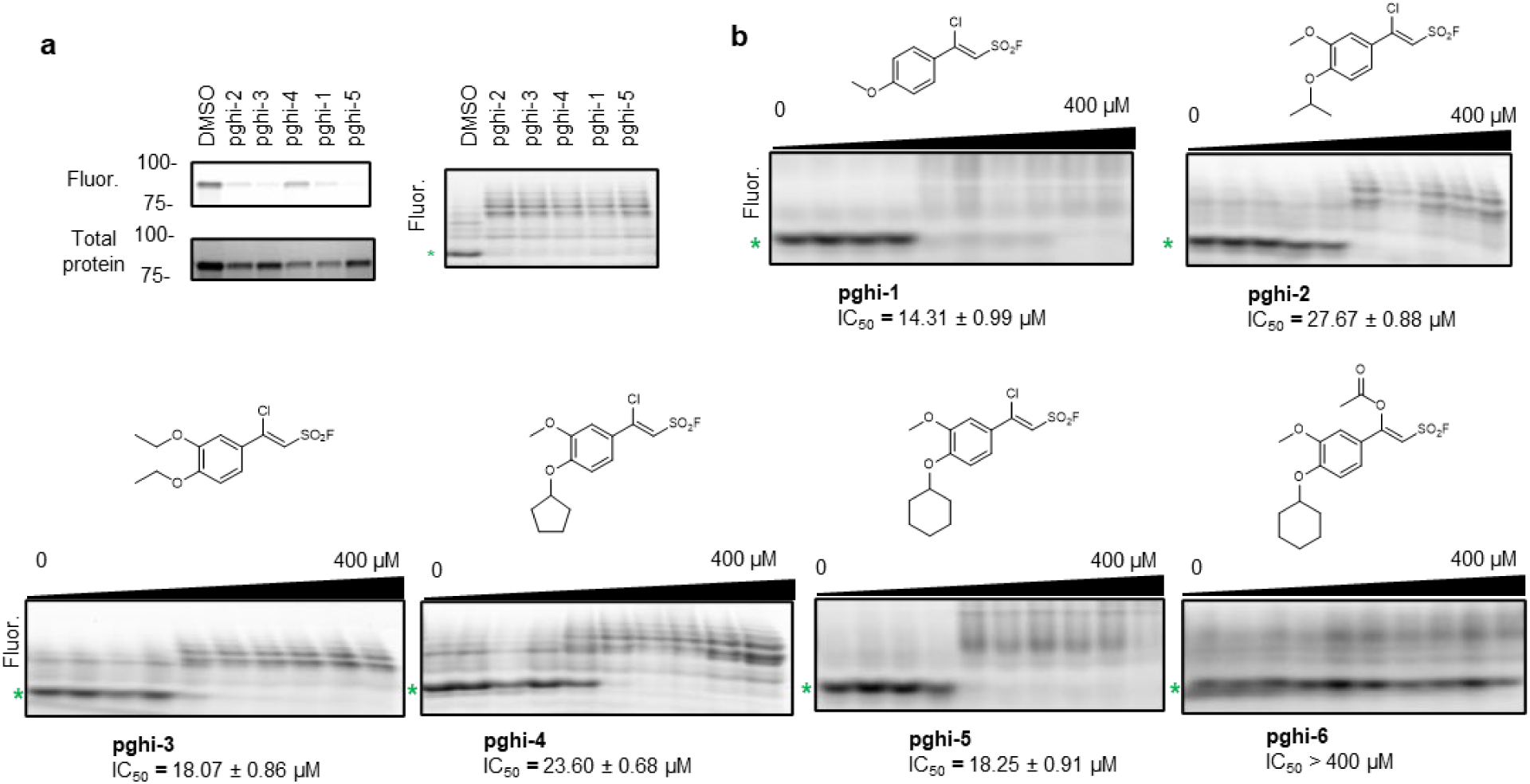
Structure-activity relationship of SagA inhibitors. **a**, Activity of top 5 sulfonyl fluorides (50 µM) against of recombinant SagA (10 µM) profiled in competitive gel-based (on the left) and peptidoglycan hydrolase activity (on the right) assays. **b**, Native gel profiling assay of hydrolase activity of recombinant SagA (10 µM) treated with sulfonyl fluorides (0 - 400 µM) and mutanolysin-digested peptidoglycan from *E. faecium* (100 µg) followed by peptidoglycan labeling with ANTS (0.2 M), separated by gel electrophoresis. For **a** and **b,** green asterisk indicates the main enzymatic product GlcNAc-MurNAc-L-Ala-D-isoGln (GlcNAc-MDP)

**Extended Data Fig. 10.**
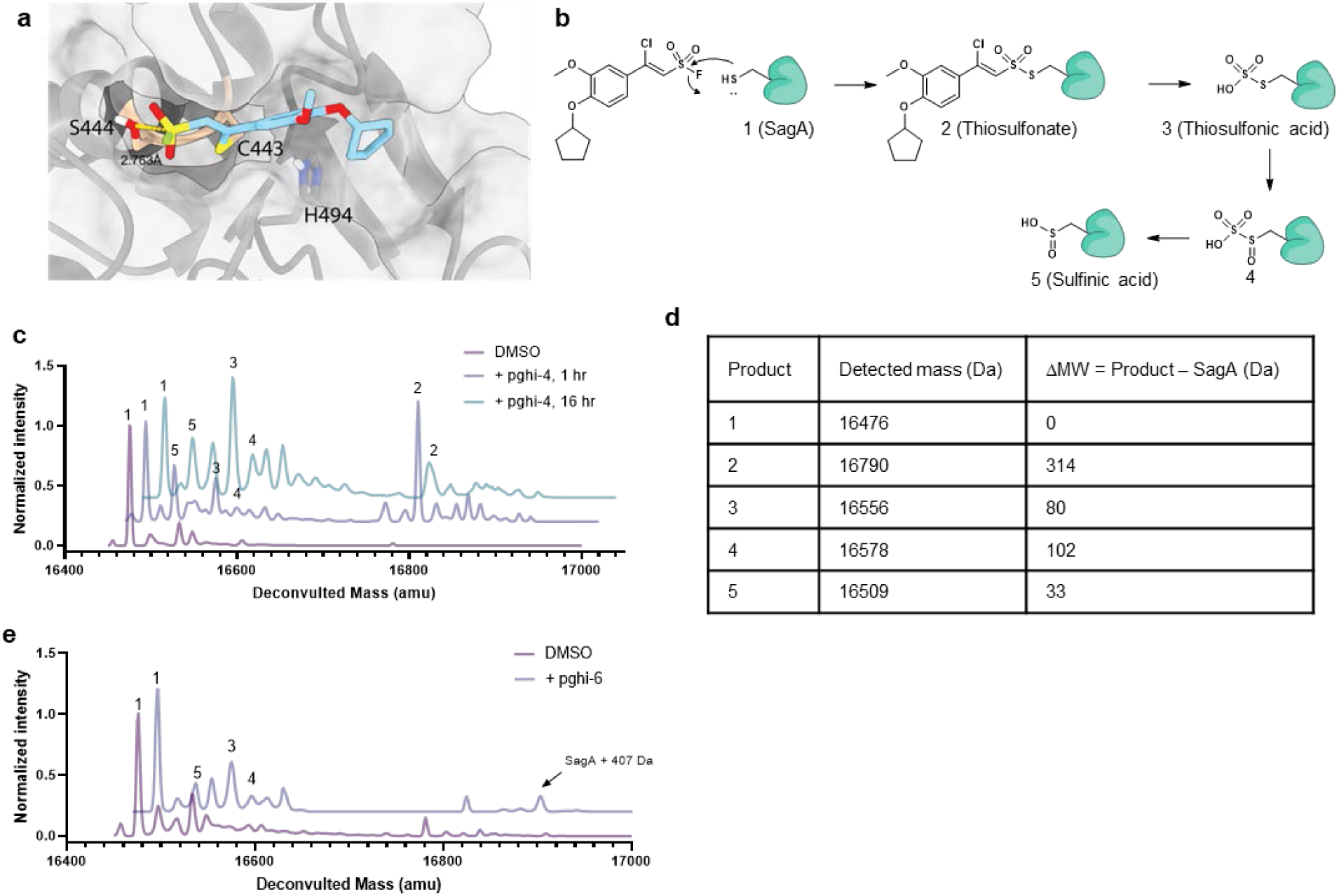
Sulfonyl fluorides are covalent inhibitors of SagA. **a,** Covalent docking model of pghi-4 to cysteine (C433) of SagA as adduct of vinyl chloride activity. **b,** Proposed structures of pghi-4-SagA adduct and products of hydrolysis detected by intact protein analysis (ESI). **c,** Intact protein analysis (ESI) of SagA (10 µM) treated with pghi-4 (50 µM) for 1 or 16 hours at room temperature. Numbers indicate products from scheme (**b**). **d,** Table of detected masses of SagA (10 µM) treated with pghi-4 (50 µM) and calculated differences in masses compared to unmodified SagA (ΔMW). **e,** Intact protein analysis (ESI) of SagA (10 µM) treated with inactive pghi-6 (50 µM) for 1 hr. A minor formation of pghi-6-SagA adduct (SagA+407 Da) and products of hydrolysis were detected: 3, 4 and 5. Numbers indicate products from scheme (**b**).

**Extended Data Fig. 11.**
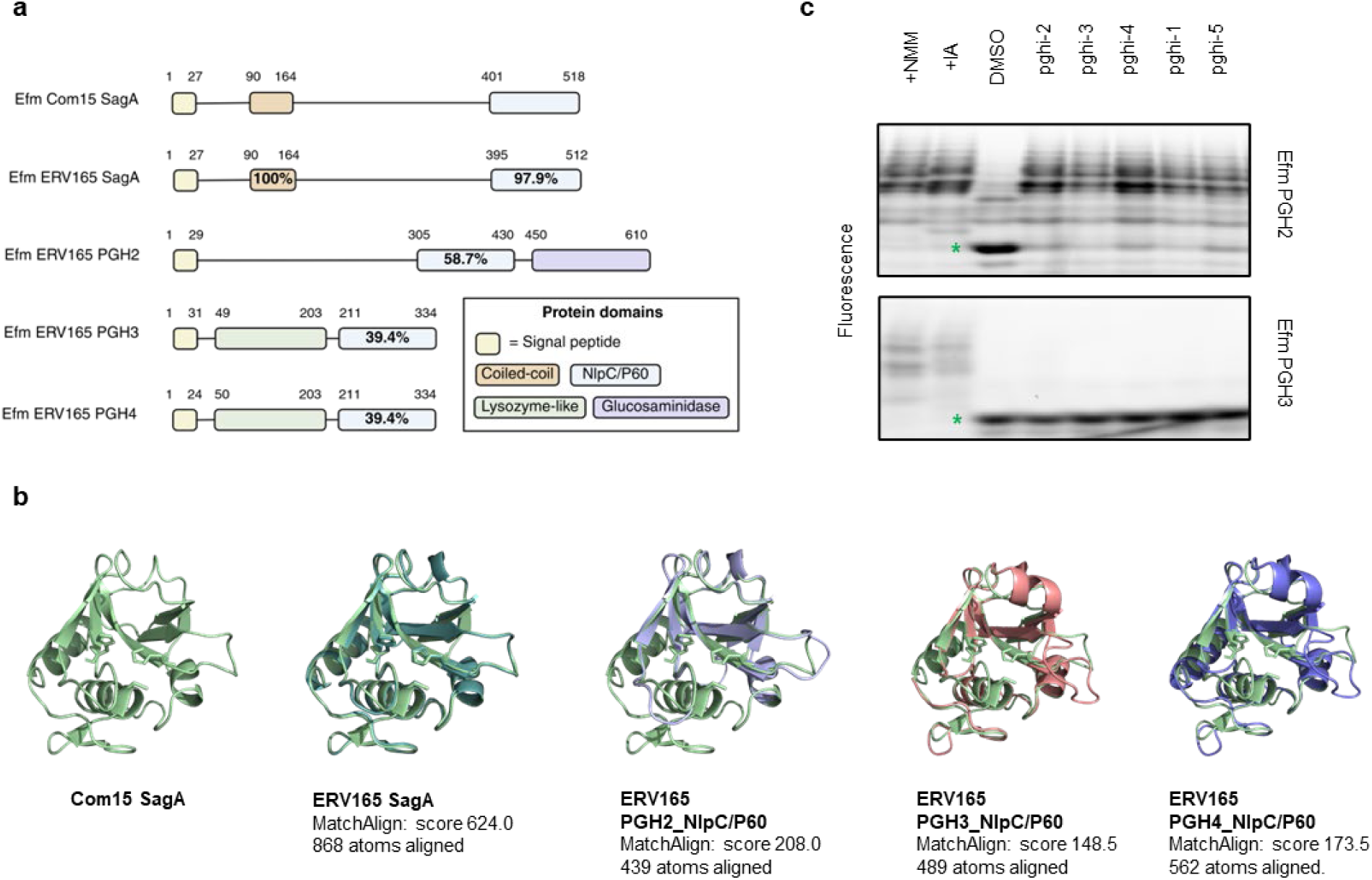
*Enterococcus* NlpC/P60 hydrolases. **a**, Comparison of primary sequence homology and domain architecture of SagA orthologs and SagA-like proteins from commensal and vancomycin-resistant *E.* faecium. Numbers above each bar are amino acid coordinates of the indicated domains, and percentages are amino acid sequence identity relative to Efm Com15 SagA. **b**, 3D homology modeling of SagA orthologs from commensal and vancomycin-resistant *E. faecium*. Predicted structure models for catalytic domains were generated in AlphaFold3. Predicted structures were aligned with published structure of SagA (PDB: 6B8C) in PyMOL. The amino acid residues of the SagA catalytic triad are shown using stick models in light green. MatchAligh score and number of atoms aligned are analyzed in PyMol and indicated. **c**, Native gel profiling assay of hydrolase activity of representative recombinant SagA orthologs containing NlpC/P60 domain from *E. faecium* and *E. faecalis* treated with sulfonyl fluorides. Green star indicates a major enzymatic product GlcNAc-MDP.

**Extended Data Fig. 12.**
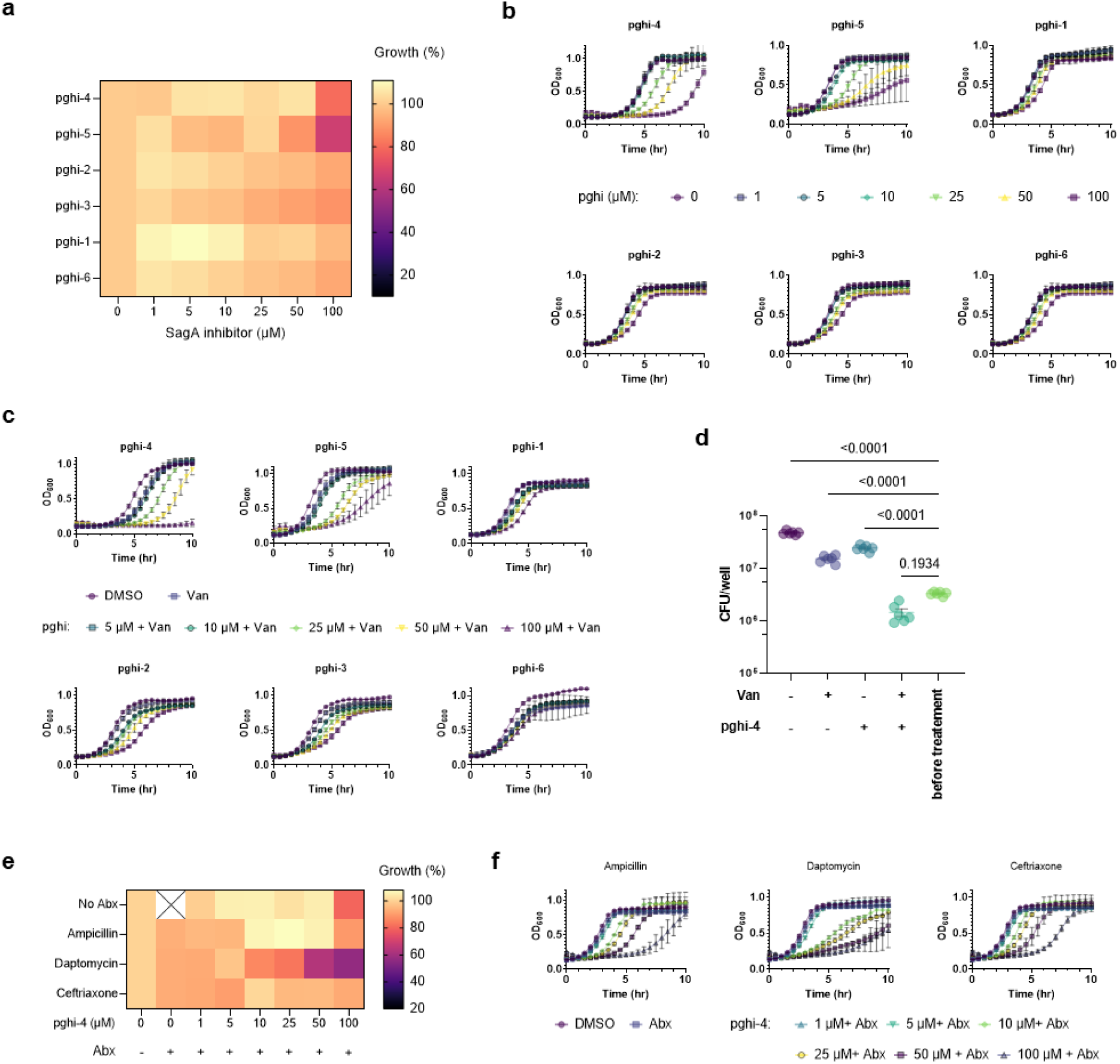
Antibacterial activity of SagA inhibitors. **a**, Heat map of VREfm ERV165 growth (%) in the presence of sulfonyl fluorides after 10 hours. **b,** Growth curves VREfm ERV165 in the presence of sulfonyl fluorides in BHI. **c,** Growth curves of VREfm ERV165 in the presence of pghi-4 and subinhibitory dose of vancomycin (Van, 5 μg/mL) in BHI. **d,** Colony-forming unit (CFU) analysis of ERV165 after treatment with pghi-4 (100 μM), vancomycin (Van, 100 μg/mL) or in combination for 18 hours. VREfm before treatment were used for comparison. Data is mean ± S.E.M. and analyzed by one-way ANOVA with uncorrected Fisher’s LSD post-test, n=3 biological replicates. **e,** Heat map of VREfm (ERV165) growth (%) in the presence of pghi-4 ± subinhibitory dose of ampicillin (1 µg/mL), daptomycin (4 µg/mL) or ceftriaxone (1 µg/mL) in BHI after 10 hours. Data is mean value, n=3 biological replicates. Cross indicates no data. **f,** Growth curves VREfm ERV165 in the presence of pghi-4 ± subinhibitory dose of ampicillin (1 µg/mL), daptomycin (4 µg/mL) or ceftriaxone (1 µg/mL) in BHI. For **a**, **e,** data are a heat map of mean values, n=3 biological replicates. For **b, c** and **f,** data are mean ± S.D., n=3 biological replicates.

**Extended Data Fig. 13.**
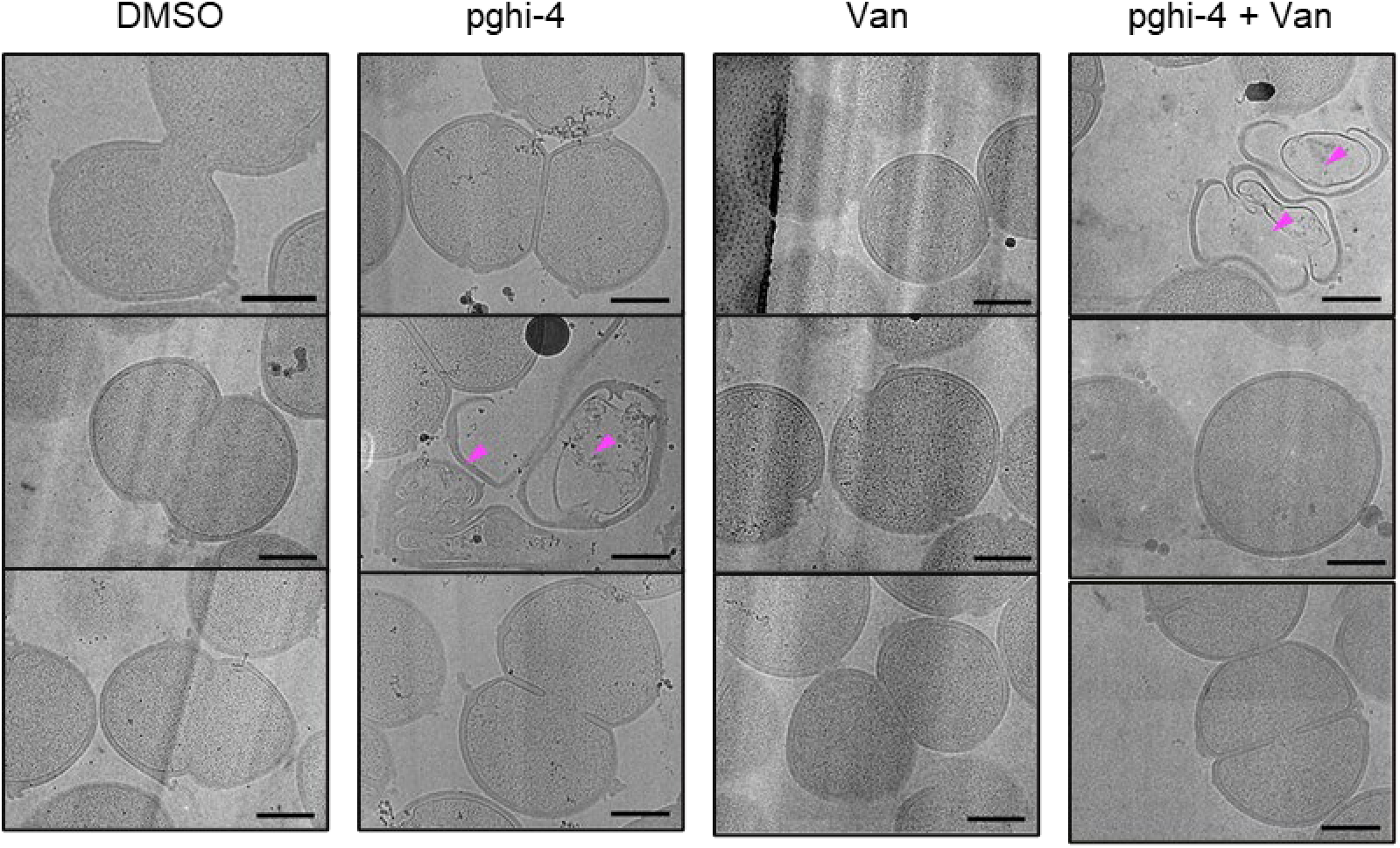
SagA inhibitor in combination with vancomycin alters VREfm cell morphology. Representative low-magnification (3600×) electron microscopy images from cryo-lamellae of VREfm in the presence of vancomycin (Van, 5 µg/mL) ± pghi-4 (50 µM). Each column contains three images of the same condition. Dead cells with undegraded peptidoglycan are indicated with magenta arrows. Scale bars, 500 nm.

**Extended Data Fig. 14.**
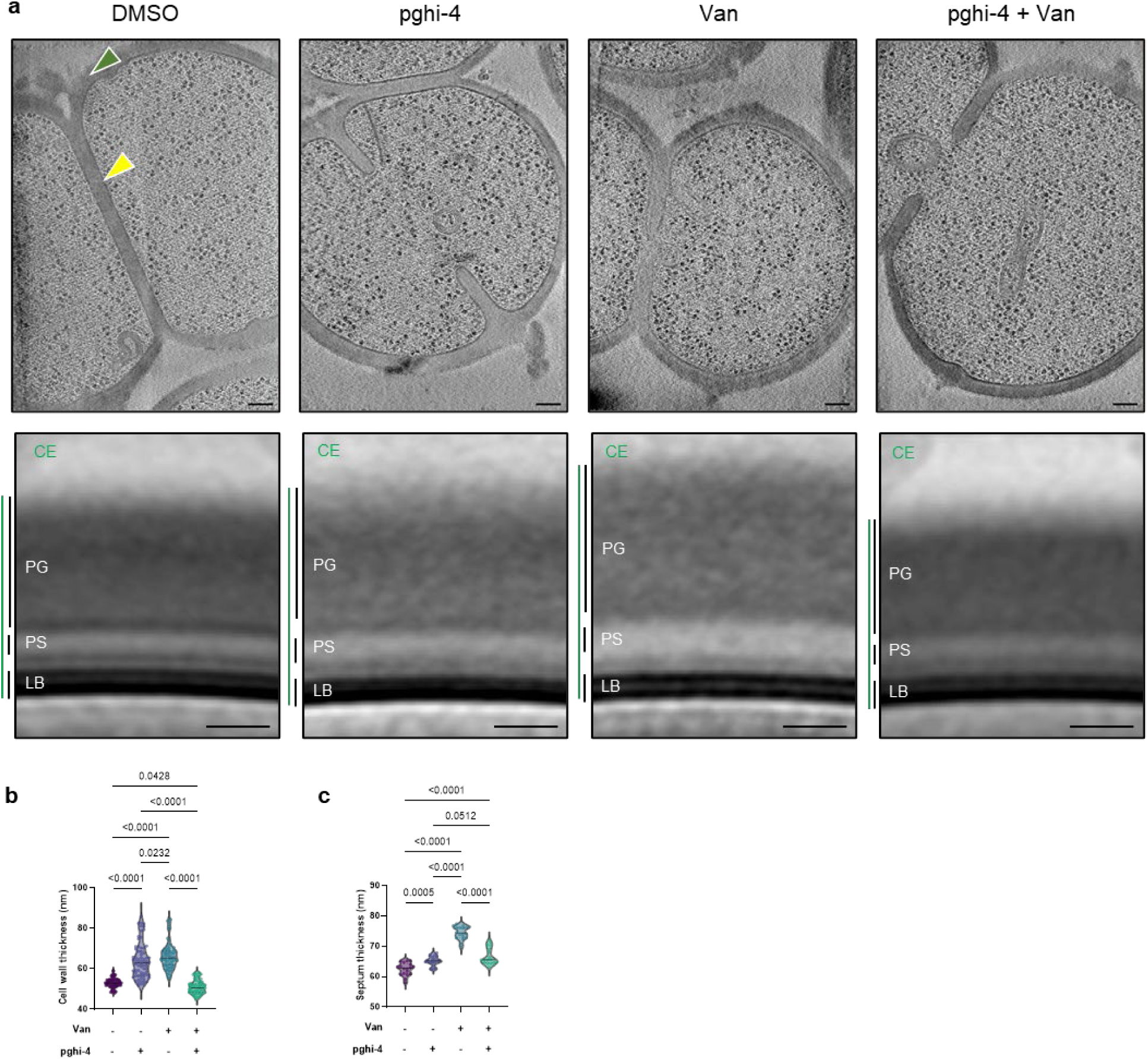
Ultrastructure of VREfm treated with SagA inhibitor pghi-4 ± vancomycin. **a,** Top: representative cryo-electron tomography (cryo-ET) images of VREfm ± vancomycin (Van, 5 µg/mL) ± pghi-4 (50 µM) in BHI are shown in the top row. The cell wall is annotated with green arrows; normal septa are indicated with yellow arrows. Bottom: subtomogram averages of the cell envelope were generated using Dynamo. Measured thicknesses of various envelope layers are indicated. Thickness measurements were derived from density plots of the subtomogram averages. Abbreviations: CE, total cell envelope; PG, cell wall; PS, periplasmic space; LB, lipid bilayer. Thickness measurements were obtained from density plots of the subtomogram averages (see Methods). Scale bar, 100 nm. Note: images and analysis of “DMSO” are the same as “WT” in Extended Data Fig. 3a. **b,** Comparison of cell wall thickness. Data is violin plot with dots representing individual data points and analyzed by one-way ANOVA with uncorrected Fisher’s LSD post-test, n=70. Grey horizontal lines represent median (DMSO: 52.93 nm, pghi-4: 63.96 nm, Van: 66.20 nm, Van+pghi-4: 50.93 nm). **c,** Comparison of septum thickness. Data are shown as violin plots with dots representing individual data points and analyzed by one-way ANOVA with uncorrected Fisher’s LSD post-test, n=70. Grey horizontal lines represent median (DMSO: 62.48 nm, pghi-4: 65.24 nm, Van: 74.45 nm, Van+pghi-4: 66.56 nm). Note: images and data for DMSO are same as WT in Extended Data Fig. 3; workflow of the analysis is in Extended Data Fig. 2.

**Extended Data Fig. 15.**
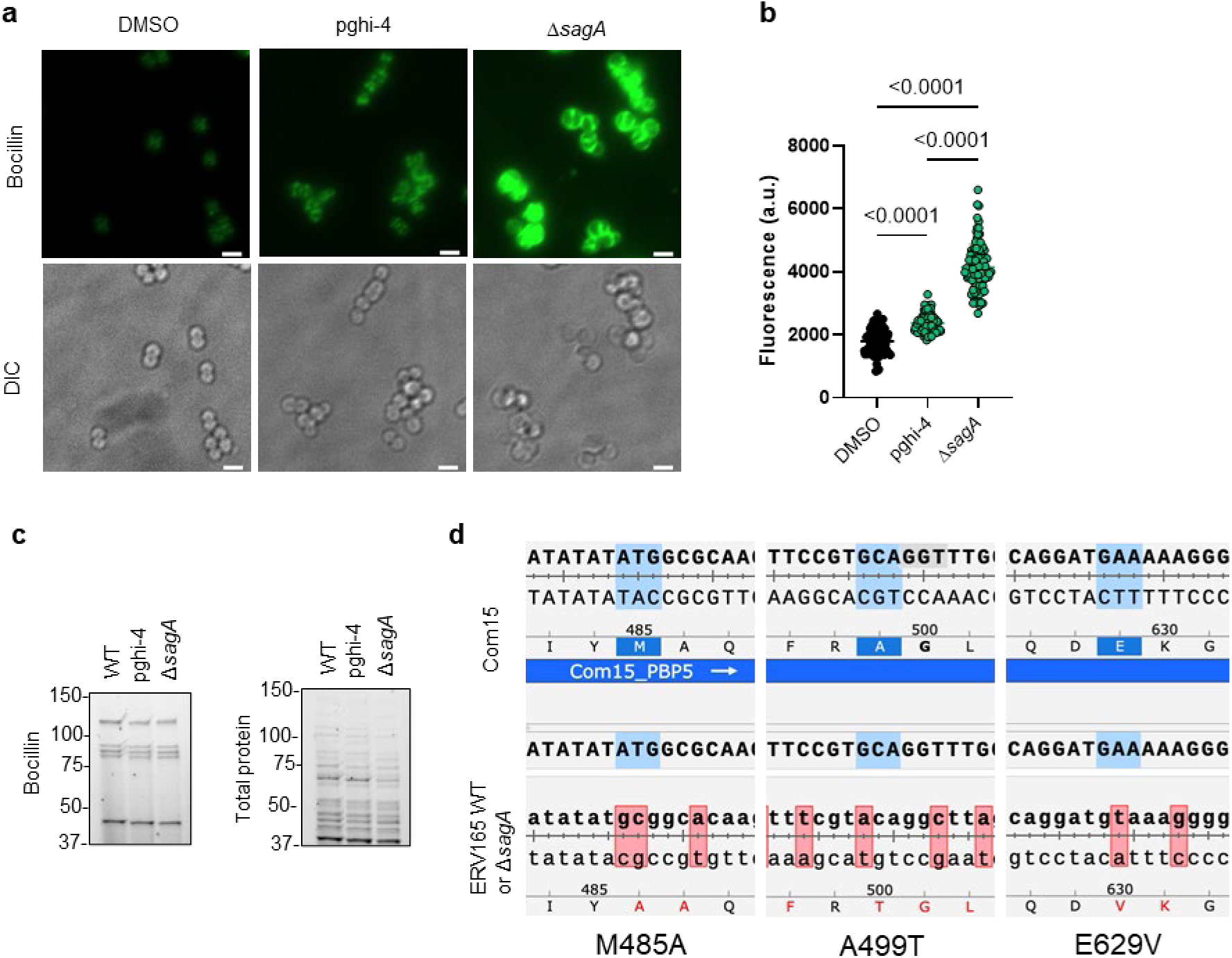
Bocillin labeling in VREfm. **a,** Fluorescence microscopy imaging of VREfm ± pghi-4 (50 µM) or Δ*sagA* stained with Bocillin (5 µM). Scale bar, 2 µm. **b**, Fluorescence of Bocillin-stained VREfm ± pghi-4 (50 µM) or Δ*sagA*. Each dot is individual cell, n=100. **c**, Left: cell lysates from VREfm ± pghi-4 (50 µM) or Δ*sagA* stained with Bocillin (5 µM) analyzed by SDS-PAGE separation and visualized by fluorescence. Right: total protein loading was visualized by Stain-free imaging and serves as protein loading. **d**, DNA sequencing reads from and VREfm ERV165 strains (WT or Δ*sagA*) aligned to the commensal *E.* faecium Com15 strain showing mutations in PBP5: M4585A, A499T and E629V.

**Extended Data Fig. 16.**
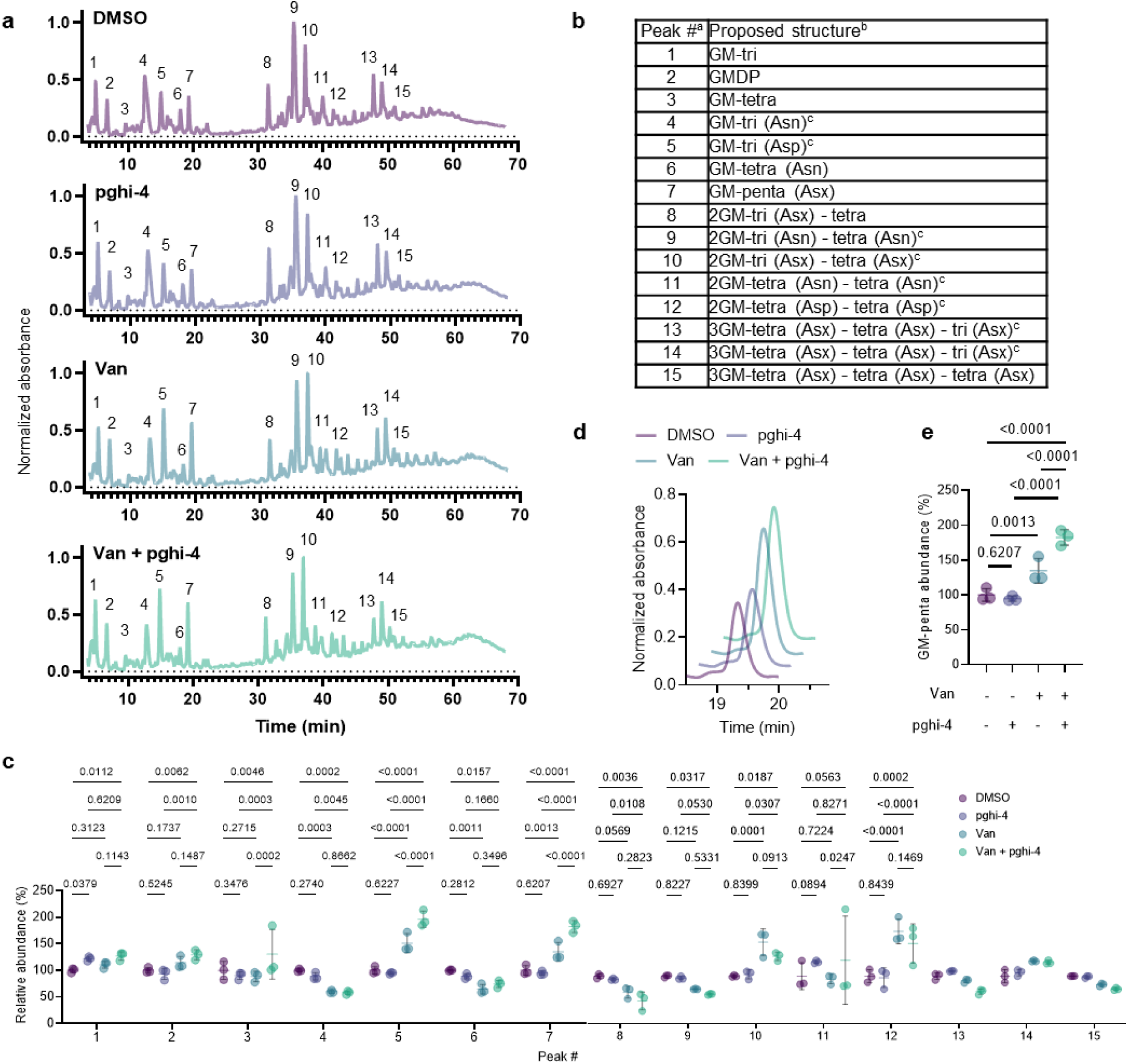
Inactivation of peptidoglycan remodeling in VREfm. **a,** Representative LC-MS chromatograms of mutanolysin-digested peptidoglycan isolated from sacculi of VREfm after treatment with pghi-4 (50 µM), vancomycin (5 µg/mL) alone, or in combination. **b,** Composition of peptidoglycan isolated from VREfm sacculi. ^a^ Peak numbers refer to (**a**). ^b^ GM, disaccharide (GlcNAc-MurNAc); 2 GM, disaccharide-disaccharide (GlcNAc-MurNAc-GlcNAc-MurNAc); 3 GM, disaccharide-disaccharide-disaccharide (GlcNAc-MurNAc-GlcNAc-MurNAc-GlcNAc-MurNAc); GM-Tri, disaccharide tripeptide (L-Ala-D-iGln-L-Lys); GM-Tetra, disaccharide tetrapeptide (L-Ala-D-iGln-L-Lys-D-Ala); GM-Penta, disaccharide pentapeptide (L-Ala-D-iGln-L-Lys-D-Ala -D-Ala). ^c^ The assignment of the amide and the hydroxyl functions to either peptide stem is arbitrary. Masses and retention time of peptidoglycan fragments are in Extended Data Table 11. **c,** Normalized abundance (relative to DMSO) of peptidoglycan fragments isolated from mutanolysin-digested sacculi of VREfm after treatment with pghi-4 (50 µM), vancomycin (5 µg/mL) alone, or in combination and analyzed by LC-MS. For **c** peak numbers indicate corresponding peptidoglycan fragment from LC-MS analysis listed in (**a**), data is heat-map of mean values, n=3 biological replicates. **d,** Abundance of GM-penta muropeptide (peak 7 in LC-MS chromatogram (**a**) shown as representative extracted LC-MS chromatogram of mutanolysin-digested peptidoglycan isolated from sacculi of VREfm after treatment with pghi-4 (50 µM), vancomycin (5 µg/mL) alone, or in combination. **e,** Normalized abundance of GM-penta muropeptide (relative to DMSO) within peptidoglycan composition of VREfm ± pghi-4 (50 µM) ± vancomycin (5 µg/mL) from LC-MS chromatograms (**a**) and (**c**). For **c**, **e**, data are mean ± S.D., n=3 biological replicates, analyzed with one-way ANOVA and uncorrected Fisher’s LSD post-test, p values of significant differences are indicated.

**Extended Data Fig. 17.**
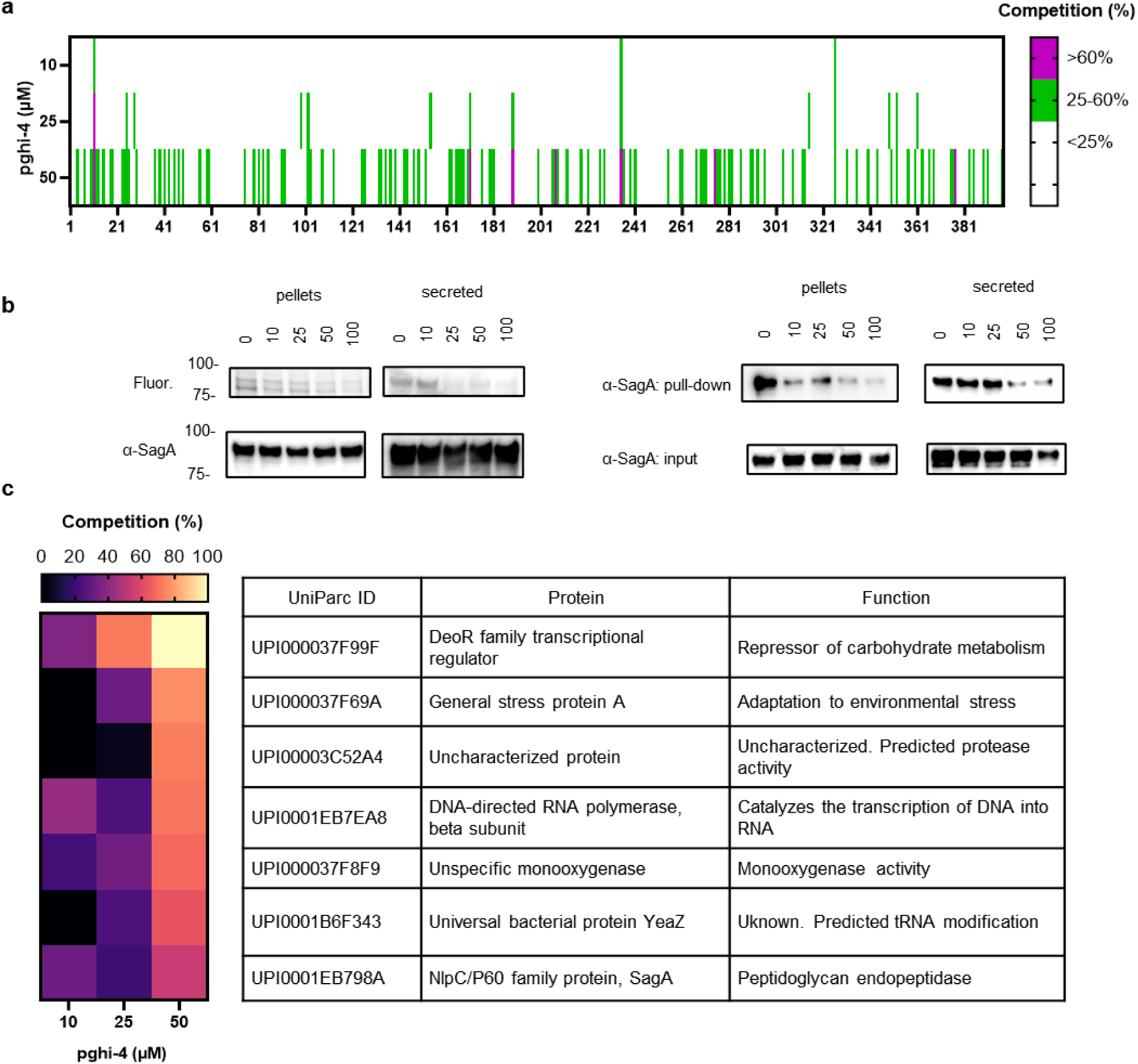
Cysteine-directed chemoproteomic analysis. **a,** Competitive cysteine-directed chemoproteomic analysis of VREfm (ERV165) proteome after treatment with pghi-4. Data are clustering as percentage of competition for cysteine occupancy relative to desthiobiotin-iocoacetamide activity. **b,** Pghi-4-treated ERV165 cell lysates and secreted proteins were reacted with iodoacetamide-alkyne probe. Left: clicked to rhodamine-azide, followed by SDS-PAGE separation and visualized by fluorescence. α-SagA western blot analysis served as protein loading control. Right: clicked to biotin-azide, followed by enrichment, SDS-PAGE separation and analyzed by α-SagA western blot. α-SagA western blot analysis of input (2.5%) served as protein loading control. **c,** Heat-map of activity of top 7 cysteine-directed protein targets of pghi-4 with their functions in ERV165. Data are representative biological replicates of two independent datasets.

**Extended Data Fig. 18.**
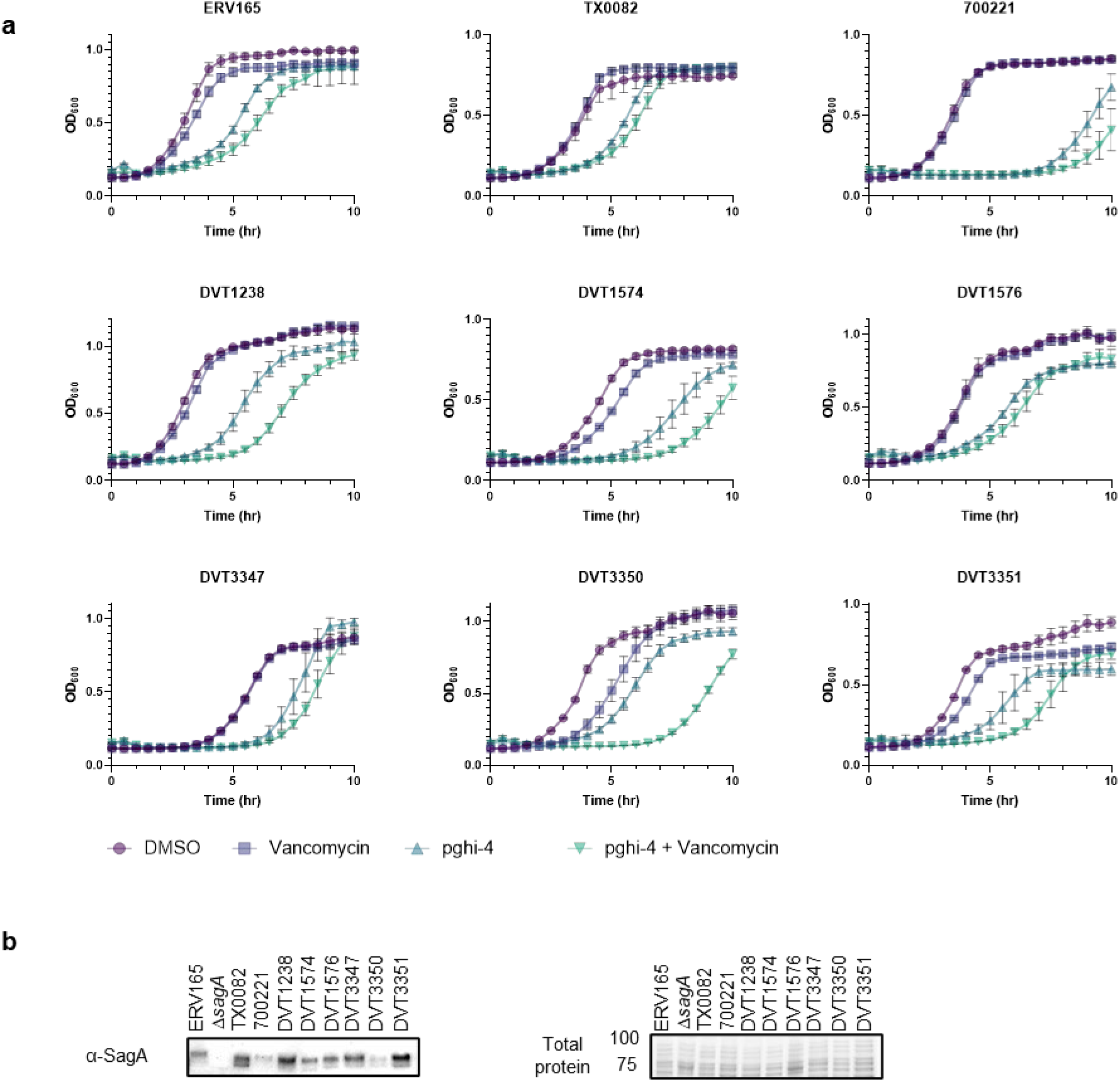
Therapeutic efficacy of pghi-4 as vancomycin adjuvant *in vitro*. **a,** Growth curves of VREfm clinical isolates ± vancomycin (Van, 5 μg/mL) ± pghi-4 (50 μM) in BHI. Data is mean ± S.D., n=3 biological replicates. **b,** Left: ɑ-SagA western blot of VREfm clinical isolates cell lysate. Right: Stain-free imaging served as protein loading control, numbers indicate molecular weight (kDa).

**Extended Data Fig. 19.**
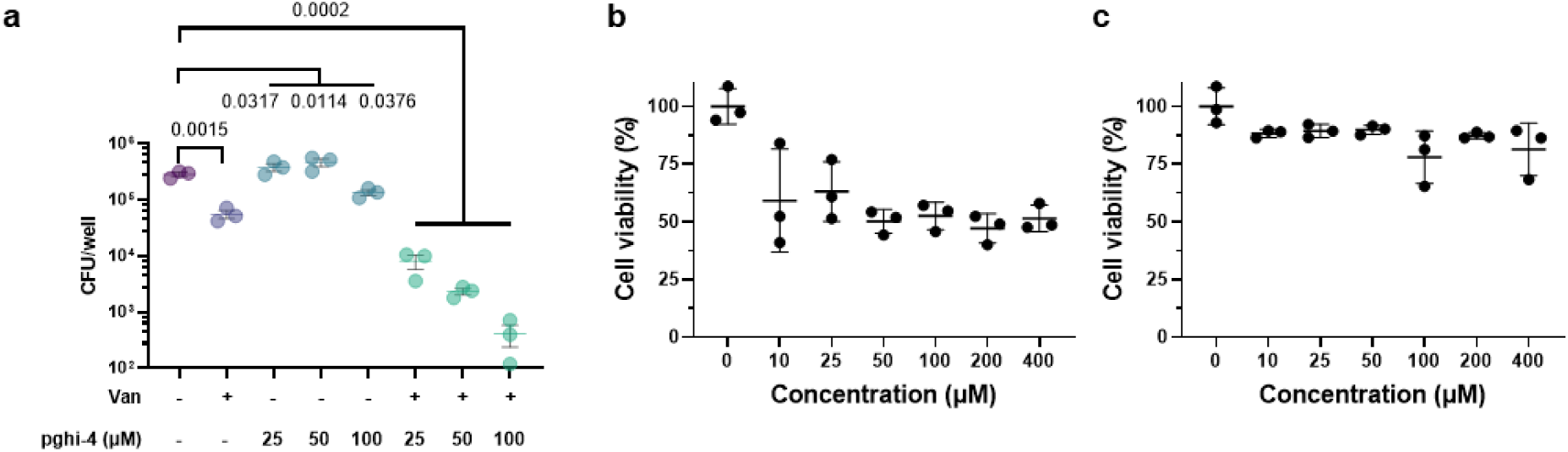
**a,** CFU analysis of VREfm-infected THP-1 cells treated with vancomycin (100 µg/mL), pghi-4 alone or in combination. Data are mean ± S.E.M. and analyzed by analyzed with one-way ANOVA and Tukey’s multiple comparison post-test, p values are indicated, n=3 biological replicates. **b,** Cytotoxicity of pghi-4 on RAW264.7 cells assessed by WST-8 assay. **c,** Cytotoxicity of pghi-4 on THP-1 cells assessed by WST-8 assay. For **b** and **c**, data are mean ± S.D., n=3 biological replicates.

**Extended Data Fig. 20.**
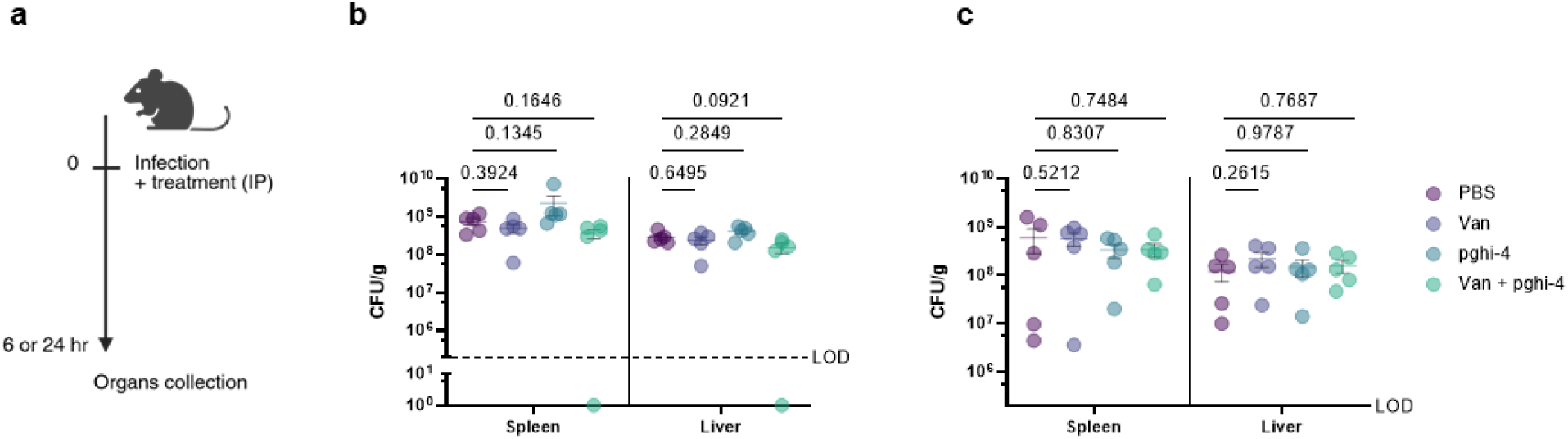
Single-dose therapeutic efficacy of pghi-4 as vancomycin adjuvant in vivo. **a,** Scheme of systemic infection *in vivo* with VREfm (ERV165): mice were infected with 10^9^ VREfm CFU with PBS (0.25% CMC), pghi-4 (25 mg/kg 0.25% CMC) ± vancomycin (Van, 100 mg/kg 0.25% CMC) intraperitoneally. 6- or 24 hours post-infection mice were sacrificed, organs were collected and VREfm burden was analyzed by CFU. **b,** CFU analysis of organs from VREfm-infected mice pghi-4 (25 mg/kg 0.25% CMC) ± vancomycin (Van, 100 mg/kg 0.25% CMC). collected after 6 hours post-infection. Dotted horizontal line is limit of detection (LOD). **c,** CFU analysis of organs from VREfm-infected mice ± pghi-4 (25 mg/kg 0.25% CMC) ± vancomycin (Van, 100 mg/kg 0.25% CMC) collected after 24 hours post-infection. Y axis starts with the limit of detection (LOD). For **b, c,** horizontal line is mean ± S.E.M. and analyzed by Kruskal-Wallis test with Dunn’s uncorrected post-test. Each dot is an individual mouse, n=5.

**Extended Data Fig. 21.**
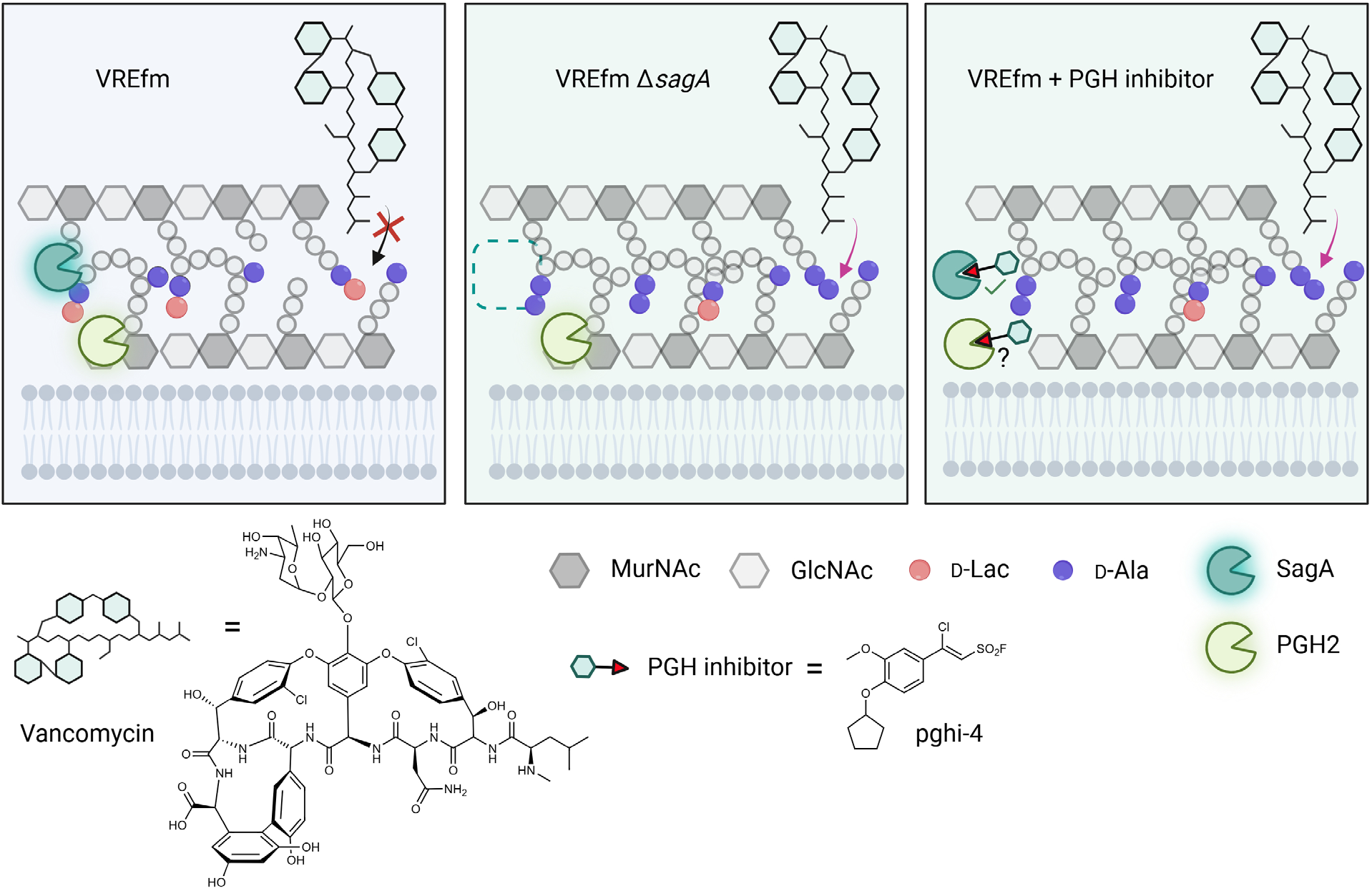
Summary of SagA inactivation and impact on vancomycin susceptibility in VREfm. VREfm with functional SagA peptidoglycan remodeling display D-Lac-ending peptidoglycan fragments enabling vancomycin binding. Genetic inactivation of SagA impairs peptidoglycan remodeling in VREfm, which results in increased vancomycin susceptibility. Pharmacological inactivation of SagA and perhaps PGH2 by PGH inhibitor (pghi-4) impairs peptidoglycan remodeling in VREfm, which results in increased accumulation of D-Ala-D-Ala-containing peptidoglycan and vancomycin susceptibility. Created with BioRender.

**Extended Data Fig. 22.**
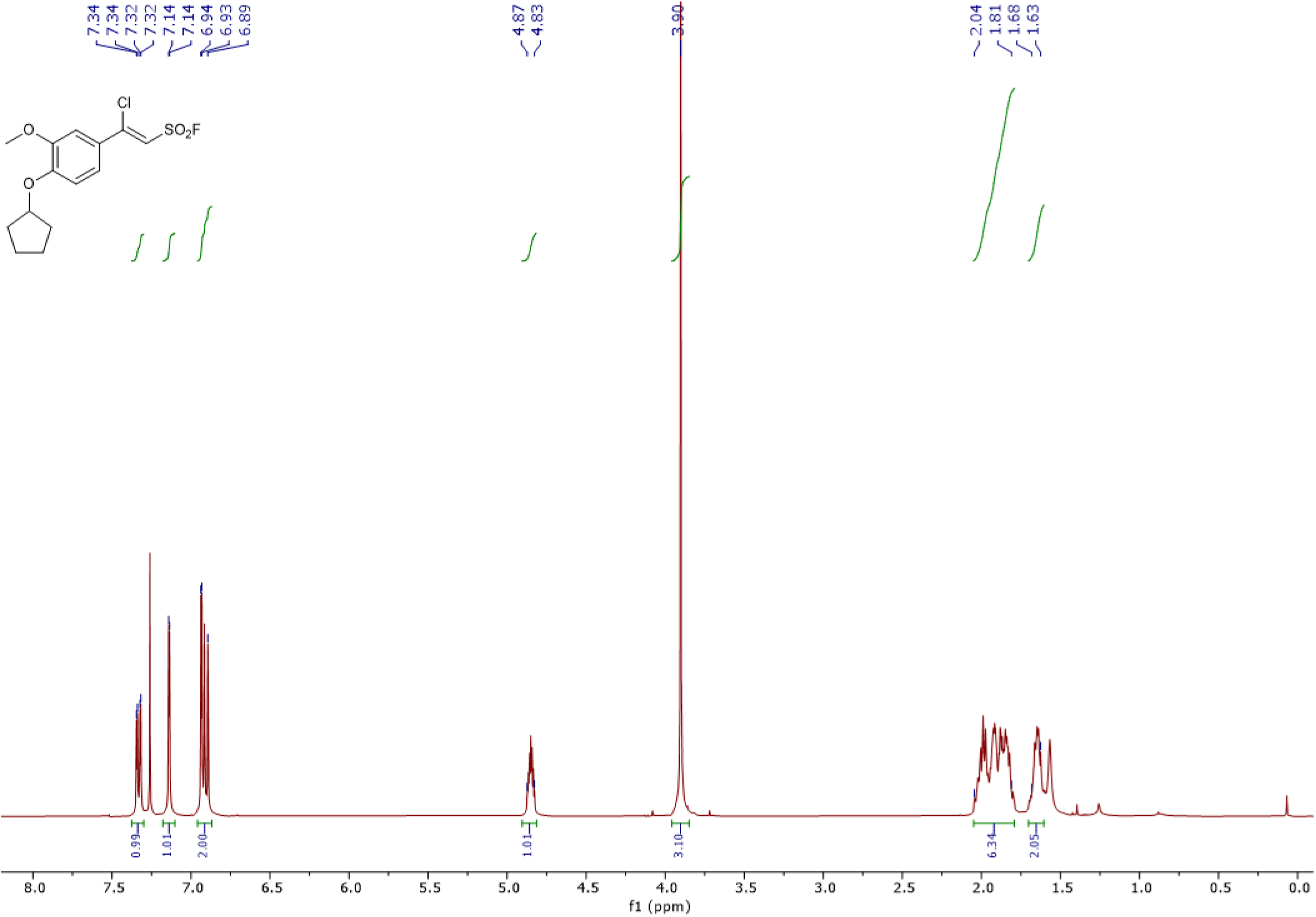
^1^H NMR Spectrum for pghi-4 (400 MHz, CDCl_3_)

**Extended Data Fig. 23.**
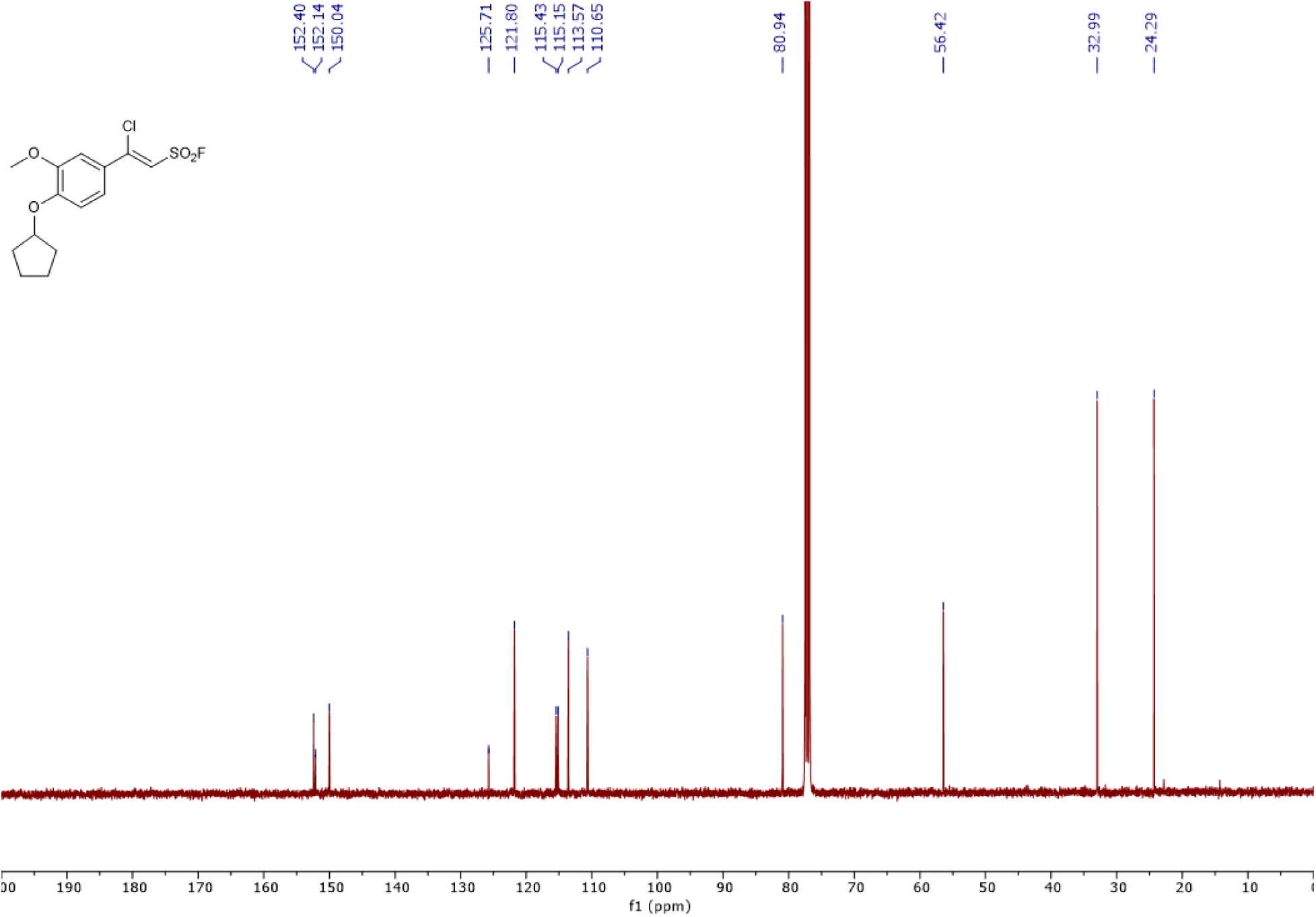
^13^C NMR Spectrum for pghi-4 (101 MHz, CDCl_3_)

**Extended Data Fig. 24.**
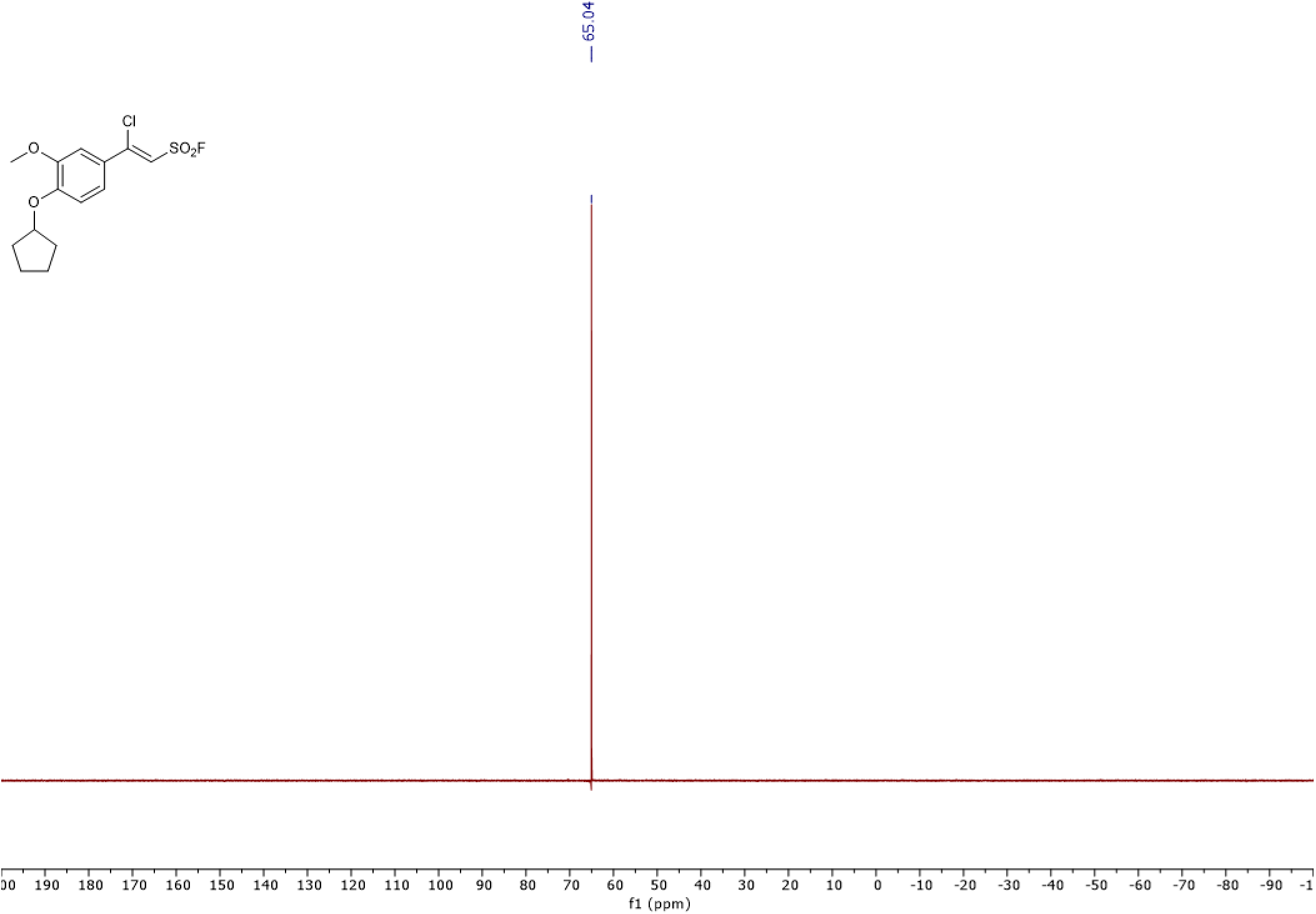
^19^F NMR Spectrum for pghi-4 (377 MHz, CDCl_3_)

**Extended Data Fig. 25.**
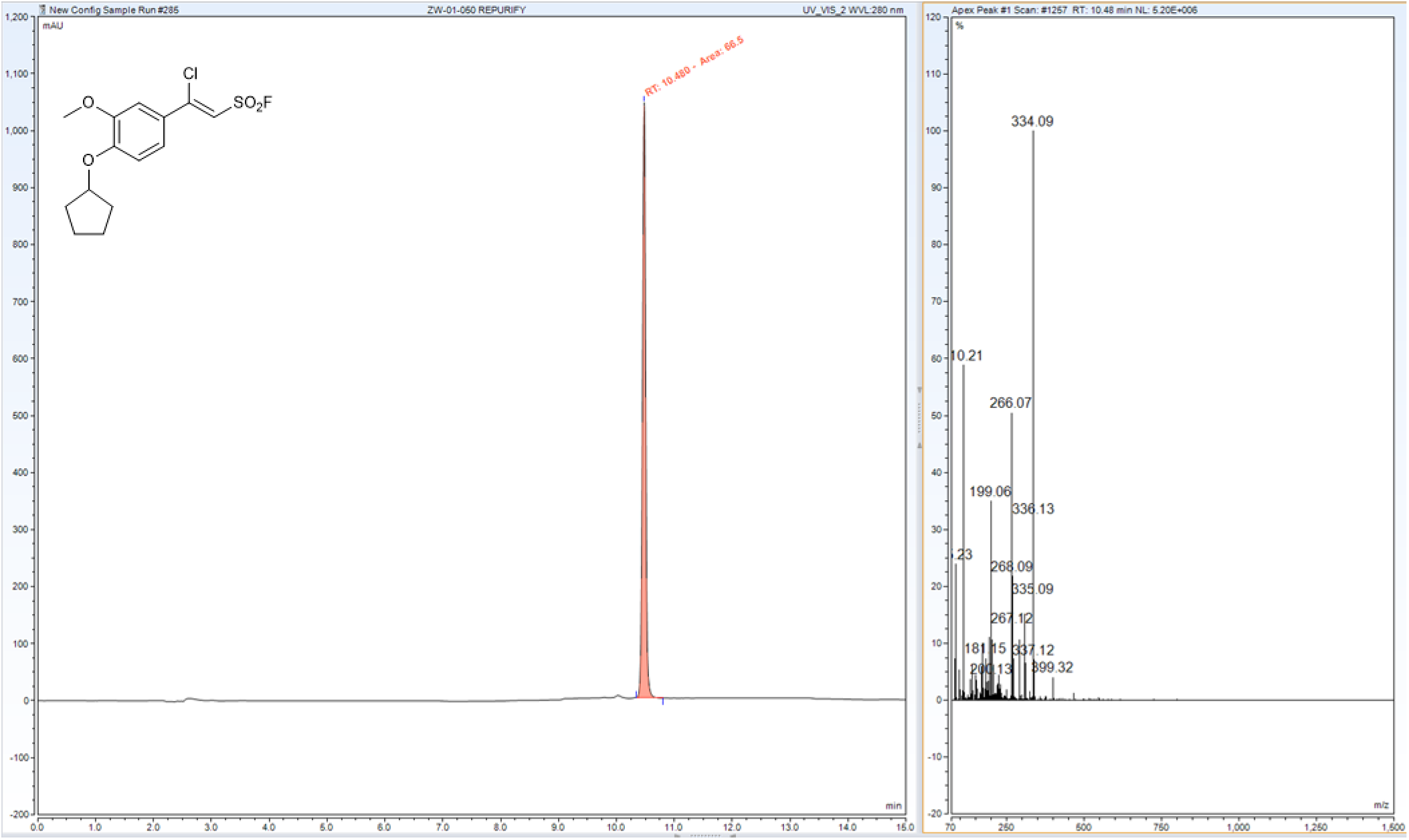
Liquid chromatography-mass spectrometry (LC-MS) analysis of pghi-4.

**Extended Data Table 1.** Minimum inhibitory concentration (MIC) values of VREfm susceptibility to ampicillin, daptomycin and ceftriaxone.

| **ERV165 strain** | **Ampicillin MIC**<br>**(µg/mL)** | **Daptomycin MIC**<br>**(µg/mL)** | **Ceftriaxone MIC**<br>**(µg/mL)** |
| --- | --- | --- | --- |
| WT | 8 | 128 | >128 |
| *ΔsagA* | 4 | 128 | >128 |
| *ΔsagA::sagA* | 8 | 128 | >128 |

**Extended Data Table 2.** Minimum inhibitory concentration (MIC) values of VREfm susceptibility to vancomycin.

| ERV165 strain | MIC in BHI<br>(µg/mL) | MIC in MHB<br>(µg/mL) |
| --- | --- | --- |
| WT | 256 | 256 |
| *ΔsagA* | 128 | 128 |
| *ΔsagA::sagA* | 256 | 256 |

**Extended Data Table 3.** Missense and frame-shift mutations detected by whole-genome sequencing (WGS) in ERV165-Δ*sagA*.

| № | Gene | Mutation | Annotation |
| --- | --- | --- | --- |
| 1 | *sagA* | Clean KO | NlpC/p60 PG hydrolase |
| 2 | EEHMGE_14260 | N233D | PTS mannose transporter subunit EIIAB |
| 3 | EEHMGE_11825 | R159H | DNA-binding protein |
| 4 | EEHMGE_08940 | N104D | glycosyl hydrolase |
| 5 | EEHMGE_08165 | T424N | glycine/betaine ABC transporter permease |
| 6 | EEHMGE_01745 | L10S, L68S | MapZ (Midcell Anchored Protein Z) locates at the division site before FtsZ and guides septum positioning. The mutation occurred outside annotated functional domains and in a region lacking conserved sequence features. |
| 7 | EEHMGE_15160 | L14S | AI-2E family transporter |
| 8 | EEHMGE_07965 | Frame-shift variant | Alpha-N-arabinofuranosidase |

**Extended Data Table 4.** Silent mutations detected by WGS in ERV165-Δ*sagA*.

| № | Gene | Mutation | Annotation |
| --- | --- | --- | --- |
| 1 | *neuC* (EEHMGE_14960) | I51I, L21L | UDP-N-acetylglucosamine 2-epimerase |
| 2 | EEHMGE_14675 | K25K | Terminase large subunit |
| 3 | *xseA* (EEHMGE_14305) | Q312Q | Exodeoxyribonuclease VII large subunit |
| 4 | EEHMGE_11955 | I74I | MarR family transcriptional regulator |
| 5 | EEHMGE_11645 | I71I | Universal stress protein |
| 6 | EEHMGE_01935 | S32S | DUF2140 domain-containing protein |
| 7 | EEHMGE_09405 | K167K | Insulinase family protein |
| 8 | EEHMGE_08170 | L116L | Glycine betaine/L-proline ABC transporter ATP-binding protein |
| 9 | EEHMGE_04975 | A162A | LTA synthase family protein |
| 10 | EEHMGE_04420 | V29V | Thioredoxin |
| 11 | EEHMGE_03730 | I213I | NAD(P)-dependent dehydrogenase |

**Extended Data Table 5.**
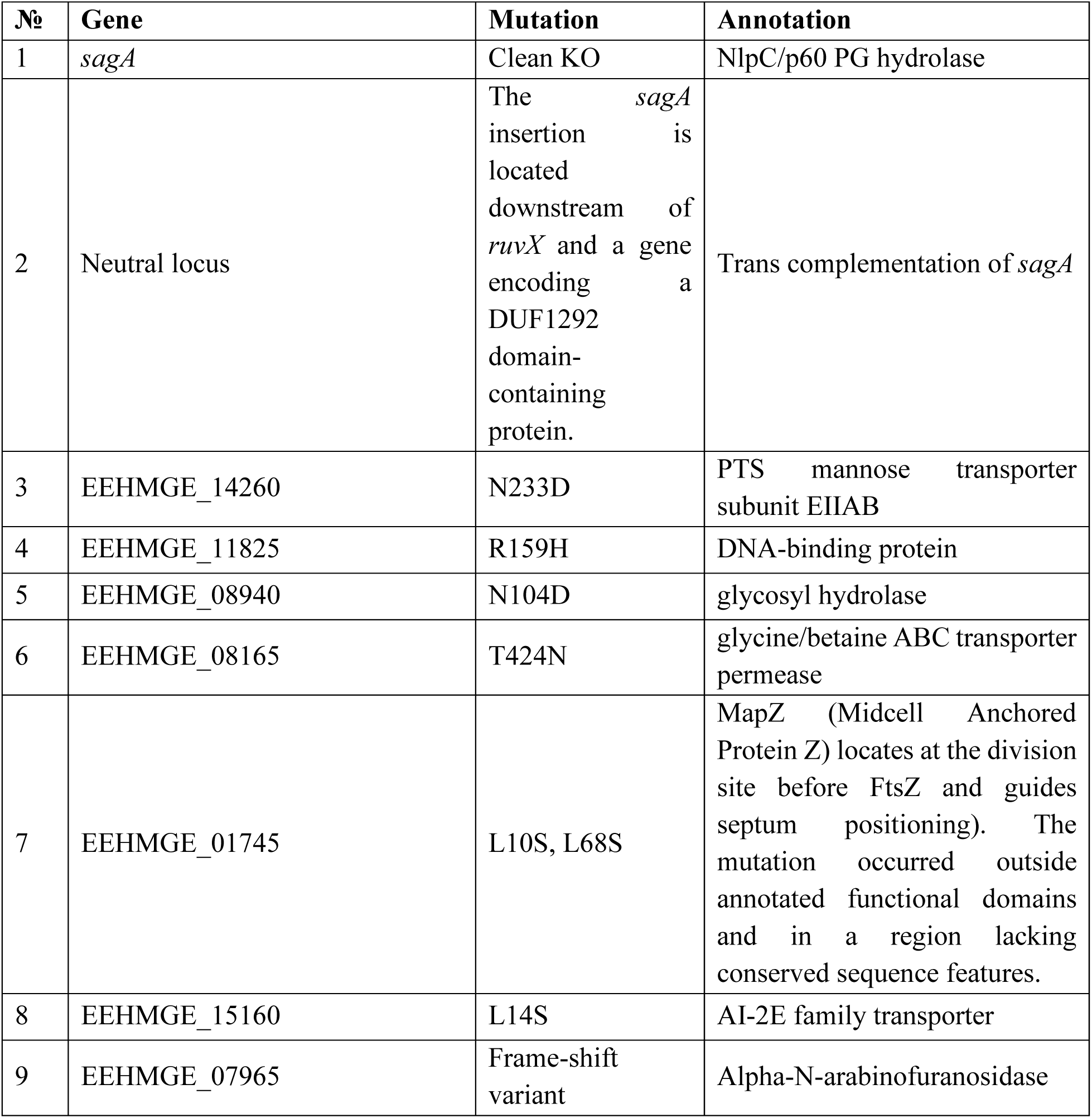
Missense and frame-shift mutations detected by WGS in ERV165-Δ*sagA*::*sagA*.

**Extended Data Table 6.** Silent mutations detected by whole-genome sequencing (WGS) in ERV165-Δ*sagA*::*sagA*.

| № | Gene | Mutation | Annotation |
| --- | --- | --- | --- |
| 1 | *neuC* (EEHMGE_14960) | I51I, L21L | UDP-N-acetylglucosamine 2-epimerase |
| 2 | EEHMGE_14675 | K25K | Terminase large subunit |
| 3 | *xseA* (EEHMGE_14305) | Q312Q | Exodeoxyribonuclease VII large subunit |
| 4 | EEHMGE_11955 | I74I | MarR family transcriptional regulator |
| 5 | EEHMGE_11645 | I71I | Universal stress protein |
| 6 | EEHMGE_09405 | K167K | Insulinase family protein |
| 7 | EEHMGE_08170 | L116L | Glycine betaine/L-proline ABC transporter ATP-binding protein |
| 8 | EEHMGE_04975 | A162A | LTA synthase family protein |
| 9 | EEHMGE_04420 | V29V | Thioredoxin |
| 10 | EEHMGE_03730 | I213I | NAD(P)-dependent dehydrogenase |
| 11 | EEHMGE_01935 | S32S | DUF2140 domain-containing protein |

**Extended Data Table 7.** Minimum inhibitory concentration (MIC) values of VREfm susceptibility to vancomycin in the presence of pghi-4 in BHI analyzed by checkerboard assay (Fig. 4b).

| **pghi-4 (μM)** | **MIC (μg/mL)** | **ΔMIC (fold)** |
| --- | --- | --- |
| 0 | 320 | 1 |
| 6.25 | 320 | 1 |
| 12.5 | 320 | 1 |
| 25 | 160 | 2 |
| 50 | 80 | 4 |
| 100 | 40 | 8 |

**Extended Data Table 8.** VREfm strain information.

| **VREfm strain** | **Sequence Type** | **Van Type** |
| --- | --- | --- |
| ERV165 | 412 | *vanA* |
| TX0082 | 17 | *vanA* |
| 700221 | 17 | *vanA* |
| DVT 1238 | 412 | *vanA* |
| DVT 1574 | 80 | *vanA* |
| DVT 1576 | 117 | *vanA* |
| DVT 3347 | 18 | *vanA* |
| DVT 3350 | 203 | *vanA* |
| DVT 3351 | 664 | *vanA* |

**Extended Data Table 9.**
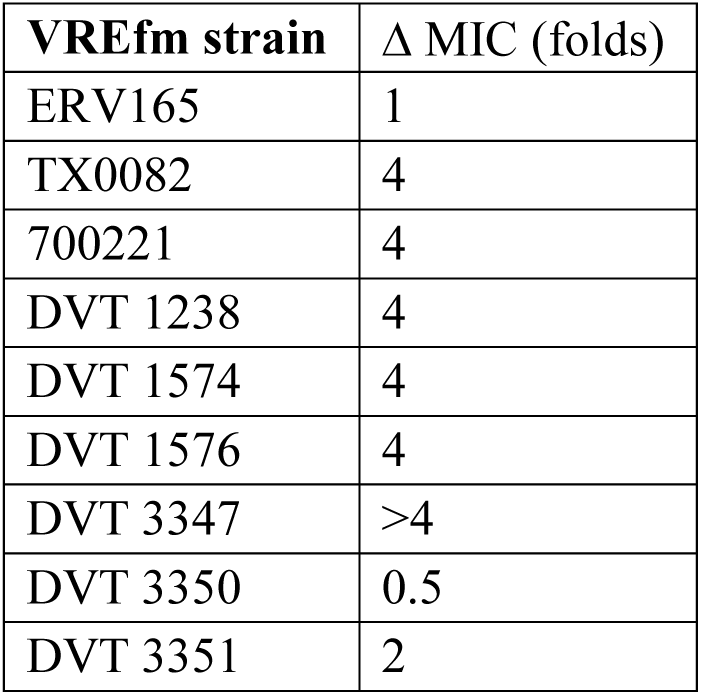
Minimum inhibitory concentration (ΔMIC) values of vancomycin susceptibility of VREfm clinical isolates relative to ERV165.

**Extended Data Table 10.** Bacterial strains used in this study.

| Organism | Description | Source |
| --- | --- | --- |
| *Enterococcus faecium* ERV165 | WT | Lab stock |
| *Enterococcus faecium* ERV165-Δ*sagA* | Δ*sagA* (clean knock-out) | This work |
| *Enterococcus faecium* ERV165-Δ*sagA*::*sagA* | Δ*sagA*::*sagA* (*sagA* chromosomal complementation in Δ*sagA* background) | This work |
| *Enterococcus faecium* ERV165-Δ*pgh2* | Δ*pgh2* (clean knock-out) | This work |
| *Enterococcus faecium* ERV165-Δ*pgh3* | Δ*pgh3* (clean knock-out) | This work |
| *Enterococcus faecium* ERV165-Δ*pgh4* | Δ*pgh4* (clean knock-out) | This work |
| *Enterococcus faecium* TX0082 | WT | Lab stock |
| *Enterococcus faecium* 700221 | WT | Lab stock |
| *Enterococcus faecium* DVT 1238 | Clinical isolate | Van Tyne lab |
| *Enterococcus faecium* DVT 1574 | Clinical isolate | Van Tyne lab |
| *Enterococcus faecium* DVT 1576 | Clinical isolate | Van Tyne lab |
| *Enterococcus faecium* DVT 3347 | Clinical isolate | Van Tyne lab |
| *Enterococcus faecium* DVT 3350 | Clinical isolate | Van Tyne lab |
| *Enterococcus faecium* DVT 3351 | Clinical isolate | Van Tyne lab |
| *Escherichia coli* BL21-CodonPlus (DE3)-RIL<sup>1</sup> | *Efm_Com15_SagA-NlpC/p60_ΔSS-His<sub>6</sub>* | Lab stock |
| *Escherichia coli* BL21-CodonPlus (DE3)-RIL<sup>1</sup> | *Efm_Com15_SagA-NlpC/p60_ΔSS_C443A-His<sub>6</sub>* | Lab stock |

<sup>1</sup> *Escherichia coli* BL21-CodonPlus (DE3)-RIL cells contain extra copies of the *argU*, *ileY*, and *leuW* tRNA genes in a plasmid encoding Cam resistance

**Extended Data Table 11.** Plasmids constructed. The details of the plasmid construction are described below.

| № | Plasmid | Description | Source | Notes |
| --- | --- | --- | --- | --- |
| 1. | pJC005.gent<sup>1</sup> | CRISPR-Cas12a genome editing plasmid | Chua et al. <sup>29</sup> | Gent<sup>R</sup> |
| 2. | pUCsRNAP<sup>2</sup> | pUC; small RNA promoter and 23-bp spacer sequence flanked by two 19-bp repeats | Chua et al. <sup>29</sup> | Amp<sup>R</sup> |
| 3. | pPK126 | pLZ12-empty vector | This work | Spec<sup>R</sup> |
| 4. | pPK111 | pLZ12-*sagA*<sup>WT</sup> | This work | Spec<sup>R</sup> |
| 5. | pPK125 | pLZ12-*sagA*<sup>C425A</sup> | This work | Spec<sup>R</sup> |
| 6. | pPK99 | pJC005.gent-Δ*sagA* | This work | Gent<sup>R</sup>. This plasmid is used for constructing Δ*sagA* using the CRISPR-Cas12a counter-selection recombineering method described in Chua et al. <sup>29</sup> |
| 7. | pPK158 | pJC005.gent-*sagA* | This work | Gent<sup>R</sup>. This plasmid carries ERV165 *sagA* for chromosomal integration at a neutral locus in sPK377 using the CRISPR-Cas12a counter-selection recombineering method described in Chua et al. <sup>29</sup> |
| 8. | pPK156 | pJC005.gent- Δ*pgh2* | This work | Gent<sup>R</sup>. This plasmid is used for constructing Δ*pgh2* using the CRISPR-Cas12a counter-selection recombineering method described in Chua et al. <sup>29</sup> |
| 9. | pPK107 | pJC005.gent- Δ*pgh3* | This work | Gent<sup>R</sup>. This plasmid is used for constructing Δ*pgh3* using the CRISPR-Cas12a counter-selection recombineering method described in Chua et al. <sup>29</sup> |
| 10. | pPK154 | pJC005.gent- $\Delta pgh4$ | This work | Gent<sup>R</sup>. This plasmid is used for constructing $\Delta pgh4$ using the CRISPR-Cas12a counter-selection recombineering method described in Chua et al.<sup>29</sup> |

<sup>1</sup>Gift from James Collins (Addgene plasmid # 182739; http://n2t.net/addgene:182739; RRID: Addgene\_182739).
<sup>2</sup>Gift from James Collins (Addgene plasmid # 182746; http://n2t.net/addgene:182746; RRID: Addgene\_182746)

**Extended Data Table 12.** Masses of peptidoglycan fragments were detected with MSD API-ES.

| Peak<sup>a</sup> | RT (min) | Calculated molecular weight | Observed mass | Proposed structure<sup>b</sup> |
| --- | --- | --- | --- | --- |
| 1 | 5.01 | 825.4 | 413.6 [M+2H]<sup>2+</sup> | GM-tri |
| 2 | 6.77 | 697.3 | 349.6 [M+2H]<sup>2+</sup> | GMDP |
| 3 | 9.63 | 896.44 | 449.2 [M+2H]<sup>2+</sup> | GM-tetra |
| 4 | 12.58 | 939.45 | 470.8 [M+2H]<sup>2+</sup> | GM-tri (Asn)<sup>c</sup> |
| 5 | 15.04 | 940.43 | 471.2 [M+2H]<sup>2+</sup> | GM-tri (Asp)<sup>c</sup> |
| 6 | 18.02 | 1010.5 | 506.2 [M+2H]<sup>2+</sup> | GM-tetra (Asn) |
| 7 | 19.33 | 1081.5 | 541.6 [M+2H]<sup>2+</sup> | GM-penta (Asx) |
| 8 | 31.54 | 1818.9 | 909.8 [M+2H]<sup>2+</sup> | 2GM-tri (Asx) - tetra |
| 9 | 35.47 | 1932.9 | 966.6 [M+2H]<sup>2+</sup> | 2GM-tri (Asn) - tetra (Asn)<sup>c</sup> |
| 10 | 37.22 | 1933.9 | 967.4 [M+2H]<sup>2+</sup> | 2GM-tri (Asx) - tetra (Asx)<sup>c</sup> |
| 11 | 39.93 | 2004.0 | 1002.4 [M+2H]<sup>2+</sup> | 2GM-tetra (Asn) - tetra (Asn)<sup>c</sup> |
| 12 | 41.57 | 2005.9 | 1004.2 [M+2H]<sup>2+</sup> | 2GM-tetra (Asp) - tetra (Asp)<sup>c</sup> |
| 13 | 47.72 | 2927.4 | 975.8 [M+3H]<sup>3+</sup> | 3GM-tetra (Asx) - tetra (Asx) - tri (Asx)<sup>c</sup> |
| 14 | 49.02 | 2928.4 | 977.0 [M+3H]<sup>3+</sup> | 3GM-tetra (Asx) - tetra (Asx) - tri (Asx)<sup>c</sup> |
| 15 | 50.94 | 2999.4 | 1001.4 [M+3H]<sup>3+</sup> | 3GM-tetra (Asx) - tetra (Asx) - tetra (Asx)<sup>c</sup> |

<sup>a</sup>. Peak numbers refer to Extended Data Fig. 4a and 15a.
<sup>b</sup>. GM, disaccharide (GlcNAc-MurNAc); 2GM, disaccharide-disaccharide (GlcNAc-MurNAc-GlcNAc-MurNAc); 3GM, disaccharide-disaccharide-disaccharide (GlcNAc-MurNAc-GlcNAc-MurNAc-GlcNAc-MurNAc); GM-Tri, disaccharide tripeptide (L-Ala-D-iGln-L-Lys); GM-Tetra, disaccharide tetrapeptide (L-Ala-D-iGln-L-Lys-D-Ala); GM-Penta, disaccharide pentapeptide (L-Ala-D-iGln-L-Lys-D-Ala-D-Ala).
<sup>c</sup>. The assignment of the amide and the hydroxyl functions to either peptide stem is arbitrary.
RT = retention time

